# Commercial *Saccharomyces cerevisiae* baker’s yeasts: strain redundancy, genome plasticity, and colonization of the sourdough environment and the human body

**DOI:** 10.1101/2025.02.07.637159

**Authors:** Hanna Viktória Rácz, Alexandra Imre, Bálint Németh, Zsuzsa Antunovics, Rizagul Bazenova, Devin Bendixsen, Andrea Harmath, Lilla Herman, Ádám Fülep, Ksenija Lopandic, Endre Máthé, Szilárd Póliska, István Pócsi, Rike Stelkens, Walter P. Pfliegler

## Abstract

Leavening dough is one of the most widespread applications of fermentative yeast and the most common practice for the general public to come into contact with microbial cultures. *Saccharomyces cerevisiae* is the typical species used for dough making, but the evolutionary origin of strains isolated from dough is mixed. Here, using 49 newly sequenced and 183 previously described isolates from the bakery environment, we show that the traditional strains used in Europe for sourdough making are more closely related to Chinese Mantou sourdough lineages than to commercially used baker yeast strains. Surprisingly, the expansion of these traditional European strains into other human-associated niches, including the human body, has been very limited. This is in stark contrast to the Mixed-origin clade commercial baking yeasts, which consists of only a few globally distributed clonal lineages that dominate the yeast market and recurrently colonize sourdoughs and human hosts. These clonal lineages consistently maintain their ploidy, unique heterozygous chromosomal rearrangements, and stable aneuploidies in addition to several diverse structural variants. In addition to the previously known diploid and tetraploid groups of commercial isolates, we describe a widely distributed stable triploid aneuploid clonal lineage. We show that company practices and global trade help the distribution of these clonal clusters and that these yeasts are characterized by their exclusively mitotic reproduction.

## Introduction

### Diversity, origin, and genomic features of the Mixed origin clade and commercial baker’s yeasts

Our knowledge on the evolutionary and domestication history of the diverse clades of the yeast *Saccharomyces cerevisiae* has significantly expanded in recent years thanks to large-scale population genomics studies (e.g. Strope *et al*. 2015; Li *et al*. 2016; Peter *et al*. 2018; Legras *et al*. 2018; Duan *et al*. 2018; Loegler *et al*. 2024). This work has focused on the global diversity of this yeast species in natural environments and the domesticated clades and mosaic (namely Mosaic region 1, 2, and 3) groups associated with human-made fermentation habitats (Almeida *et al*. 2015; Peter *et al*. 2018; Legras *et al*. 2018; Pontes *et al*. 2020; Han *et al*. 2021; Lee *et al*. 2022). Other studies focus on the genomic analysis of strains found in ale (Gallone *et al*. 2016, 2018; Gonçalves *et al*. 2016; Preiss *et al*. 2018, 2024; Fay *et al*. 2019; Saada *et al*. 2022), alpechin (Pontes *et al*. 2019), wine (Borneman *et al*. 2016; Gayevskiy *et al*. 2016; Coi *et al*. 2017; Higgins *et al*. 2021; Basile *et al*. 2021; Alexander Marr *et al*. 2023; Ward *et al*. 2024), and clinical isolates (Strope *et al*. 2015; Zhu *et al*. 2016; Ramazzotti *et al*. 2019; Morard *et al*. 2023). However, the genomic analysis of strains used in baking has gathered less attention, although leavening bread and other products is probably the most ubiquitous use of *S. cerevisiae* by the general public (e.g. Lahue *et al*. 2020). Usually, only a limited number of baking strains in the form of sourdough isolates or commercially available active dry or pressed baker’s yeast is included in phylogenetic studies. What is currently lacking is a comprehensive and comparative analysis of *S. cerevisiae* strains isolated from baking environments that takes the recently published collections of genomes into account all together.

So far, most sequenced bakery strains were shown to cluster together in a clade named ‘Mixed origin’ (Gallone *et al*. 2016; Peter *et al*. 2018), characterized by no clear ecological or geographic enrichment – hence the name. Yeasts in this clade were isolated mostly from bakeries, baker’s yeast products, clinical samples, ale beers, and from nature (plants and water bodies), from Europe, Eastern and Western Asia, Africa, or South America. The clade was found to share some of its polymorphisms with the Ale 1 (Ale beer) clade, but is clearly distinct from it (Peter *et al*. 2018; Pontes *et al*. 2020). In their analysis, Duan et al. (2018) focused on yeast from China and identified a Chinese Active Dry Yeast (ADY) clade that was later shown to be identical to the Mixed origin (Han *et al*. 2021). Importantly, the analysis of Chinese mantou (traditional steamed bun made from fermented dough) isolates also led to the identification of divergent local baking clades named Mantou 1 to 7 that clustered into the Solid State Fermentation superclade along with the newly described Baijiu, Huangjiu, and the previously established Sake and Asian Fermentation clades (Duan *et al*. 2018; Han *et al*. 2021). The authors also delineated a Mantou Mosaic group containing several Chinese dough isolates. However, at the time of these publications, sequenced bakery isolates from other parts of the world were scarce, preventing more detailed comparisons.

### True sourdough yeasts compared to baker’s yeasts in sourdough

A recent study by Bigey et al. (2021) clearly identified two distinct domestication trajectories and five distinct main populations of baker’s and sourdough yeasts by comparing 51 previously sequenced and 17 new genomes. While the Mixed origin clade was shown to contain most of the commercial bakery yeasts in two sub-populations, yeasts isolated from traditional sourdoughs from France, Belgium, the Italian mainland, Sicily, and Turkey clustered mostly into the Mosaic region 3, the Asian fermentation clade, the Wine/European clade, and the Mixed origin clade. In the latter, sourdough isolates clustered with commercial strains, suggesting colonization of sourdough by commercial bakery strains. The samples from maize dough from Ghana were found to be unrelated to these, and belonged African beer clade (Bigey *et al*. 2021). The authors also concluded that the vast majority of commercial baker’s strains in the Mixed origin clade are tetraploid or close-to-tetraploid aneuploids, in line with other observations on a smaller collection of genomes (Zhu *et al*. 2016; Peter *et al*. 2018; Rácz *et al*. 2021). These yeasts showed shorter fermentation latency, a hallmark of artificial selection to the industrial environment. The true sourdough yeasts populations, however, showed adaptations to the artisanal bakery processes, e.g. increased copy number of the maltose and isomaltose permeases and maltase/isomaltase transcription regulators, which leads to increased growth on maltose and in sourdough (Bigey *et al*. 2021). Most sourdough-derived isolates that were not related to the Mixed origin clade of commercial strains were diploids, clearly differentiating these from the commercial yeasts. However, comparisons to the several described Mantou yeast clades were not conducted, and their potential relatedness was not investigated. As more and more knowledge started to accumulate on traditional sourdoughs, it is furthermore noteworthy that novel types of yeasts unrelated to the above-mentioned clades have also been tested in sourdough-making. This includes the *S. cerevisiae* var. *‘boulardii’* probiotic strains, paving the way for additional lineages to colonize the sourdough environment (Koj and Pejcz 2023).

### Clinical isolates of the Mixed origin clade and the M3 Mosaic region

Studying baker’s and sourdough yeasts may also be important in understanding how members of the Mixed origin clade and the M3 Mosaic region successfully colonize the human gastrointestinal tract and in rare cases, cause opportunistic infections. Several studies have linked the usually polyploid commercial baker’s yeasts of the Mixed origin clade to clinical isolates (de Llanos *et al*. 2006; Zhu *et al*. 2016; Pfliegler *et al*. 2017; Morard *et al*. 2023). Whether diploid sourdough yeasts are related to clinical isolates has not been evaluated so far, but the M3 Mosaic region has many described clinical isolates with several sourdough yeasts also clustering with this mosaic region (Bigey *et al*. 2021). This raises the question whether true sourdough yeasts may also be colonizers of the human host.

### Yeast manufacturers

An often-neglected aspect in the study of yeast diversity and phylogenomics is the role that the commercial manufacturing and distribution of yeast strains for bakeries, breweries, wineries, and for the public, may play in shaping the biogeography and ecology of the species. Yeast manufacturing is a multinational industry that distributes strains worldwide. This can lead to the repurposing of yeast strains originally adapted to one fermentation niche into a new fermentation environment. Yet, manufacturer information is usually absent from most studies (but see Borneman *et al*. 2016), or not used to add context to the results about the genetic relationships of strains and their biogeography. Modern yeast manufacturing of pure baking cultures was established only relatively recently in the 1920s (Frey 1930; Lahue *et al*. 2020), while for millennia before that people relied on spontaneous fermentations or used and exchanged traditional microbial starter cultures. Ever since large-scale manufacturing started, companies have been established, merged with each other, or gone out of business; accompanied by a constant change in brand names. Considering the exact origin of commercial baker’s yeasts may thus add important context to comparative analyses, especially where various brand names and samples may all be linked to a single, globally present manufacturer. For instance, a recent publication on commercial wine yeast strains (Borneman *et al*. 2016) used information on nine different producer companies and found a high level of strain redundancy, i.e. many identical or nearly identical genomes across products and isolates from wineries. Across these samples, genetic variation was often so minor that it likely accumulated after long-term passaging from the same source at various producers and wineries (Borneman *et al*. 2016). Industrial strains are, by definition, genetically uniform. However, passaging, large-scale propagation during production, along with serial repitching in some industries (common practice in breweries or in sourdough) inevitably leads to mutation accumulation, manifesting in clonal heterogeneity and the emergence of subclone lineages. Among the accumulated mutations, structural chromosomal changes are especially prevalent in aneuploids strains (Large *et al*. 2020; Rácz *et al*. 2021).

### Sourdough and baker’s yeasts in Hungary and the rationale of our study

In our study, our aims were to (1) systematically survey yeast diversity in the bakery environment using an extensive collection of yeasts that covered the majority of commercially available products in Hungary; (2) to determine the strain redundancy in baker’s yeast products and the level of genomic clonal heterogeneity; (3) and to compare baking and sourdough isolates to the numerous clinical samples from across the world in the Mixed origin clade and the Mosaic 3 region. Our analysis of a substantial collection of samples from a well-defined niche, namely the bakery-related fermentation environment, contributes to a better understanding of the origin, genomic features, strain redundancy, and niche expansion of the Mixed origin and Mosaic 3 yeasts.

## Materials and Methods

### New commercial baker’s yeast and sourdough samples

We built a collection of commercially available baker’s yeast products by purchasing two separate batches of altogether 16 different products from different retailers. The first batches of each product were obtained in a period between September 2017 and November 2017, the second batches between March 2018 and May 2018. Formulation, manufacturer, and country of origin were noted. Manufacturers and brands were given individual identifiers. Additionally, a bread mix containing active dry yeast and a self-rising refrigerated dough were also obtained for yeast isolation. Yeast products were rehydrated and/or suspended in YPD (yeast extract, peptone, dextrose, VWR Chemicals, Solon, OH, USA) and spread to YPD agar plates and incubated for 2–3 days at 22°C. A single colony isolate was obtained from each batch of each product. Sourdoughs were obtained from a bakery in Balmazújváros, Eastern Hungary that maintains spontaneous wheat and rye sourdoughs of type 1 (as defined by Hammes *et al*. 2005) and backslopped sourdoughs with starters (type 3). As rural, local spontaneous sourdoughs are thought to be extinct in the country (see Discussion), we also obtained a spontaneous type 1 sourdough from an ethnic Hungarian village community in Transylvania, Romania. Yeast with different morphologies from sourdoughs were isolated after spreading to DRBC (Dichloran Rose Bengal Chloramphenicol, VWR Chemicals) agar (incubation at 2–3 days at 22°C). Isolates were deposited into our department’s culture collection at –70°C in YPD medium supplemented with 30% v/v glycerol.

### Yeast identification

Colony DNA for PCR tests was isolated according to Lõoke et al. (2011) from single-cell colonies and stored in 1×TE. These colony DNA samples were used to differentiate *Saccharomyces* samples from other yeasts. We used the interdelta and microsatellite fingerprinting multiplex PCR combining interdelta, microsatellite (*YLR177w*, *YOR267c*), and as a control, ITS 1–4 primer pairs (Imre *et al*. 2019). After gel electrophoresis (2% TBE agarose, 90 min, 100 V) we identified *S. cerevisiae* samples and whole genome sequenced them. To provide details on the sourdough mycobiome, non-*Saccharomyces* yeasts from sourdoughs were also identified to the species level. This was achieved by Sanger sequencing as described in Supplementary File S1. A list of isolates and genomes is provided in Supplementary Tables S1–S2.

### Bacterial long-read 16S metabarcoding of sourdough samples

To provide ecological context to the *Saccharomyces* isolates from sourdoughs, we used metabarcoding with the 16S long-read metabarcoding kit (SQK-16S024) of Oxford Nanopore Technologies (Oxford, UK) according to the manufacturer’s instructions, and analyzed the data using Emu (Curry *et al*. 2022) as described in Supplementary File S1.

### Companies and brands

Manufacturers and brands of newly isolated yeasts and those of previously sequenced genomes (where known) were evaluated extensively using public data on companies, trademark owners, manufacturing plants, and company structure (parent companies, recent mergers) using Google internet searches (last accessed: September 2024). Brand names and company names were both included in the searches and a list was compiled from the results (Supplementary Table S2). Individual identifiers were given to the companies.

### Karyotyping of *Saccharomyces* isolates

Karyotyping was performed using 1 % agarose gel (chromosomal grade, Bio-Rad, Hercules, CA, USA) by a counter-clamped homogenous electric field electrophoresis device (CHEF-Mapper; Bio-Rad). The following running parameters were used: run time 26 h, voltage 6 V/cm, angle 120°, temperature 14 °C, and pulse parameters 60 to 120 s. As a control, a haploid *S. cerevisiae* control was used. After electrophoresis gels were stained with ethidium bromide and washed in sterile water for 48 h before photographing using UV-transillumination. Gel images were then compared for supernumerary chromosome bands (compared to the control) and variable bands in each clade. Karyotyping was conducted only with 41 local isolates from our culture collection.

### Sporulation tests, tetrad analysis, and isolation of spore clone cultures

To sporulate baker’s yeasts, potassium acetate agar was used (1% potassium acetate, 0.1% yeast extract, 0.05% glucose, 2% agar). Cultures were incubated on these plates for 10 days at 22°C, sporulation was checked under 400× magnification in a light microscope, and sporulation efficiency was calculated after counting 300 sporulating and/or non-sporulating cells. The Zeiss Axioskop fixed-stage microscope modified for tetrad dissection was used to isolate spores from at least ten asci, number of spores in each ascus was recorded, spore clone colonies were observed and isolated after 3–6 days of incubation and saved to our culture collection in glycerol stocks as above with identifiers referring to yeast isolate, tetrad, and spore. The number of dissected asci forming 0, 1, 2, 3, or 4 colonies was recorded.

### Inclusion of previously sequenced genomes in genomic analyses

To represent the highest possible diversity of baker’s and sourdough (incl. Chinese Mantou clades) yeasts, we surveyed a wide range of previous publications on yeast phylogenomics. For each yeast, collection data, formulation, and manufacturer were also determined wherever possible. In several cases, we could determine whether a strain was a spore clone using the methods section of the publication and the name of the strain (that contained plausible tetrad and spore identification numbers). We also included all human isolates we found in previous publications belonging to the Mixed origin clade and to the Mosaic region 3 described by Peter et al. (2018). Finally, representative genomes from the clades described by Peter et al. (2018) and Duan et al. (2018) were included in the analyses to cover the phylogenomic diversity of the species. Genomes used in this study are listed in Supplementary Table S1. Details on members of the Mixed origin clade and Mosaic 3 group and other baker’s yeasts are listed in Supplementary Table S2.

### Whole genome sequencing and assembly of *S. cerevisiae* isolates

Genomic DNA was isolated after 24 h growth of freshly revived stocks streaked onto YPD and incubated at 30°C. For Illumina sequencing, DNA isolation followed Hanna and Xiao (Hanna and Xiao 2006). Library preparation was performed using tagmentation with the Nextera DNA Flex Library Prep kit (Illumina, San Diego, CA, USA) according to the manufacturer’s protocol. Sequencing was performed using 150 bp paired-end reads on an Illumina NextSeq 500 system, with approximately 50–80× coverage of the nuclear genome (two genomes were sequenced with ∼400× coverage). Raw reads of new samples were deposited to NCBI SRA under BioProject PRJNA1218442 (reads of BY0012 were deposited under the already existing BioProject PRJNA646688). The Illumina FASTQ sequencing files were trimmed and filtered using fastp for further analysis (Chen *et al*. 2018).

For Oxford Nanopore sequencing of a baker’s strain UDeb-BY0012, genomic high molecular weight DNA was isolated with the Quick-DNA™ HMW MagBead Kit (Zymo Research, Irvine, CA, USA) according to the company’s instructions, except that cell lysis was not performed with lysozyme, but with R-Zymolyase (RNase-A containing zymolase; Zymo Research). Library construction was performed with the Ligation Sequencing Kit (SQK LSK109; Oxford Nanopore) and sequenced on a MinION R9.4.1 Flow Cell with approximately 300× coverage calculated for the S288c genome. Basecalling (super accurate model) of the resulting FAST5 sequencing files was performed with Guppy v6.4 software provided by the company, reads were uploaded to NCBI SRA (SRX27544410). The genome was assembled by using the obtained FASTQ files with the LRSDAY v1.6 pipeline (Yue and Liti 2018) combined with the following softwares: long reads were first adapter trimmed, filtered and downsampled to 60× coverage, assembly (Flye v2.9) (Freire *et al*. 2022) was polished with long reads (with 3× Racon v1.4, 3× Medaka v1.4) (Vaser *et al*. 2017) and reads from the Illumina sequencing (1× with POLCA) (Zimin and Salzberg 2020). The LRSDAY pipeline was also used to annotate the genome. The assembly was deposited to GenBank, accession number SUB15072669. The assembly was aligned against the S288c reference genome in LRSDAY. As the assembly was highly syntenic with the S288c reference and did not capture heterozygous structural variations, we used the S288c reference in the following analyses except for polyploid phasing. The software nPhase (Abou Saada *et al*. 2021) was applied for polyploid phasing using both long-and short-read data as described in Supplementary File S1.

### Comparative and phylogenomic analysis

First, we mapped Illumina reads to a reference panel of eight *Saccharomyces* species (downloaded from GenBank) to test whether isolates are hybrids or contain introgressed regions, and to run a phylogenomic analysis focusing on the nuclear genome of *S. cerevisiae*. Hybrid identification was based on visualizing read coverage across the eight species. As the reference involved the other species of the genus, this mapping approach allowed us to only compare the *cerevisiae* subgenomes of hybrids and to exclude introgressed regions. Second, to properly assess chromosome-wide read coverage, we also mapped solely to the S288c reference genome (or in the case of hybrids, to a composite reference of two species). See also Supplementary file S1 for examples on the use of the dual mapping approach.

### Mapping

Mapping was performed using the mem option of BWA 0.7.17. (Li and Durbin 2009). Sorted BAM files were obtained using Samtools 1.7. (Li *et al*. 2009) and Picard-tools 2.23.8. was used to mark duplicated reads (Van der Auwera *et al*. 2013). We used BEDTools 2.30.0 (Quinlan and Hall 2010) to create per-base BedGraph files and then to calculate the median coverage of chromosomes in 10000 base windows sliding every 5000 bases. Plots generated from this data were corrected for ploidy and were used to identify potential segmental duplications, deletions, or aneuploidies (see also Supplementary file S1).

### Variant calling

Using BAM files, local realignment around indels and joint variant calling and filtering for the strains and isolates were performed with GATK 4.1.9.0. [Poplin et al. 2018; Van der Auwera et al. 2013] with regions annotated in the S288c reference as centromeric regions, telomeric regions, or LTRs excluded. First, genomic VCF files were obtained with the Haplotype Caller, and joint genotyping of the gVCF files was applied. After joint calling, in the resulting VCF files, only SNPs or only indels (insertions and deletions) were selected. SNPs were filtered according to the parameters (Fay *et al*. 2019): `QD < 5.0; QUAL < 30.0; SOR > 3.0; FS > 60.0; MQ < 40.0; MQRankSum < -12.5; ReadPosRankSum < -8.0`. INDELS were filtered according to the parameters `QD < 5.0; QUAL < 30.0; FS > 60.0; ReadPosRankSum < -20.0`. Indels were then left-aligned. For the final VCF files, indels and SNPs were merged, filtered and non-variant sites were removed. Combined called VCF files were uploaded to FigShare (doi: 10.6084/m9.figshare.28365482).

Variants in the individual strains were selected and exported to a .csv file using the query option of SAMtools/BCFtools 1.10.2. Allele frequency plots were obtained from these using a custom pipeline in R (R Core Team 2021) using a for loop to calculate allele depth ratios from the exported .csv (defined as the ratio of each allele’s sequencing depth to the sum of all alleles’ depth at a given locus) for every genome and the ggplot2 package (Wickham 2016) for data visualization. Depth ratio values for all alleles were plotted for each variant locus along the chromosomes. Allele frequencies were used to estimate ploidy, with the assumption that disomic chromosomes have allele ratios of approx. 1:0 or 1:1, trisomic of 1:0, 1:2, and 2:1, tetrasomic of 1:0, 1:3, 1:1, or 3:1, etc. (Large *et al*. 2020; Rácz *et al*. 2021; Imre *et al*. 2022). The results obtained from allele ratios were compared to coverage plots to determine whether any of the isolates are non-diploids and to identify large deletions and duplications as described in detail in Supplementary File S1. BCFTools’ stats command was used to export per-sample heterozygosity data.

### Identity-by-state (IBS) analysis

IBS analysis was carried out using SNP data as described above with the R package “SNPRelate” 1.24.0 (Zheng *et al*. 2012) using the function “snpgdsIBSNum” with default settings, calculating the minor allele frequency and missing rate for each SNP over all the samples after converting the input VCF file with the snpgdsVCF2GDS function and creating three n-by-n matrices. These contain the number of SNPs sharing zero (e.g. AA and GG homozygous SNPs in compared genomes), one (e.g. AA and AG SNPs), or two identities-by-state (e.g. AA and AA SNPs). It is noted that IBS analysis is developed for diploids and has never been optimized for polyploids. Input VCFs contained diploid format genotypes only and thus the IBS analysis was conducted as if all genomes were diploid euploids. Results were visualized with the R package “heatmaply” (Galili *et al*. 2018) with default settings. Values used for heatmaps are uploaded to FigShare (doi: 10.6084/m9.figshare.28369349) in a table format.

### Phylogenomics based on SNP matrices and Alignment and Assembly-Free method

The VCF file from cohort calling as described above was used to produce genotype matrices using vcf2phylip (Ortiz 2019), using heterozygous nucleotide IUPAC codes where appropriate. The matrix was imported into SplitsTree 4 (Huson and Bryant 2006) in which the Uncorrected P distance was applied and heterozygous sites were averaged (importantly this step allowed us to visualize neighbor nets that took the many heterozygous positions of polyploids into account, as well as variants that are not necessarily biallelic). A neighbor-joining tree was also created from the same data in SplitsTree and iTOL was used to visualize the output (Letunic and Bork 2019). The network and tree files were deposited in FigShare (doi: 10.6084/m9.figshare.28365479) along with PDF versions of figures with searchable text (doi: 10.6084/m9.figshare.28365470). Furthermore, alignment and assembly-free (AAF) phylogenomics was applied to create neighbor nets. The AAF approach is based on identifying k-mers and comparing k-mer lists in genomes, usually in FASTQ files without a mapping step, and was shown to be favorable for analyzing polyploids (VanWallendael and Alvarez 2022). It is also noted that AAF methods can accurately capture local haplotypes if variant positions are close and it can also capture heterozygous indels as such a variant will result in multiple different k-mers for a single genome that can be compared with other genomes (see also Supplementary File S1 for details). The SNP-matrix based approach described above cannot achieve this, as homozygous deletions simply result in an unknown allele at a given position, while a heterozygous position having an SNP allele and a deletion allele will simply be scored as a homozygous SNP (see also Supplementary File S1 for details). We used the AAF software (Fan *et al*. 2015) to create k-mer libraries and k-mer based comparisons with the following options: k-mer length 30, minimum occurrence 5 (`-k 30 - n 5`). To avoid possible contaminating sequences in some yeast genomes originating from bacteria or other sources, we did not simply use AAF on FASTQ files but converted the mapping BAM files from the eight species mapping to FASTA files with SAMTools and used these. This method eliminated contaminations, but at the same time removed any possible introgressions from other genera as well. Based on initial optimization, this tradeoff was deemed useful and necessary. The distance values (their calculation is described in more detail in Supplementary File S1) from the k-mer comparisons with AAF were converted to similarity values (calculated as: 1 - distance) and used as input for SplitsTree to create an additional neighbor net representing relatedness among the genomes. From the AAF analysis, the total number of k-mers in each genome and shared k-mers between genomes were also extracted (see also Supplementary File S1 for details).

## Results

### Phylogenomic approach: From clades to clonal clusters

In this study, 49 novel yeast genomes from baker’s yeasts and sourdough samples have been sequenced with short-read technology, including with four spore-clone cultures of a baker’s yeast, UDeb-BY0012. The genome of the UDeb-BY0012 isolate has also been assembled and phased using both high-coverage short-read and long-read sequencing (Supplementary File S1). We also included genomes from the literature, which were members of the major described clades (94 genomes), and an extensive collection of previously sequenced baker’s yeast and sourdough yeast samples (184 genomes). In addition, three newly sequenced and 91 previously published human/clinical isolates were included in our analysis, these all represented the M3 Mosaic and Mixed origin groups. We conducted a multi-faceted phylogenomic analysis with these altogether 421 genomes, using SNP-based phylogenomic networks, dendrograms, and a similarity network based on alignment and assembly-free phylogenomics. As described in detail in Supplementary File S1 this combined approach was designed to focus both on phylogeny and on small-scale differences especially among tetraploid genomes. In several cases, genomes allocated to mosaic groups or not classified anywhere proved to branch together with described clades, these were tentatively delineated as regions related clades (*e.g.* aff. Mixed origin, see below). These regions of the dendrogram or network do not necessarily make a monophyletic group with a given clade, but represent a group of closely related genomes that are not assigned to existing clades.

In some clades, especially in the case of the numerous Mixed origin clade yeasts, the analyses distinguished subclades, i.e. separated lineages within the given clade. As these appeared to show consistent patterns in origin and in genome structure variations, subclades and the genomes within these were closely inspected, while other baker’s yeasts with more distinct phylogenetic position were omitted from further analyses. Within the subclades, clusters of apparently clonally identical yeast genomes were often identified. These showed negligible differences on the level of SNPs and were characterized by long parallel branches on networks, suggesting unclear branching order and in general, extremely close evolutionary relationships. Our assessment that they are clonal versions of a lineage was corroborated by the fact that these clusters included isolates from different batches of the same product. However, these never included genomes from sporulated strains (which are by definition non-clonal but meiotic descendants of a given yeast culture). The approach to delineate these groups of yeasts at various levels of relatedness is summarized in Figure 1. and Table 1. and their characteristics are described in the sections below. In line with the aims of our study, subclades and clonal clusters give the framework for our further analyses. Subclades and clades are marked with unique colors in the figures and supplementary tables throughout this work.

**Figure 1.**
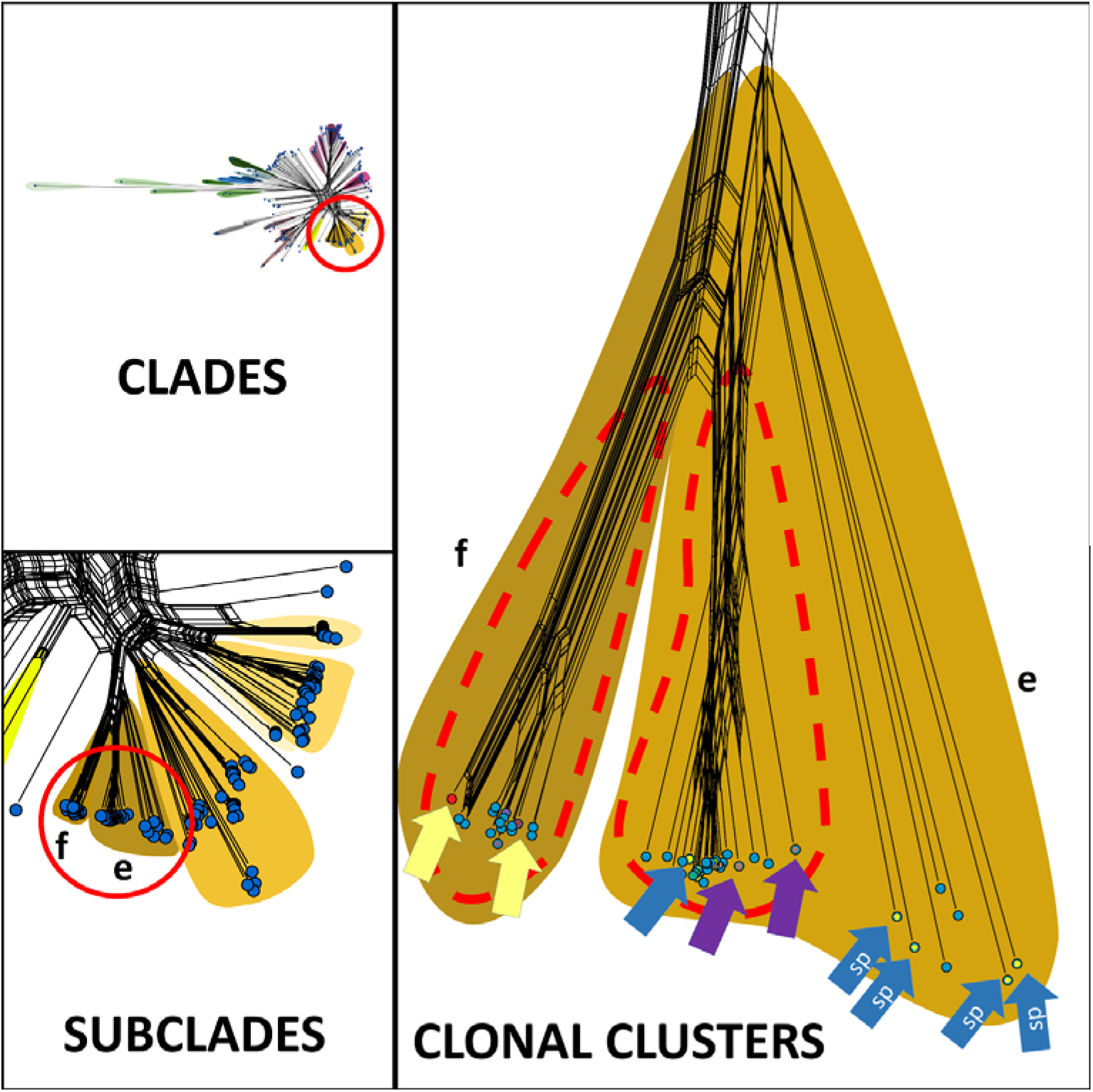
Illustration of clades, subclades, and two clonal clusters (delineated with red dashed lines), using Mixed origin clade (Peter et al. 2018), subclades ‘e’ and ‘f’ as examples (marked on the panels). Arrows in clonal clusters with the same color represent genomes from two batches of the same product that branch separately in the network. Note that other isolates branch between these genomes that are known to be clonally identical. Blue arrows marked with ‘sp’ show spore culture genomes from a yeast (UDeb-BY0012) in the neighboring but separated clonal cluster (blue arrow), to illustrate that sporulated yeasts are not inside clonal clusters and are well-separated from clusters and often each other in the network. Note the long, parallel branches in the clonal clusters (unclear branching order) and the simple, separated long branches placing two pairs of spore clones from a single tetrad distant from each other and from the clonal clusters.

**Table 1.** Delineation and features of clades, regions associated with clades, subclades, and clonal clusters as defined in this study.

|  | Clades and regions | Regions associated with clades | Subclades | Clonal clusters |
| --- | --- | --- | --- | --- |
| **General description** | The clades and mosaic regions as described in literature. | Genomes branching close to known clades, may be paraphyletic or monophyletic, not (yet) named in literature. | Apparently monophyletic groups of genomes branching within clades, containing several genomes. | Monophyletic cluster of genomes with negligible genetic distance apparently originating from clonal/mitotic expansion of a single progenitor culture. |
| **Delineation strategy in this study** | Following previous phylogenomic analyses. | Comparing position of genomes related to clades in phylogenomic networks (long parallel edges splitting off with a small genetic distance to known clades) and branching on dendrogram (branching near clades). | Comparing major within-clade branches on dendrograms and alternative edges in networks: a subclade's genomes show many within-clade edges and longer parallel edges connect them to other subclades. | Comparing branching order and distances on dendrograms (known clonal replicate genomes branching separately); comparing relatively long parallel edges and short dense edges within a subclade; comparing alleles in IBS1+2, SNP distances, and AAF distances to show very close relatedness. |
| **Phyletic origin** | The macroevolution of the species. | The macroevolution of the species, monophyly relating to clade not unequivocal. | The macroevolution of clades. | Mutations and microevolution in clonal reproduction during propagation of yeast cultures and during their application in fermentation. |
| **Genetic diversity** | High, structured, except for some clades consisting of only a few described genomes. | Variable. | Variable, by definition lower than in clades. | Very low in terms of nucleotide mutations, but may be considerable for microevolutionary phenomena, like genome structure variations, aneuploidies, LOH. |

### Phylogenomics confirms the relatedness of European sourdough isolates to Asian ones

The present phylogenomic analysis of 421 *Saccharomyces* isolates included yeasts from the bakery and sourdough environment and clinical samples of the Mosaic 3 group and the Mixed origin clade (Peter *et al*. 2018; Bigey *et al*. 2021). Phylogenomic network analysis using SNP data (Figure 2a) and k-mer data (alignment and assembly-free phylogenomics, Figure 2b), and a neighbor-joining tree for all genomes (Figure 3) clearly identified the clades and mosaic groups previously described in literature (Peter *et al*. 2018). The mosaic M3 region, which has already been defined as a polyphyletic assemblage, however, was split into distinct regions (two in SNP-based network, and three in AAF-based).

**Figure 2.**
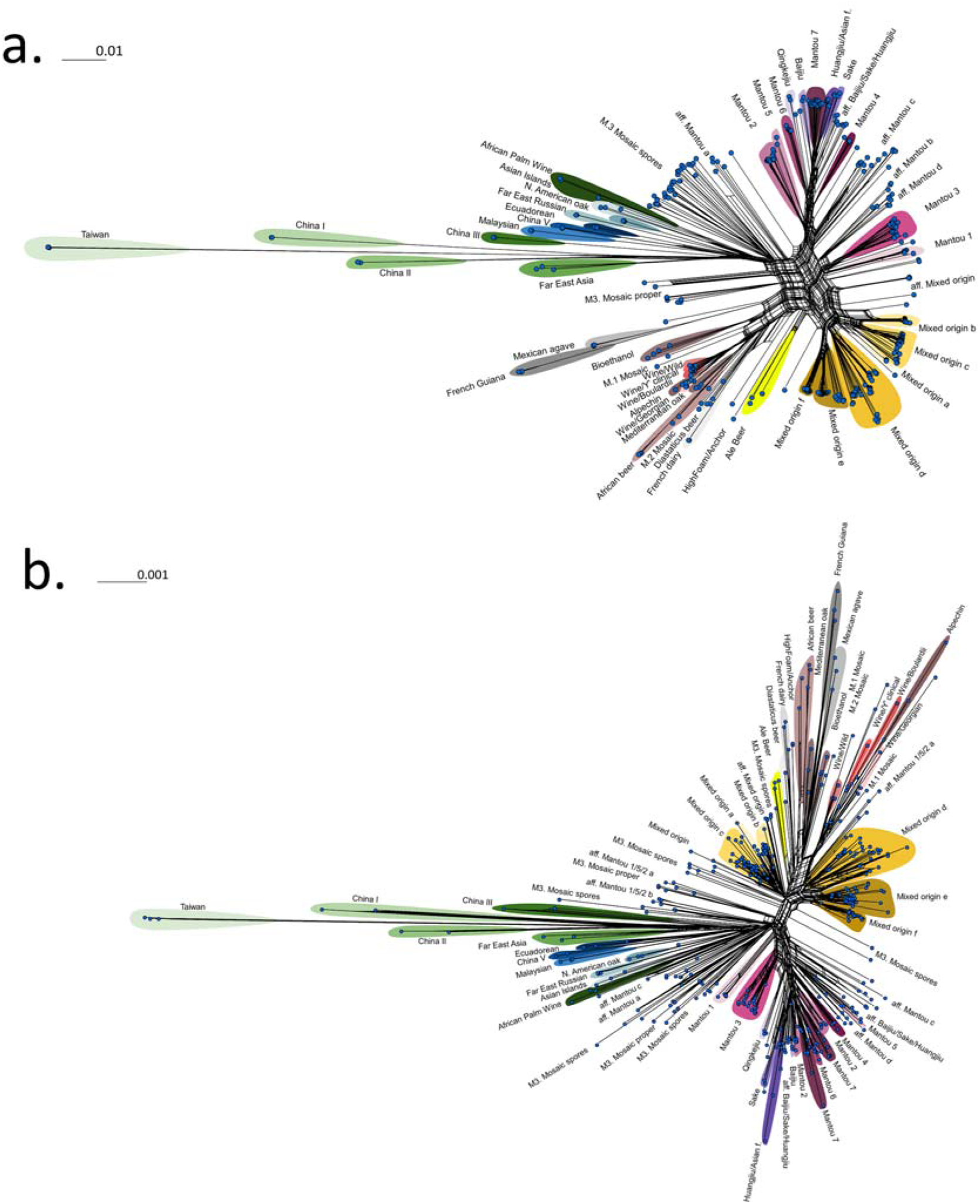
Phylogenomic networks for the genomes analyzed in this study representing the previously described clades of the species and an extensive collection of yeasts from the bakery environment and from human samples. Clades are marked with different background colors; mosaic groups are indicated without background. **a)** Network based on allele calling and SNP-based similarity. **b)** Network based on AAF analysis and k-mer based similarity scores calculated as described in Supplementary File S1.

**Figure 3.**
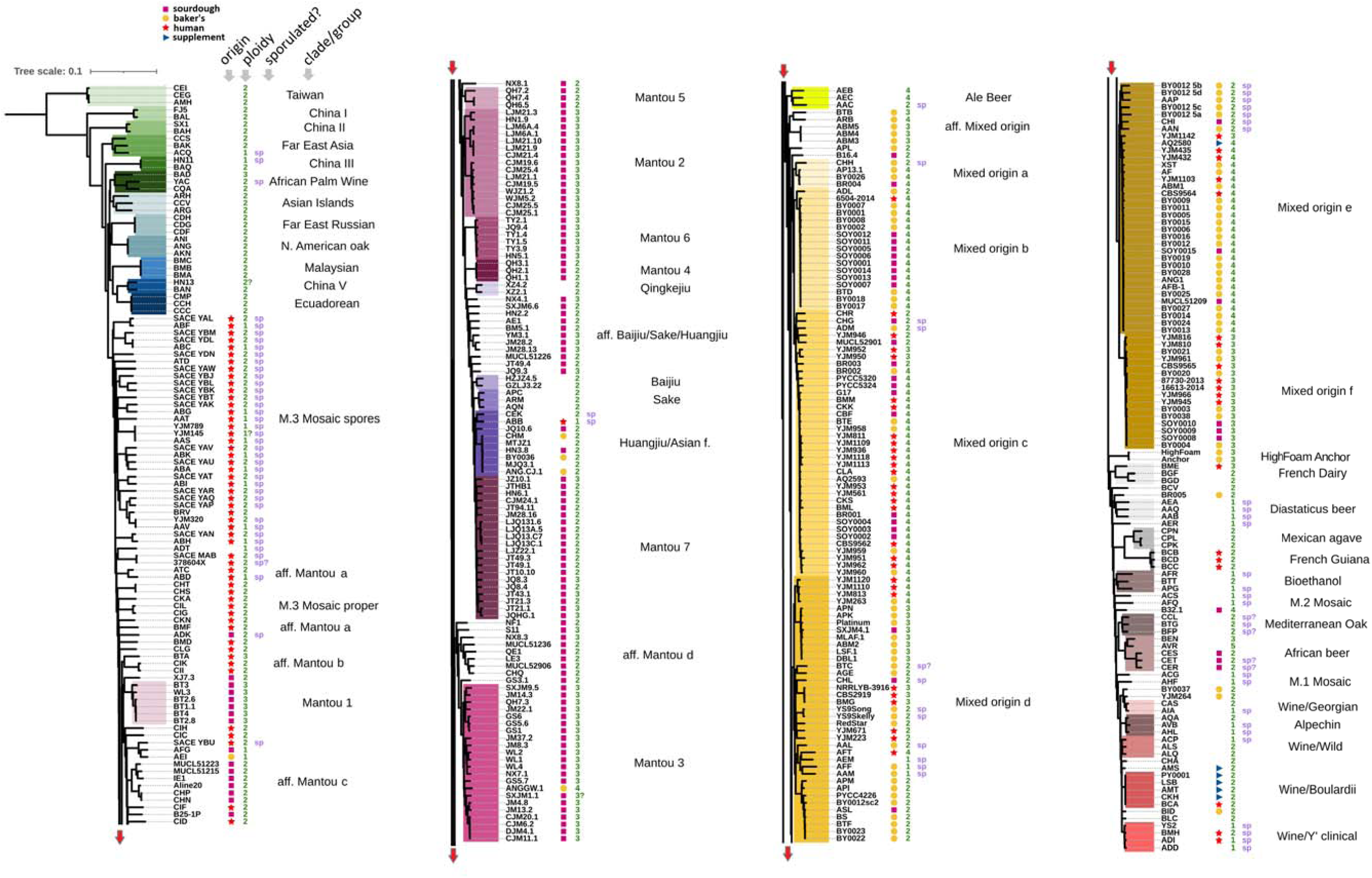
Neighbor-joining dendrogram obtained from SNP-data, with clades, subclades, and various regions highlighted. Information on isolation source (in the case of sourdough, baker’s yeast products, dietary supplements or probiotics, and human hosts), ploidy (values in green), and whether the yeast sample is derived from sporulation (marked as ‘sp’) is given. A PDF version of this image with searchable text is uploaded to the FigShare folder of this work.

Sourdough isolates, including those sequenced and placed in the Mosaic 3 region by Bigey et al. (2021) mostly clustered in the different – and often not closely related – Chinese Mantou clades as well as in the Mixed origin clade (Figures 2–3). Furthermore, four regions related to the Mantou clades, but necessary monophyletic with them, were identified based on the phylogenomic network and the NJ tree based on SNP data (Figure 2a, Supplementary Table S2). These also showed clear separation in the AAF phylogenomics (Figure 2b), thus they were analyzed separately and were given temporary identifiers (‘a’ to ‘d’). The branching positions of these regions differed in the SNP and AAF analyses. These newly delineated regions mostly contained human isolates (named “aff. Mantou a” with one sourdough and six clinical isolates, and “aff. Mantou b” with three clinical isolates), or mostly sourdough isolates (“aff. Mantou c”, with 11 sourdough genomes and one clinical isolate, and “aff. Mantou d”, with nine sourdough isolates). One region related to Baijiu, Sake and Huangjiu clades was also identified, containing solely sourdough isolates (named here “aff. Baijiu/Sake/Huangjiu”, with 11 sourdough isolates). The origin of the isolates in the various clades and groups is depicted in Figure 4 with geographic context. Among the Mantou clades and the above-mentioned Mantou-related regions, sourdough isolates from East Asia were dominant, but a high number of European sourdough isolates also clustered with these. The “aff. Mantou c” contained European isolates predominantly (Figure 3, altogether ten European sourdough yeasts) and European sourdough isolates were also found in the “aff. Baijiu/Sake/Huangjiu” and in the “aff. Mantou d” region.

**Figure 4.**
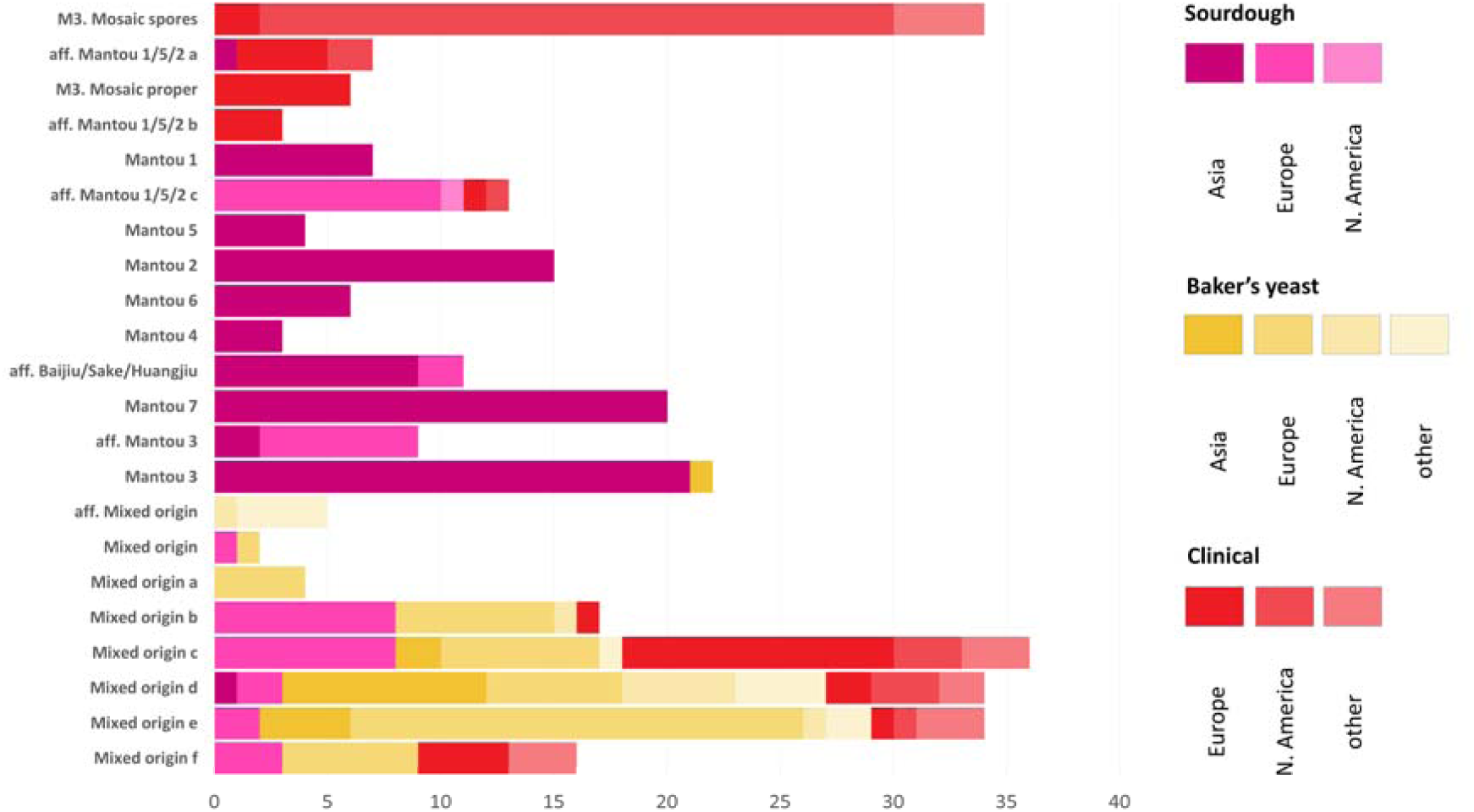
Isolation sources for the various subclades and regions of sourdough and baker’s yeasts. Origin from sourdoughs, baker’s yeast products, or from human/clinical samples is shown in the form of stacked bars, shades of colors display geographic origin as explained on the legend on right side of the figure.

Surprisingly, none of the sourdough or bakery isolates clustered with the M3 Mosaic region after the above-mentioned Mantou-related regions had been delineated. The M3 region only contained clinical isolates (Figures 2–4) and separated into two groups of genomes: one with genomes all derived from single-spore isolates (Supplementary Tables S1–S2) and one with samples that are not single-spore clinical isolates (M3 “proper” region).

### Phylogenomics of sporulated yeast samples

Since the separation in the M3 Mosaic region clearly correlated with whether the sequenced yeast was a spore-clone or not, we investigated the phylogenomic placement of sporulated yeast in other clades as well. Information on whether a yeast was subjected to meiosis and spore formation before sequencing was extracted from the literature and a tetrad of the tetraploid UDeb-BY0012 Mixed origin baker’s yeast was also sequenced (genomes named UDeb-BY0012 5a, b, c, and d). These members of the tetrad only loosely clustered in the phylogenomic networks and dendrogram (Figures 2–3) and were separated clearly from the unsporulated tetraploid UDeb-BY0012 genome. The latter clustered with several other commercial baker’s yeasts, with extremely low divergence among them. Based on these results, we decided to omit sporulated yeasts from detailed comparisons as their phylogenomic placement is unreliable.

### Phylogenomics of the Mixed origin clade

The Mixed origin clade, containing most of the previously sequenced baker’s yeast genomes, was recovered as a single monophyletic clade with several subclades based on SNPs (Figure 2a), but was split into separated clades in AAF network analysis (Figure 2b). As shown in examples in Supplementary File S1, the SNP- and AAF-based similarity networks are built on different approaches and calculations, thus this difference in the positioning of the Mixed origin clade is not contradictory. Nevertheless, we could delineate subclades based on SNP-data and AAF-analysis that corresponded to each other. These subclades were given single letters (‘a’ to ‘f’, Figures 2–3) as identifiers for a more detailed analysis (Supplementary Table S2). A closely related region of genomes was also identified, containing five baker’s yeasts, which we assigned to the “aff. Mixed origin” region tentatively (Figure 3, Supplementary Table S2). The subclade ‘a’ contained merely four baker’s yeast genomes, while genomes in subclades ‘b’ to ‘f’ were more numerous, containing 16 and 36 samples, respectively. These subclades also contained baker’s yeast product isolates, sourdough isolates, and human isolates as well. All newly sequenced sourdough yeasts from Hungary and Transylvania belonged to the subclades of the Mixed origin clade and not to one of the clades associated with sourdoughs, although they were derived from true sourdoughs with otherwise diverse microbiota. These sourdoughs contained Lactobacilli and *Acetobacter* species, and in three cases non-*Saccharomyces* yeasts as well, including *Pichia kudriavzevii* and *P. membranifaciens* (in the Transylvanian traditional sourdough), and *Maudiozyma humilis* (in two commercial bakery sourdoughs). A more detailed analysis of the microbiota of these sourdough samples is uploaded to FigShare (doi: 10.6084/m9.figshare.28193456 and 10.6084/m9.figshare.28193495). Information on which newly sequenced sourdough yeast originates from which sourdough is described in Supplementary Table S2. None of the sourdough samples harbored isolates of more than one subclade of the Mixed origin clade.

### Phylogenetically unique yeast isolates in the baking environment

The phylogenomic comparisons identified several yeast samples from the bakery environment that had unique placement. The Huangjiu/Asian fermentation clade contained Asian sourdough and baker’s yeast samples, as well as a European baker’s yeast and a yeast isolated from a commercial bread mix (Figure 3, Supplementary Table S2). The commercial baker’s yeasts HighFoam and Anchor formed a clade well-separated from any other named clades, while the yeast BR005 in a singleton position and proved to be a *S. cerevisiae* × *S. kudriavzevii* hybrid. One sourdough sample clustered in the M2 Mosaic region, and three sourdough genomes in the African beer clade (Figure 3, Supplementary Table S2). Three yeasts, namely UDeb-BY0037 (from commercial frozen dough), YJM264, and BID clustered with the Wine/European clade (Figure 3, Supplementary Table S2). Of these, YJM264 was a *S. cerevisiae* × *S. kudriavzevii* hybrid as described previously (Zhu *et al*. 2016), while the newly sequenced UDeb-BY0037 proved to be a *S. cerevisiae* × *S. uvarum* hybrid. None of the yeasts from the bakery environment clustered with the probiotic subclade of the Wine/European (it is noted that the dietary supplement yeast AQ2580 clearly clustered into the “Mixed origin e” subclade, not in the probiotic yeasts’ subclade). The genomes of the above-listed unique yeasts in the baking environment were not included in subsequent analyses due to their unique phylogenomic placement and sporadic occurrence in our dataset.

### Ploidy and clonal clusters among true sourdough clades

The phylogenomic analysis of the genomes included in this study not only helped to determine the placement of sourdough and baker’s yeast genomes, but also revealed conspicuously similar clusters of genomes in various subclades, especially in the case of the Mixed origin clade (as in Figure 1), but also to a smaller extent in the Mantou clades. As described in detail in Supplementary File S1, the ploidy, aneuploidies, and gross chromosomal rearrangements were determined using a combination of coverage and allele depth data. Information on ploidy was incorporated onto the phylogeny (Figure 3) and was compared among subclades and clonal clusters first among the Mantou clades and related regions. Some of the yeasts in Mantou 1 (one cluster of triploids), Mantou 2 (three clusters of triploids), Mantou 7 (two clusters of diploids), Mantou 3 (two clusters of triploids), and “aff. Mantou c” (one cluster of diploids) showed very low divergence and can be considered clonal groups despite the fact that in several cases, they were collected from various provinces and regions of China (Supplementary Tables S1-2). Furthermore, the Mantou 6 (triploids) and Mantou 4 (diploids) clades’ genomes were so similar that these likely correspond entirely to clonal groups (Figure 3). As none of the yeasts in these clonally similar clusters originated from commercial yeast products or from human hosts, the Mantou clonal clusters were not investigated in more detail. Genomes assigned to clonal groups of these clades are listed in Supplementary Table TS2.

### Ploidy and clonal clusters among commercial baker’s yeasts

The identification of subclades within the Mixed origin clade and the fact that most of these contained yeasts that originated from all three major isolation sources (sourdough, baker’s yeast, clinical) enabled us to conduct a detailed comparative genomic analysis focusing on the most closely related groups of isolates, the identified clonal clusters. The Mixed origin “a” subclade contained tetraploid yeasts and a single, sporulated, diploid one. The subclade “b” also contained tetraploids with a single, separately branching genome showing diploidy. The subgroup “c” was characterized by the abundance of tetraploid genomes and a few, more dissimilar triploid and diploid genomes, some of them sporulated. The Mixed origin “d” subgroup contained four clusters, and fine phylogenetic relationships were recapitulated in ploidy differences. Tetraploid, triploid, and diploids clusters were identified as well as haploid sporulated genomes. Similarly, the subgroup “e” also had a cluster of tetraploid isolates, and several, more separately branching diploid sporulated isolates, including members of the UDeb-BY0012/5 tetrad.

To ensure that the clonal clusters identified using the phylogenomic analyses indeed contain genomes that are similar to a level that implies clonal isolates from a single original strain, we analyzed variants in identity-by-state 0 (completely different variant positions, Figure 5a) and in IBS 1 or 2 (positions where one allele is identical, or two alleles are identical between two genomes, respectively, Figure 5b–c). Furthermore, we compared the similarity values based on AAF analysis (Figure 5d) and the amount of shared k-mers (Figure 5e). The clonally similar clusters were well-defined in all cases, meaning that these indeed contained genomes that are almost indistinguishable in the phylogeny, and are characterized by an almost complete lack of alleles in IBS 0, and a very high prevalence of alleles in IBS2. Similarly, k-mer based similarity and total number of shared k-mers also make them extremely highly similar. As all 421 genomes were included in these analyses, the Mantou clades could also be assessed according to identity of variant by state and according to k-mers (Figure 5), highlighting the extremely high similarities among the genomes of the previously identified Mantou clonal clusters. The Mantou 3 clade showed an interesting pattern of relatively high similarity with other Mantou clades and also with the Mixed origin clade in the case of IBS0 and IBS2 alleles and shared k-mers. Their AAF-based similarity to other Mantou clades was markedly higher than to the Mixed origin clade.

**Figure 5.**
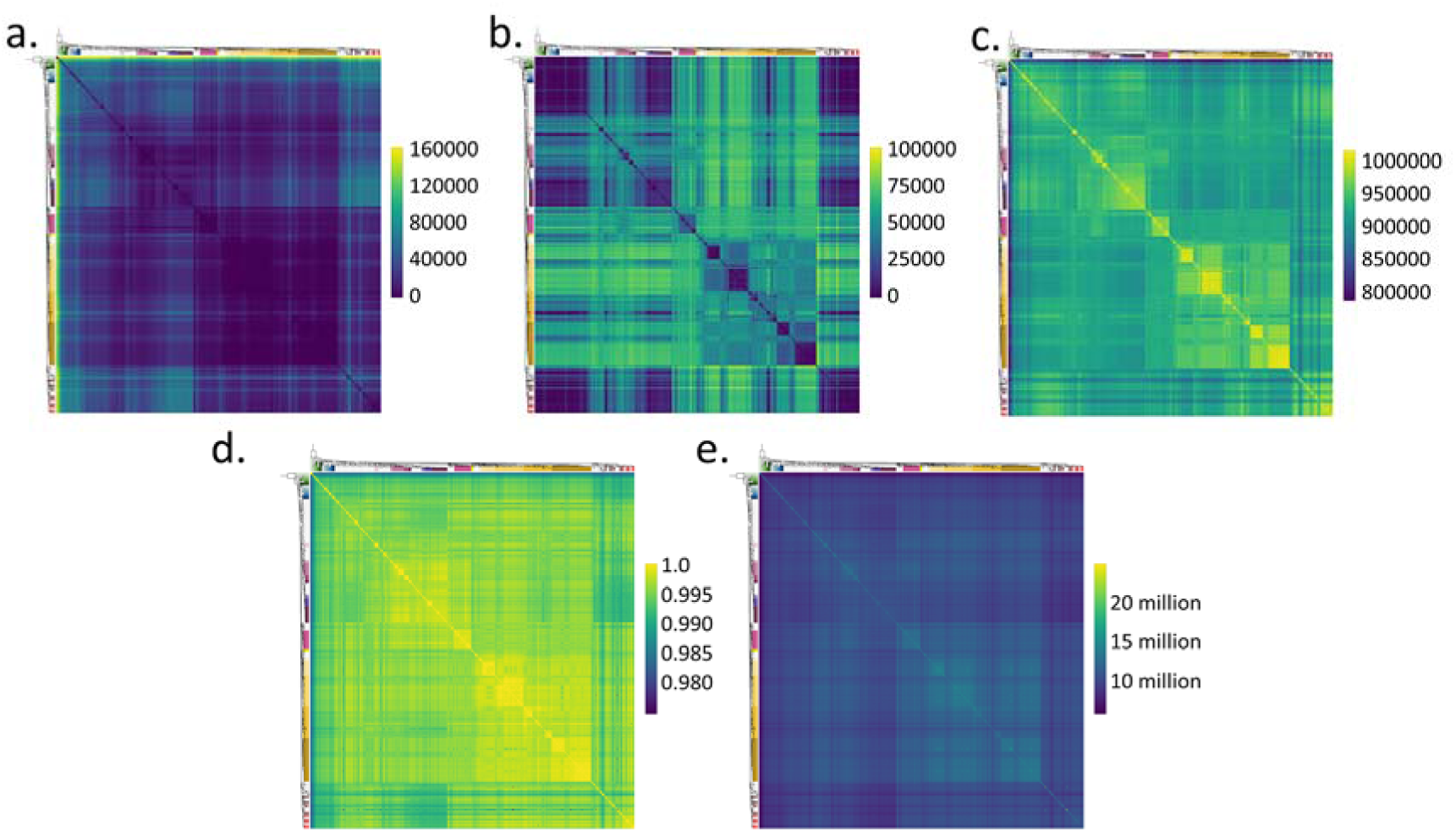
Heatmaps illustrating similarities among and within clades. The order of genomes corresponds to the phylogeny of Figure 3, the dendrogram is added to the left and top edges of heatmaps. Each heatmap’s scale is added on the right side. **a.** Alleles in IBS 0 state among the genomes. **b.** Alleles in IBS 1 state among the genomes. **c.** Alleles in IBS 2 state among the genomes. **d.** AAF-based similarity values among the genomes, as described in Supplementary File S1. **e.** Number of shared k-mers among the genomes.

Data on clonal clusters is summarized in Supplementary File S2, including AAF similarity, SNP similarity, and the average of these values among a clonal cluster’s genomes and among the members of a cluster and other genomes of the subclade. In all cases, the clonal clusters’ genomes were markedly more similar to each other than to the rest of the genomes in the subclade. Ploidy and genome structure variation data (Figures 6–8, Supplementary Files S3– S4) as well as the origin of isolates/strains and manufacturer where applicable (Supplementary File S5) were all taken into account to give a detailed description on the clonal clusters. The various gross chromosomal rearrangements (GCRs) identified are summarized in Figure 9a. We also conducted electrokaryotyping on the newly isolated yeasts, to identify structural variations that may not be detectable solely from sequencing data (as described in Supplementary File S1). The karyotypes are shown on Figure 9b. Each subclade’s clonal clusters are detailed below.

The “aff. Mixed origin” region and the “Mixed origin a” subclade contained five and four genomes only. In both, clonal clusters were apparent, a very similar group of three triploid aneuploidy genomes in the former (all isolates from Company 1a’s yeast products, with 50644–51653 heterozygous SNPs), and three tetraploid genomes in the latter with various aneuploidies (with 70190–71526 heterozygous SNPs). Although the yeasts in the “a” subclade originated from various European countries, they all contained a unique GCR, a terminal duplication of a region on chr. XI, starting at an ARS (autonomously replicating sequence) (Figures 6 and 9a; Supplementary Files S2-5).

In the tetraploid “Mixed origin b” subclade, all but one genome was found to be so similar that they were assigned to a clonal cluster (Supplementary File S2). Among these 16 genomes, a clinical sample from Hungary, seven commercial baker’s yeast isolates (from Company 12d from Hungary, Company 12c from Canada, from Company 10 from Germany, and from an undisclosed manufacturer) and eight sourdough isolates from Hungary were found (Supplementary File S5). Aneuploidies were detected in eight of the 16 genomes in the cluster, affecting both baker’s yeast and sourdough isolates (one to two aneuploidies per genome) and the clinical isolate 6504/2014 as well (with three different aneuploidies) (Figure 6, Supplementary Files S2-4). Heterozygosity was high in the subclade (70119–74870 heterozygous SNPs, the lowest number was found in the clinical isolate). The yeasts in this clonal cluster showed an almost ubiquitous genome structure variation, namely a deletion of one out of four copies of an intergenic region from the left telomere on chr. VII (Figure 9a, Supplementary File S4). This terminal deletion was not detectable in the clinical isolate 6504/2014. The latter isolate had a terminal duplication affecting one of four copies of chr. IV and a terminal deletion of one of five copies of chr. XV. The breakpoints for these were the genes *EFT2* and *EFT1*, respectively, which are ohnolog genes (Byrne and Wolfe 2005) (Figure 6a, Supplementary File S4). This raises the possibility of a reciprocal translocation between the two chromosomes followed by unequal distribution of the affected chromosomal copies. The ten “Mixed origin b” subclade samples subjected to karyotyping all showed complex karyotypes with supernumerary bands compared to the reference karyotype, and variable bands that occasionally differed between genomes in the same clonal subcluster. Such examples include UDeb-BY0007 and UDeb-BY0008 (isolates from two batches of the same product with differing aneuploidies), or UDeb-BY0017 and UDeb-BY0018 (euploid isolates from two batches of the same product) (Figure 9b).

**Figure 6.**
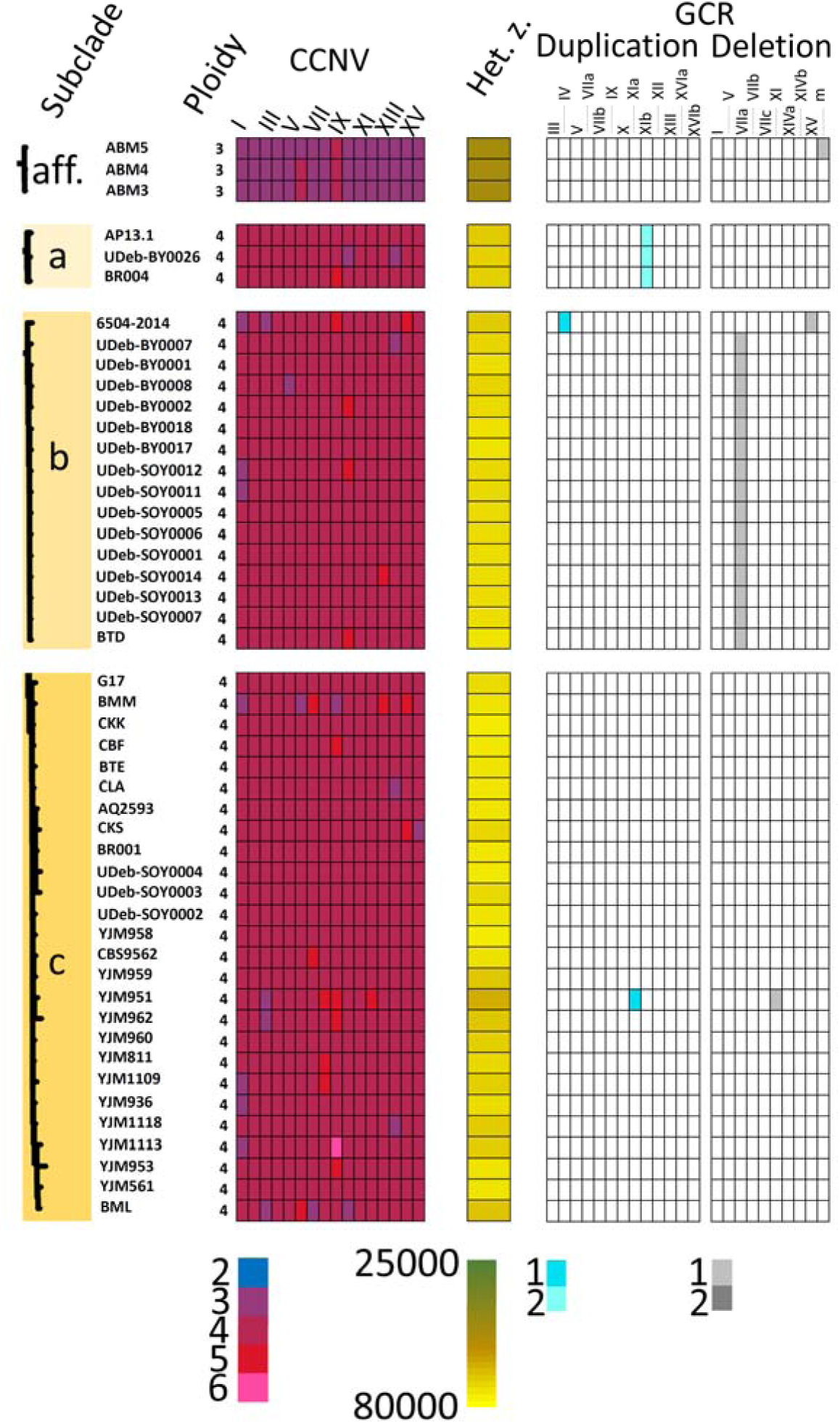
Comparison of the genomes in the clonal clusters of the aff. Mixed origin region, and subclades “a”, “b”, and “c” of the Mixed origin clade. Subclades are shown with close-ups of their dendrogram sections from Figure 3. Ploidy is indicated, and chromosomal copy number variations (CCNVs) are shown with color-codes. In the next column, heterozygous SNPs are shown color-coded. Finally, Gross Chromosomal Rearrangements (GCRs) identified are shown, their copy numbers are also color-coded.

In the “Mixed origin c” subclade, the single clonal cluster contained 26 genomes. It consisted of five sourdough isolates (three from Hungary, and one each from Morocco and Italy), six baker’s yeasts from two known manufacturers (Company 12a, Company 1a, from Italy and Spain, respectively, and the rest from unknown manufacturer) as well as 15 clinical isolates, mostly from Italy and from the USA (Supplementary File S5). The genomes in the clonal cluster were tetraploids, with prevalent aneuploidies affecting various chromosomes (up to six chromosomes per genome affected by aneuploidies). Heterozygosity was high as in the subclade ‘b’ (64857–76132 heterozygous SNPs) (Figure 6, Supplementary Files S2-4). Other structural variants, however, were only found in a single genome, YJM951, affecting two chromosomes. This sample had the lowest heterozygosity in the cluster (Figures 6,9a, Supplementary File S4). The samples collected for this study, namely the three Hungarian sourdough yeasts, were subjected to karyotyping and had mostly identical karyotypes and no supernumerary bands in the region of small chromosomes (Figure 9b).

The “Mixed origin d” subclade contained the highest number of genomes in the clade, was more diverse genetically, and was not dominated by a single clonal cluster of nearly identical genomes. Clonal groups with various ploidies were delineated, and overall, 26 of 37 genomes were assigned into five separate clonal groups (Supplementary File S2). A clonal group containing only the two replicates of the YS9 strain with relatively low sequencing coverage was omitted from subsequent analysis. The first clonal cluster containing three clinical isolates (two from the USA) and a single baker’s yeast (YJM263, Fleischmann, collected before 1994) was characterized by tetraploidy, several aneuploidies, high heterozygosity (70701 to 71912 heteryozygous SNPs) and no GCRs (Figure 7, Supplementary Files S2–4). The second cluster containing the genomes APN, Platinum, MLAF.1, ABM2, LSF.1, APK, SXJM4.1, and DBL1 consisted exclusively of triploid baker’s yeasts (from China, Philippines, USA, and South Africa) and a single sourdough isolate from China. These had lower heterozygosity (47221 to 54153). Aneuploidies were found in all but one of these (APK, the genome with the highest heterozygosity), and a terminal deletion on chr. I was prevalent, affecting all except the same APK sample. The deletion started at a delta LTR sequence and ended at the right telomere (Supplementary File S4, Figure 9a). Three genomes shared a combination of two neighboring deletions of different copy number on chr. XIV, associated with LTR and ARS sequences and reaching the telomere. A third, smaller clonal cluster was delineated containing three versions of the same triploid clinical isolate Dietrichson V-253 from Norway sequenced by three different studies, originally deposited in collections before 1954. The three versions originated from the CBS, NCYC, and NRRL collections and even though they represented the single original isolate, they differed in some aspects. They displayed different aneuploidies and other genome structure variations unique to these genomes (Figure 7, Supplementary Files S2–4). Their heterozygosity was, however, remarkably similar (45365 to 45629 SNPs). The fourth clonal cluster in the “Mixed origin c” subclade was peculiar among members of the whole clade as it contained diploids with aneuploidies (one, the trisomy of chr. X was ubiquitous, four other chromosomes had occasional trisomies). GCRs were only found in the UDeb-BY0012-sc2 yeast that was isolated from the same batch as the UDeb-BY0012 tetraploid in the “Mixed origin e” subclade (Figure 7, Supplementary Files S2-4). Eight genomes were from baker’s yeasts from Korea, Japan (Company 4), China, and Germany (Company 8), and a single genome from a Turkish sourdough (Supplementary File S5). This clonal cluster had the lowest heterozygosity (number of heterozygous SNPs 25354–37587) with the lowest heterozygosity observed in the UDeb-BY0012-sc2 sample that also showed the GCRs. Two isolates, UDeb-BY0022 and BY0023 were subjected to karyotyping and despite their diploidy, they showed complex karyotypes and supernumerary bands and were not completely identical (Figure 9b).

**Figure 7.**
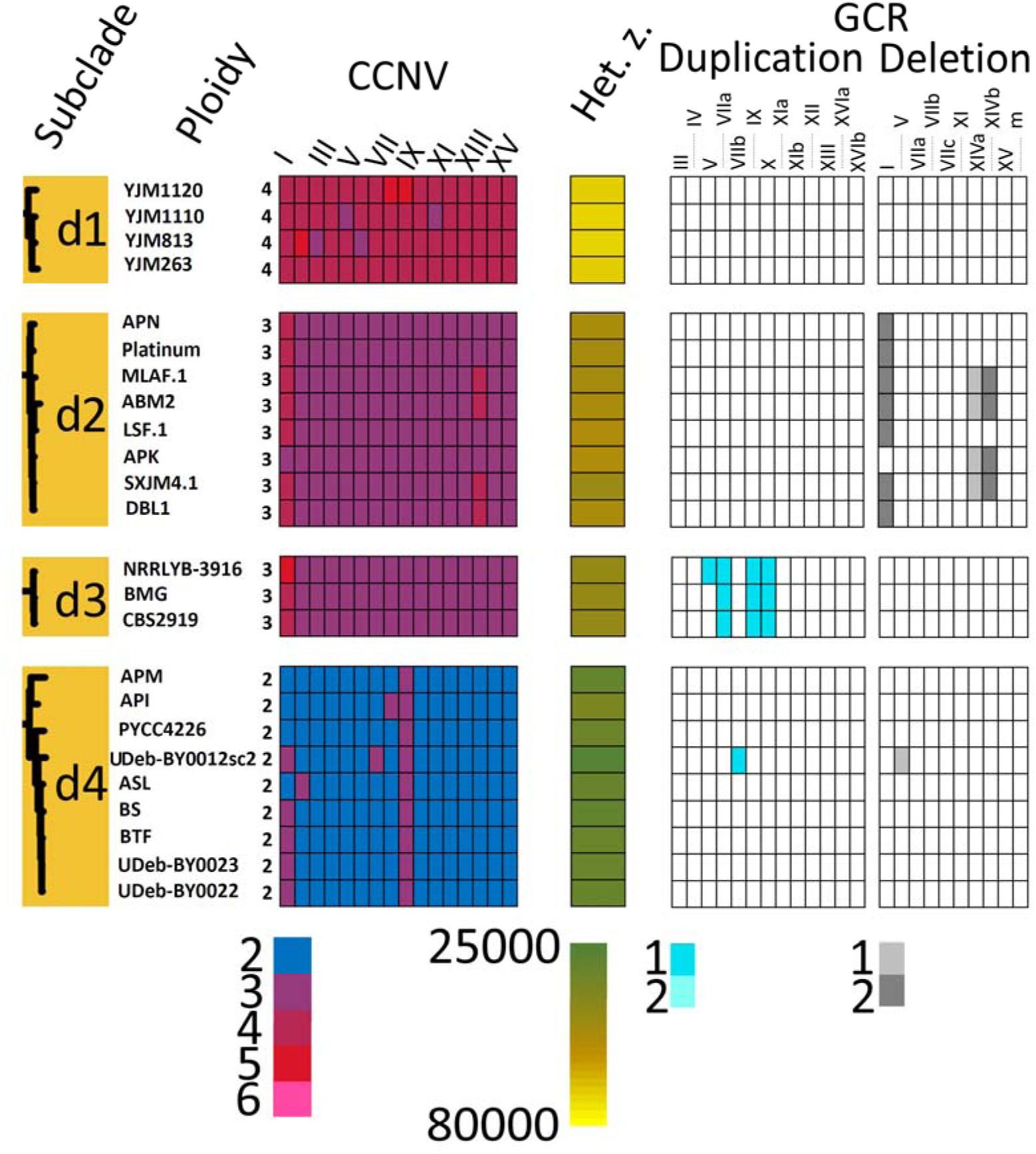
Comparison of the genomes in the clonal clusters of the “d” subclade of the Mixed origin clade. Subclades are shown with close-ups of their dendrogram sections from Figure 3. Ploidy is indicated, and CCNVs are shown with color-codes. In the next column, heterozygous SNPs are shown color-coded. Finally, GCRs identified are shown, their copy numbers are also color-coded.

The “Mixed origin e” clade contained several spore clone genomes (including the UDeb-BY0012/5 tetrad created for this study), and a larger clonal cluster of 28 mostly tetraploid/tetraploid aneuploidy genomes including the UDeb-BY0012 sample (Supplementary File S2). Only the genome YJM1142 was non-tetraploid, but unclassified, conspicuously with nine trisomic and seven tetrasomic chromosomes. Of the 28 samples, 20 were baker’s yeast manufactured by various companies and sold by various retailers predominantly in Central Europe and in China, two were sourdough isolates (one Belgian and one Hungarian), one was a dietary supplement from Spain, and five isolates were clinical ones from Tanzania (the aforementioned YJM1142), Greece, USA, and from unspecified locations (Supplementary File S5, Supplementary Table TS2). Aneuploidies, mostly pentasomies of chromosomes (13 genomes) and a few trisomies (five genomes) were prevalent among the genomes (Figure 8, Supplementary Files S2–S3). In the clonal cluster, 24 of 28 yeasts shared a conspicuous interstitial deletion between two LTRs on chrom. VII affecting one chromosomal copy, and in two additional cases, this GCR could not be evaluated. 26 of the 28 genomes also shared a unique terminal duplication on chr. III starting at the MAT locus and ending on the right telomere, affecting one copy (Figure 8,9a; Supplementary File S3–4). In two cases, this GCR was also not evaluated. Heterozygosity in the cluster was high, ranging from 60364 SNPs (YJM1142) to 71330 (Figure 8). The karyotypes of the altogether 16 samples assessed here (Figure 9b) were complex, with supernumerary and variable bands even among inter-batch replicate samples, e.g. UDeb-BY0005 and BY0006, or UDeb-BY0013 and BY0014.

**Figure 8.**
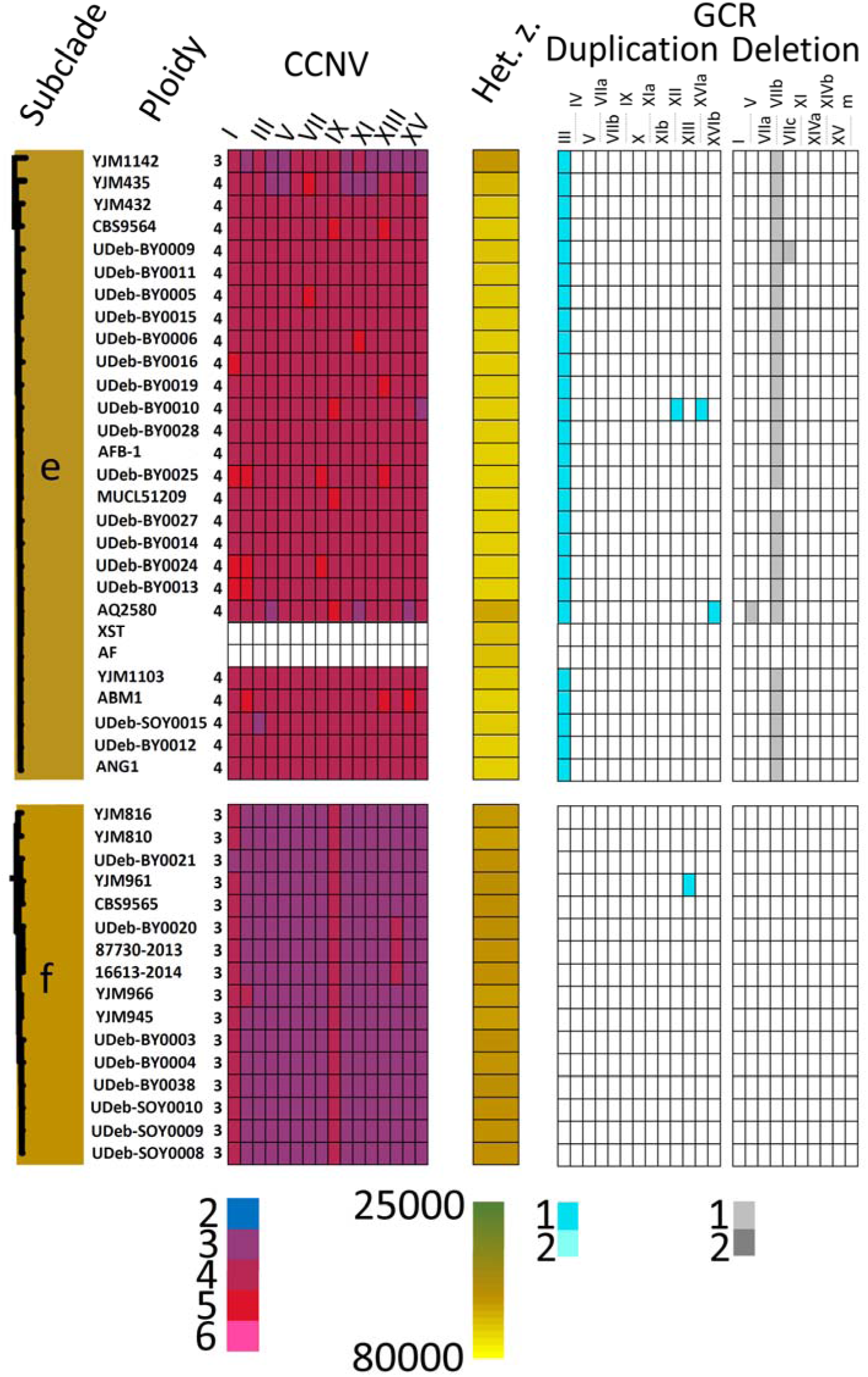
Comparison of the genomes in the clonal clusters of the “e” and “f” subclades of the Mixed origin clade. Subclades are shown with close-ups of their dendrogram sections from Figure 3. Ploidy is indicated, and CCNVs are shown with color-codes. In the next column, heterozygous SNPs are shown color-coded. Finally, GCRs identified are shown, their copy numbers are also color-coded. Structural variants could not be evaluated for two genomes in subclade “e”.

**Figure 9.**
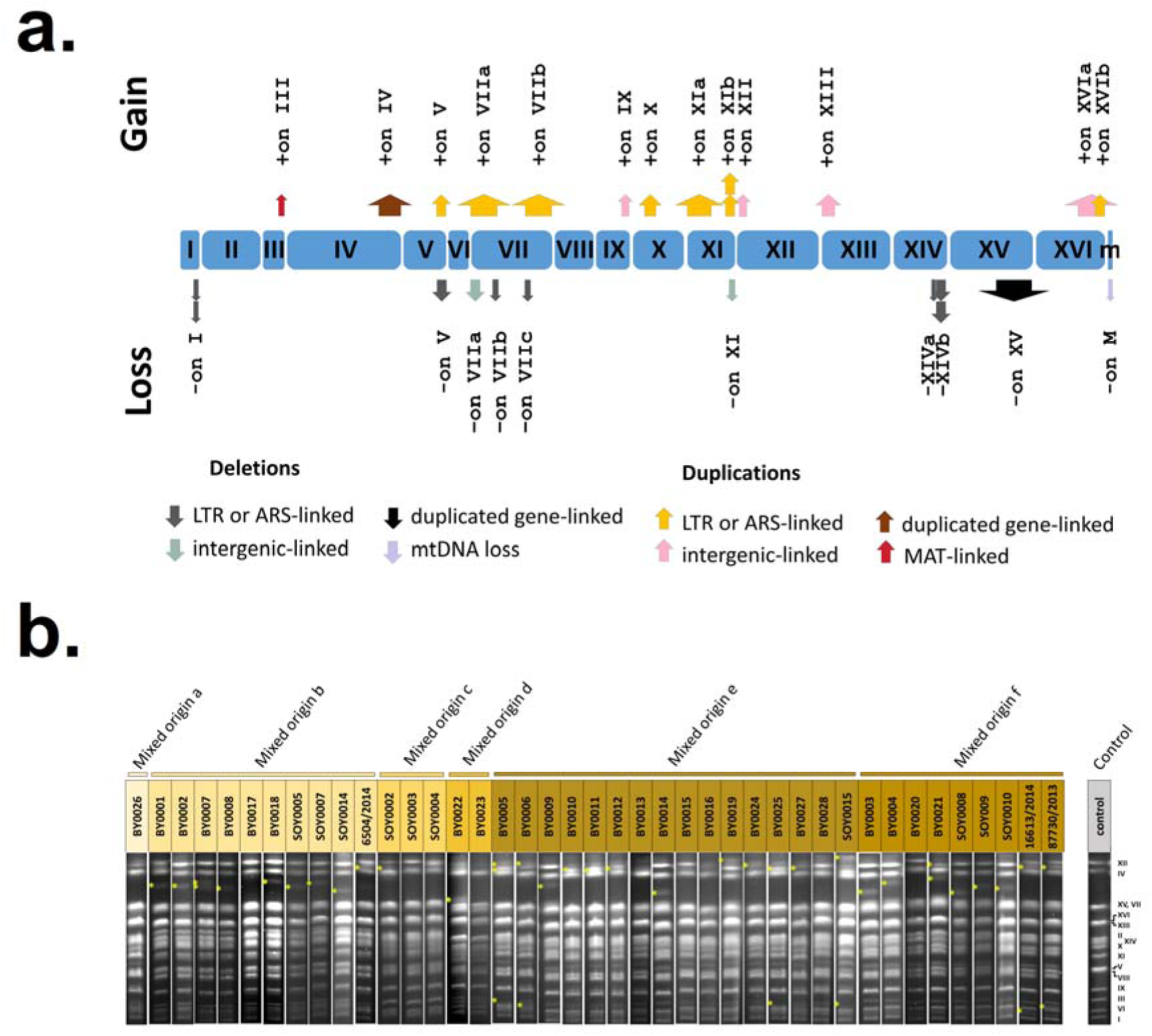
Structural variations observed in the samples with two different approaches. **a.** An overview of the locations of identified deletions and duplications. Chromosomes are represented by blue boxes, and arrows show the regions duplicated or lost according to coverage mapping. Double arrows represent GCRs affecting two chromosomal copies. Multiple GCR events on the same chromosomes are identified by letters a–c. Types of GCRs according to start and end loci are color-coded as shown at the bottom of the image. **b.** CHEF electrokaryotyping chromosomal bands of the samples collected for this study and available in our culture collection. Subclades are color-coded. Asterisks represent variable chromosomal bands; control karyotype is on the right side of the panel. Note that the lanes are cut and rearranged from individual runs each containing a maximum number of ten samples.

The “Mixed origin f” group was in its entirety a triploid-aneuploid clonal cluster with 16 genomes, six of them baker’s yeasts from China, Italy, Ukraine, and Germany (the latter from Company 3c), three sourdough isolates from Hungary, and another six isolates from clinical samples (from Italy, Greece, and Hungary) (Supplementary Files S2,5; Supplementary Table TS2). Tetrasomy of chr. IX was ubiquitous and in the case of chr. I, all but one samples were tetrasomic. A single genome, YJM961 showed a GCR, a terminal duplication on chr. XIII (Figure 8,9a; Supplementary Files S3–4). Heterozygosity was lower than that of the tetraploid clusters, but higher than the ones observed in the diploid cluster (Figure 8), and ranged between 58204 and 61693 SNPs. The karyotypes of the altogether nine samples assessed here (Figure 9b) were similarly complex and variable as in the case of the “Mixed origin e” subclade.

### Sporulation in di- and polyploidy baker’s yeasts

Lastly, we used tetrad dissection to assess the sporulating ability and spore viability of the baker’s yeasts isolated from Hungary. Sporulation percentage varied between 35–58% in six tested tetraploids of the “Mixed origin b” subclade, and none of the dissected tetrads were fully viable. The two tested “Mixed origin d” diploids showed 4–5% sporulation and were not dissected due to their very low number of asci. From the “Mixed origin e” subclade, 15 tested strains showed an overall low sporulation ability, ranging between 4–44%, and only four samples had high enough sporulation for tetrad dissection. However, these had high spore viability and more completely viable tetrads, e.g. the UDeb-BY0012 isolate, from which the tetrad no. 5 originated (that was sequenced in this study). The triploid “Mixed origin f” showed very low sporulation (0–7%) and asci with an aberrant number of spores only. Sporulation data is presented in Supplementary File S6.

## Discussion

### Clonal clusters and phylogeny

In the current study, 421 *Saccharomyces* genomes were used to provide a backbone for a phylogenomic analysis and to identify relationships among baker’s, sourdough, and clinical yeast isolates. As the majority of commercially available baker’s yeast strains have been found to belong to the Mixed origin clade, we focused on this group, along with Chinese Mantou sourdough yeasts and the different clades found within them. We also included clinical and other isolates of the Mosaic 3 group as this group was found to be related to recently sequenced sourdough yeasts (Bigey *et al*. 2021). To add more context to the genomes in the phylogeny, we conducted an extensive review of the literature and online resources to compile information on isolation origin and date of the sequenced yeasts and identified which yeasts have been artificially sporulated in labs before sequencing.

Our phylogenomic networks and dendrogram (Figure 2–3) recovered the clades already described in the literature and showed that some previously unplaced genomes cluster with established clades. Furthermore, well-separated subclades in the Mixed origin clade have been identified. A substantially different phylogenetic placement of the Mosaic 3 (M3) region was apparent compared to previous analysis (Peter *et al*. 2018). Numerous, sporulated M3 strains formed one cluster, while other, non-sporulated strains (designated here as ‘M3 proper’) were separated. A recent population structure analysis of more than 3000 *S. cerevisiae* by Loegler et al. (2024) also did not recover the M3 region, and the genomes formerly associated with it formed well-separated groups. Surprisingly, the clinical isolates of this (former) M3 region showed no close relationship to baker’s yeasts or sourdough isolates at all, thus we did not analyze them in detail in the current study (Figure 4).

The European sourdough strains earlier associated with the Mosaic regions (Bigey *et al*. 2021) proved to be closely related to Chinese Mantou yeasts in our phylogeny, clarifying their phyletic origin (Figure 3). We found that in several Mantou and especially in the Mixed origin clades, clusters of genomes showed remarkable similarity and hence, unclear branching order (Figures 3,5; Supplementary File S2). This suggests that these clusters are made up of clonally identical yeasts (Figure 1), which we will discuss next.

The clonal clusters we propose here contain genomes that are almost indistinguishable based on their shared SNPs and shared k-mers, and generally have the same ploidy. Clonal clusters among Mantou yeasts contained isolates from traditional fermentations, but not from commercially available yeast products or clinical isolates (Supplementary Table TS2). Clonal clusters in the subclades of the Mixed origin clade on the other hand were remarkably diverse in their origin: they often contained baker’s yeasts, sourdough yeasts, and clinical isolates (and in a single case, a dietary supplement) as well. This suggests that commercially available baker’s yeasts are not only closely related to some sourdough (compare to Bigey *et al*. 2021) or human isolates (compare to Zhu *et al*. 2016; Pfliegler *et al*. 2017; Morard *et al*. 2023), but that they are often as similar to them as two batches of a commercial product are to each other (Figures 6–7; Supplementary File S2). This suggests that these clinical strains have only recently colonized humans, while maintaining relatively stable genomes, even in the new and likely stressful conditions inside the human host. We note however, that the clonal clusters in our study only contain genomes found to be conspicuously similar. It is possible that processes like extensive loss-of-heterozygosity, whole chromosome losses, and other large structural rearrangements, especially in heterozygous polyploids (Gilchrist and Stelkens 2019; Large *et al*. 2020; Smukowski Heil 2023; Bautista *et al*. 2024), may have resulted in mitotic offspring lineages in various environments that are too dissimilar (and thus separated in the phylogeny) to be recognized as members of the same cluster.

We also speculated that meiosis in the tetraploid baker’s yeasts may lead to even larger changes in the genome (primarily due to reduction in ploidy) and in phylogeny. Thus, we also sequenced a complete tetrad of the tetraploid UDeb-BY0012 yeast. Results indeed showed that the meiotic spore clone cultures are not just reduced in their ploidy, but unmistakably become separated from the clonal cluster of the parental yeast (Figure 3).

### Strain redundancy and relative stability in clonal clusters

Clonal clusters displaying almost identical genomic characteristics were prevalent in our phylogeny, often with a large number of isolates originating from various geographic settings, and from products of different brands and companies. Thus, strain redundancy in commercial products was high, similarly to what has been observed for commercial wine yeasts (Borneman *et al*. 2016). By assessing company structure, subsidiaries, and brand ownership, we found that most subclades and clonal clusters contain baker’s yeast from multiple companies, even after subsidiary companies were grouped together (Supplementary File S5). The Mixed origin “d” and “e” subclades had strains manufactured by as many as five and six companies, respectively, across the globe. Their country of purchase and in-lab isolation is often not the same as the country of manufacturing. Our assessment of strain redundancy and close relatedness of samples, even when collected decades apart (Supplementary Table S2), shows that the observable diversity and biogeography of the Mixed origin baker’s yeasts is, and has mostly been determined by company practices, international brands, and worldwide trade.

The high number of clonally identical genomes and inter-batch replicates of product isolates allowed us to study aneuploidies and GCRs, karyotype changes, and their prevalence in the Mixed origin yeasts. The novel long-read based assembly representing a large clonal cluster failed to capture the heterozygous structural variations due to inherent limitations (as described in Supplementary File S1). Nevertheless, short-read-based analysis of the clonal clusters found many aneuploidies in line with previous works (Zhu *et al*. 2016; Peter *et al*. 2018) and 23 additional large deletions and duplications of chromosomal regions (Figure 9, Supplementary File S4). Most of these were associated with transposon LTRs and ARS sequences or the MAT locus, in line with previously reported observations of recombination hotspots (Sui *et al*. 2020). A unique pattern observed in the clinical isolate 6504/2018 furthermore suggests a peculiar heterozygous translocation between ohnologs on two chromosomes (Supplementary File S3–4).

Polyploid yeast genomes are thought to undergo a gradual genome stabilization and reduction process, manifesting in chromosome losses and a reduction in ploidy, while at the same time increasing in the number of genome structure variants (Selmecki *et al*. 2015; Wertheimer *et al*. 2016). Similar genome reductions have been observed in pathogenic yeasts, e.g. *Cryptococcus* especially during exposure to stress (Gerstein *et al*. 2015). Temporarily increased CCNVs have also been described as a common microevolutionary process when adapting to antimycotic exposure (Harrison *et al*. 2014; Berman 2016), niches of a host (Forche *et al*. 2019), or industrial fermentation environments (Gorter de Vries *et al*. 2017), among others. In our case, surprisingly, the clonal clusters showed remarkable stability in many aspects. The majority of the members of clonal clusters, regardless of ploidy, habitat (baker’s yeast product, sourdough, or the human host), or collection dates spanning decades, often maintained the exact same heterozygous gross chromosomal rearrangements with the same copy number (Supplementary File S4). Several surprisingly stable CCNV patterns were furthermore observed, but only in the diploids and triploids, not the tetraploids (Figures 6–8; Supplementary File S3). This highlighted that these structural variants often regarded as transient may persist across environments (from sourdough to the human host) and throughout decades, with up to 28 years spanning between the collection of isolates in question (Supplementary Table TS2). Stable non-hybrid triploid and triploid-aneuploid groups, like the ‘f’ subclade, have very rarely been reported before, and interestingly, the already described examples are among Mantou clades, also from the bakery environment.

### Lack of niche expansion in Mantou clades

The fine-scale phylogenetic relationships in the Mantou clades, additional samples related to these, and in the Mixed origin clade, revealed patterns indicating a recent colonization of novel niches. Only two commercial baker’s yeast strains were found to cluster with Mantou isolates (namely, AEI, ANGGW.1). An additional three commercial strains were found in the Huangjiu/Asian fermentation clade, of which two were found in products from Europe (Figure 3; Supplementary Table TS2). Human isolates were very rare among the Mantou sourdough clade yeasts, despite the fact that it was shown that sourdough yeasts are capable of passing through the intestinal tract in a viable form (Perricone *et al*. 2014). It appears that this doesn’t lead to colonization.

### Mixed origin yeasts colonize sourdough and the human host worldwide with clonal populations

In contrast to Mantou sourdough yeasts, widespread niche expansion was observed in most of the subclades of the Mixed origin yeasts. In line with previous observations (Bigey *et al*. 2021) these commercial baker’s yeasts often colonized the sourdough environment, predominantly in Europe (Figure 3, Supplementary Table TS2). That included the bakery sourdough samples in our study and the traditional Transylvanian isolate as well, which didn’t harbor true sourdough *Saccharomyces*. Ethnographic accounts exist on the rich and unique traditions and preparation methods for sourdough in previous centuries in Hungary and in ethnic Hungarian regions in the Carpathian Basin (Dénes *et al*. 2012; Tömösközi *et al*. 2023), yet, we did not find local, true sourdough yeasts, suggesting that they may have gone extinct in the country.

Niche expansion into the human host was also prevalent among the subclades of the Mixed origin yeasts. Apparently, both the human body and the sourdough environment is under recurrent colonization by various clonal clusters. During this colonization, meiotic reproduction does not play a detectable role. The clonal clusters may display diversity in genome structure variants, but at the same time, they also retain their polyploidy and certain hallmark aneuploidies and heterozygous rearrangements in a remarkably stable form. These observations reveal a more nuanced and complex picture on the evolutionary advantage of phenomena like meiotic reproduction, high genomic instability, and high mutation rate proposed earlier for *Saccharomyces* colonizing the high-stress environment of the mammalian host (Grimberg and Zeyl 2005; Zhu *et al*. 2016; Raghavan *et al*. 2018, 2019).

### Summary

Our study reveals the widespread niche expansion and strain redundancy of baker’s yeast, one of the most commercially important groups of *Saccharomyces*. The worldwide success of these yeasts apparently relies on a few, mostly tetraploid lineages propagated by multiple companies. These lineages possess their own unique gross chromosomal rearrangements, often stable aneuploidies, and at the same time show a wide range of genetic variability both in the baking environment (bakery industry and traditional sourdoughs) and upon the colonization of the human host.

## Data availability

Raw genome sequencing files are uploaded to NCBI GenBank (BioProject PRJNA1218442). Metabarcoding analysis and data on yeast species in sourdoughs is uploaded to FigShare (doihttps://doi/: 10.6084/m9.figshare.28193456 and doi: 10.6084/m9.figshare.28193495). Called cohort vcf, dendrogram and network files, raw data for various analyses on genomes, and values used to create heatmaps are uploaded to FigShare as a single project (https://figshare.com/projects/Baker_s_and_sourdough_yeasts_from_Hungary/233972), along with files with searchable text for the phylogenomic dendrogram and for the figures detailing CCNVs and GCRs.

## Acknowledgements

The research was supported by the NRDI Fund of Hungary grant FK 138910 [to W.P.P.]. The project was also supported by the Higher Education Institutional Excellence Program (NKFIH-1150-6/2019) of the Ministry of Innovation and Technology in Hungary, within the framework of the Biotechnology thematic program of the University of Debrecen. This project has received funding from the HUN-REN Hungarian Research Network. Part of this work (study of meiosis) was funded by the Austria-Hungary Action (AÖU) program (105ÖU3). Personal funding was received from the New National Excellence Program of the Ministry for Innovation [ÚNKP-22-5-DE-409 to W.P.P.; ÚNKP-22-3-II-DE-187 to H.V.R.]; and the János Bolyai Research Scholarship of the Hungarian Academy of Sciences [BO/00227/20/8 to W.P.P.]. Sz.P. was supported by the project TKP2021-NKTA-34, which has been implemented with the support provided from NRDI Fund under TKP funding scheme. We are very grateful to Guadalupe Pinar (BOKU, Vienna) for her always selfless and useful help during AÖU projects.

## Author Contributions

Conceptualization W.P.P., I.P., K.L., E.M., R.S.; methodology W.P.P., H.V.R., B.N., D.B., A.I., K.L., Sz.P.; formal analysis W.P.P., H.V.R.; A.I., B.N., A.H., D.B., Zs.A., L.H., R.B.; resources W.P.P., K.L., I.P., R.S., E.M., Á.F., Sz.P.; data curation H.V.R., A.I., B.N., W.P.P.; writing – original draft preparation W.P.P., H.V.R.; writing – review and editing W.P.P.; visualization, W.P.P., H.V.R., R.B., B.N.; supervision W.P.P., K.L., R.S.; project administration W.P.P., K.L., E.M.; funding acquisition, W.P.P., I.P., E.M., K.L.

## Supplementary data

Supplementary File S1. Supplementary methods and validation of bioinformatics pipelines.

Supplementary File S2. Detailed data on clonal clusters.

Supplementary File S3. Coverage and allele ratio plots.

Supplementary File S4. Gross chromosomal rearrangements.

Supplementary File S5. Companies and brands of yeasts in the baking industry.

Supplementary File S6. Sporulation and spore viability data.

Supplementary Table S1. Genomes and their metadata used in this study.

Supplementary Table S2. All yeast samples according to phylogeny, with additional data.

