## Supplementary File S1 for "Commercial *Saccharomyces cerevisiae* baker’s yeasts: strain redundancy, genome plasticity, and colonization of the sourdough environment and the human body"

**Supplementary File S1. Supplementary methods and validation of bioinformatics pipelines.**

**Phylogenomics.**

As described in the Materials and Methods section, phylogenomic networks were generated in this study both from the SNPs called from mapping files using GATK and from an alignment and assembly-free (AAF) approach. The former method involved creating an SNP matrix that by definition only contains SNP allele information and since vcf2phylip was used, it also captured heterozygosity in the form of heterozygous nucleotide codes (e.g. A and G = R). The matrix also contains spanning deletions, i.e. if a SNP at a locus is unknown due to a spanning deletion, the final matrix will contain a ‘*’ character instead of a SNP. However, if a genome is heterozygous at a given position, possessing one SNP allele and one spanning deletion simultaneously, the SNP matrix will simply contain an SNP allele as there are no IUPAC codes for such a combination of alleles, as described below:

|  | Genome I | Genome II |
| --- | --- | --- |
| Allele 1 | A | A |
| Allele 2 | G | * |
| Encoding in SNP matrix used for phylogeny | R | A |

Applying an AAF approach (Fan *et al.* 2015) circumvents this limitation by using k-mers, spanning *k* nucleotides from the sequencing reads. In the case of a genome heterozygous at a given position for a SNP and a spanning deletion, the AAF software will identify multiple different k-mers, ultimately capturing the heterozygosity and allowing its incorporation into the distance matrix of all genomes. This distance matrix then can be used to generate phylogenomic networks. This unique characteristic of the AAF is shortly illustrated below, with the SNP colored red, the neighboring nucleotides in the sequencing reads pink.

|  | Genome I | Genome II |
| --- | --- | --- |
| Allele 1 on a read | AGCTGAAGAGTTCACT | AGCTGAAGAGTTCACT |
| Allele 2 on a read | AGCTGAAGGGTTCACT | AGCTGAAGGTTCACT |
| Identified k-mers | AGCTGAAGA  AGCTGAAGG  GCTGAAGAG  GCTGAAGGG  CTGAAGAGT  CTGAAGGGT  TGAAGAGTT  TGAAGGGTT  etc. | AGCTGAAGA  AGCTGAAGG  GCTGAAGAG  GCTGAAGGT  CTGAAGAGT  CTGAAGGTT  TGAAGAGTT  TGAAGGTTC  etc. |

Furthermore, k-mers are derived from reads that represent a region on physical chromosomal copy and so, they capture local haplotypes if two variant positions are sufficiently close (and covered by a single k-mer). This is not the case for called alleles that are exported to haplotype matrices, as any haplotype information is lost in such matrices. This is exemplified in the below table on the example of two hypothetical tetraploid heterozygous genomes that only differ in their haplotypes and not in alleles they possess. Variants are colored in red. The blow example shows SNPs, but as shown above, k-mers can efficiently capture haplotypes with short indels as well.

|  | Genome I with 2 different haplotypes | Genome II with 3 different haplotypes |
| --- | --- | --- |
| chromosome copy 1 | ATATGCAAAATCAA | ATATGCAAAATCAA |
| chromosome copy 2 | ATATGCAAAATCAA | ATATGCAATATCAA |
| chromosome copy 3 | ATATCCAATATCAA | ATATCCAATATCAA |
| chromosome copy 4 | ATATCCAATATCAA | ATATCCAATATCAA |
| Identified k-mers that contain both variant positions | ATATGCAAA  TATGCAAAA  ATGCAAAAT  TGCAAAATC  GCAAAATCA  ATATCCAAT  TATCCAATA  ATCCAATAT  TCCAATATC  CCAATATCA | ATATGCAAA  TATGCAAAA  ATGCAAAAT  TGCAAAATC  GCAAAATCA  ATATGCAAT  TATGCAATA  ATGCAATAT  TGCAATATC  GCAATATCA  ATATCCAAT  TATCCAATA  ATCCAATAT  TCCAATATC  CCAATATCA |
| Called alleles when allele calling is applied | G or C; A or T | G or C; A or T |

The AAF method can be used to determine the total number of k-mers and shared k-mers between genomes, a valuable information to supplement called allele-based comparisons. This information can be used to infer whether two genomes have similar ploidy, or whether two polyploid heterozygous genomes are almost identical even when haplotype-aware k-mers are compared. The AAF analysis can also be used to calculate a similarity value between genomes. As described in Fan et al. (2015), the distance value (D) for AAF-based comparisons with the software applied here is calculated as follows:

$$D=-\frac{1}{k}\log\frac{ns}{nt}$$

where *k* is the k-mer length, *ns* is the number of shared k-mers between two compared genomes and *nt* is the total number of k-mers in the samples with smaller total k-mer value.

To obtain similarity values from the calculated distances, we used a simple equation for similarity (S):

$$S=1-D$$

Considering these calculation methods, we can apply the above equations to show how the AAF phylogenomics may handle the peculiarities of tetraploid genomes. In the case of a hypothetical highly heterozygous tetraploid and a diploid genome, the diploid would most probably have fewer k-mers and hence would determine the *nt* value. In a hypothetical case of the tetraploid containing all the alleles of the given diploid, the equations would give a similarity of 1 between these two genomes, regardless of what additional variants the tetraploid has that are missing from the diploid in question. On the other hand, when two heterozygous polyploids with similarly high total k-mer numbers are compared, they have to share almost all of their alleles including the heterozygosities and the local haplotypes to have a similarity value close to 1. Thus, similarity is calculated in a manner that emphasizes shared variants when low-heterozygosity (*e.g.* diploid) and high-heterozygosity (especially polyploid) genomes are compared to each other, while being more stringent when to highly heterozygous genomes are assessed.

In summary, the AAF analysis adapted here can take SNP and indel mutations and local haplotypes (that do not exceed the k-mer length) even in heterozygous form into account and is very sensitive to differences among tetraploid genomes (*e.g.* haplotype-level differences), more so than to differences among diploids. It gives information on total number of k-mers in a genome, and the amount of shared k-mers between pairs of genomes. It can be considered an addition to comparisons based on called SNPs that is the standard in yeast comparative and phylogenomics. Based on the above described distance values, a phylogenomic network can also be created (*e.g.* in SplitsTree) that will emphasize shared variants between genomes with low and high heterozygosity, and emphasize differences when genomes of similar heterozygosity and k-mer number are compared. Such a network can be used as an addition to networks and phylogenies based on SNP matrices.

**Comparative genomics.**

Illumina read mapping approach. As described in Materials and Methods, reads were first mapped to a concatenated reference of eight *Saccharomyces* species. Coverage was assessed in 10 kb windows sliding every 5 kb. Coverage data was corrected for ploidy. Coverage was visualized in the form of bar charts at every sliding window position. This allowed for a visual identification of hybrids and introgressed regions. In the former case, whole chromosomes have relatively even coverage, while in the latter, peaks of coverage are visible in a specific region of the donor species’ reference (and depleted coverage at the corresponding region of the *cerevisiae* reference).

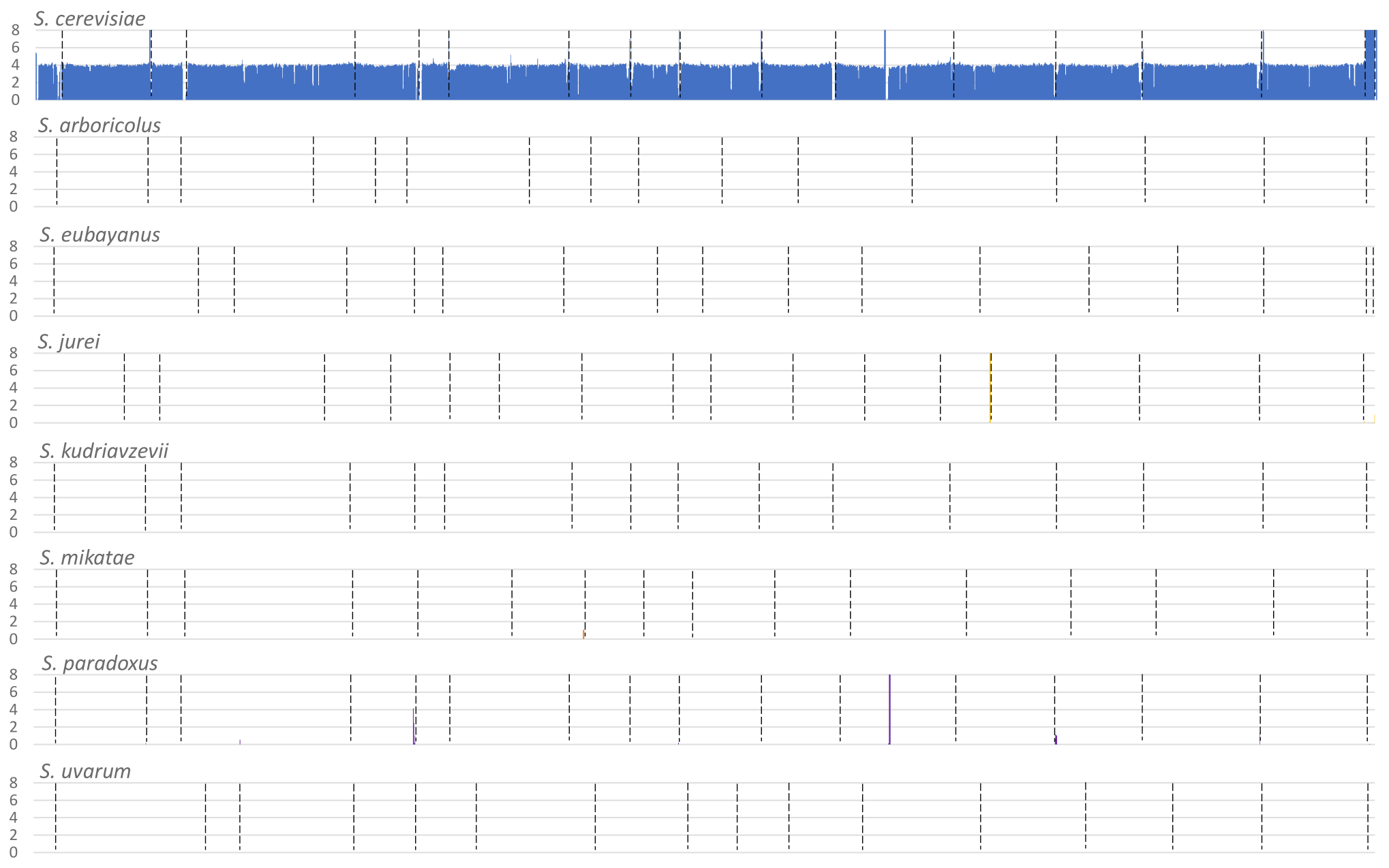

**Above: Example of a coverage mapping to an 8-species reference in the case of a non-hybrid baker’s yeast (BY0017). For each species, the *y* axes show copy number calculated from coverage of a given sliding window, the *x* axes show chromosomal positions, with chromosomes separated by dashed lines.**

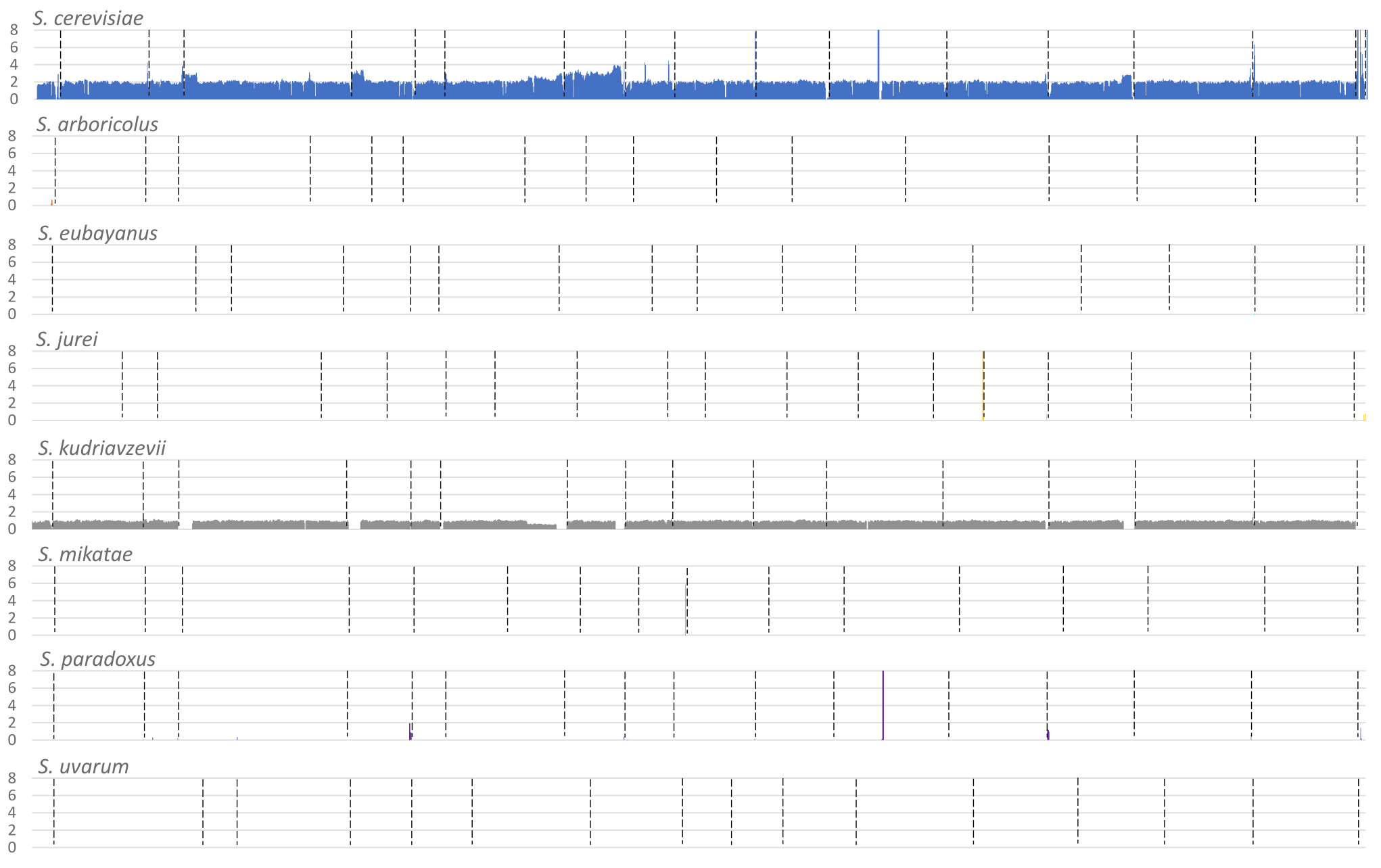

**Above: Example of coverage mapping to an eight-species reference in the case of a *S. cerevisiae* × *S. kudriavzevii* hybrid baker’s yeast (BR005). Note the coverage for the *S. cerevisiae* (in blue) and *S. kudriavzevii* (in grey) subgenomes.**

In a subsequent mapping to only the S288c reference or to a two-species reference in hybrids, coverage was not affected by introgressions, this was used to identify large deletions and duplications.

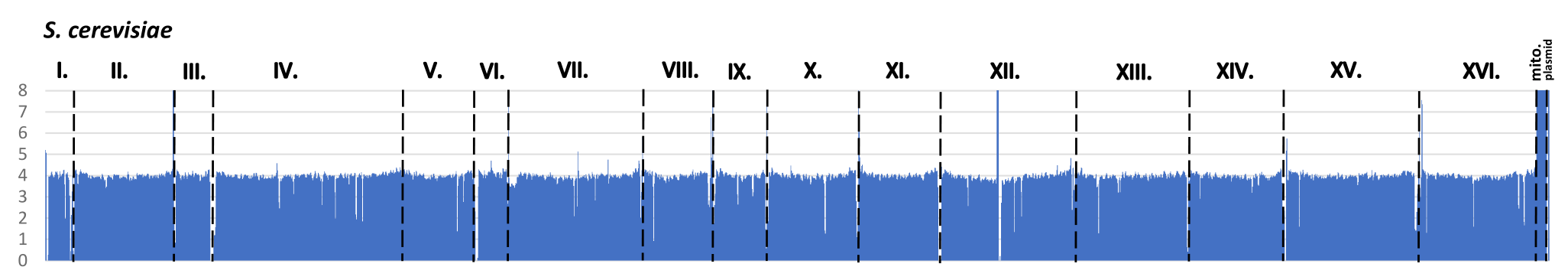

**Above: Example of a coverage mapping to a single-species reference in the case of a non-hybrid baker’s yeast.**

Finally, sudden changes in coverage graphs were manually examined in IGV (Thorvaldsdottir *et al.* 2013) to determine locations of large deletions and duplications. This was needed because the above-described coverage analysis used sliding windows and thus smoothed out the exact location of these structural variations. Here, we used BEDTools 2.30.0 (Quinlan and Hall 2010) to create per-base BedGraph files (first, per-base coverage files were created with the genomecov option, then an extra column was added as start position in order to convert the coverage file to a BedGraph). Per-base BedGraphs can be loaded into IGV and dynamically zoomed in, thereby allowing to assess large-scale structural variants that were first broadly identified by the 10 kb sliding window coverage analysis. Per-base BedGraphs were viewed together with the .bam files of the given genomes in various resolutions, enabling the identification of chromosome break-points. Possible breakpoints were compared in multiple genomes in a single IGV window that had the given rearrangement and also ones that did not have it, making the identification of breakpoints more reliable. Regions before and after breakpoints were also compared on the allele plots (see below). Since the mapping was performed onto the reference genome or genomes, these breakpoints always refer to the S288c reference genome. The breakpoints were further investigated by viewing their loci in the *Saccharomyces* Genome Database (Engel *et al.* 2021). Genes, LTRs, ARSs and also recombination hotspots (Pan *et al.* 2011) at or near the identified breakpoint were listed. An example of this fine-scale coverage analysis is shown below.

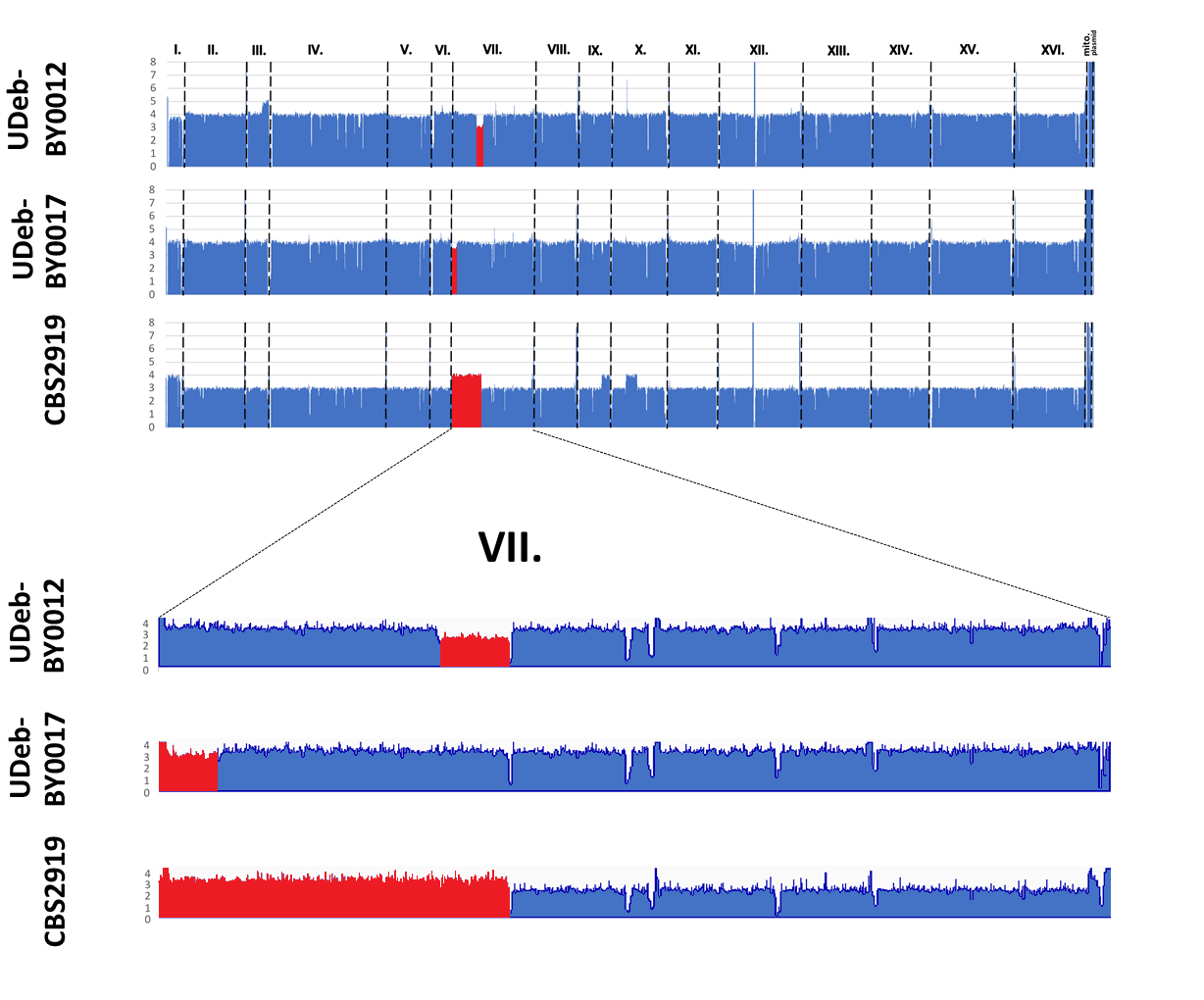

**Above: Example of the coverage-based identification of three different gross chromosomal rearrangements (GCRs) affecting chr. VII in three genomes. On top, two deletions (UDeb-BY0012: interstitial deletion; UDeb-BY0017: terminal deletion) and a terminal duplication on the left chromosomal arm (CBS2929) are shown. These are highlighted in red on the 10 kb sliding window coverage graphs. On the bottom, BedGraph coverage data is shown, exported from IGV. The GCR regions are also highlighted in red. The breakpoints were identified by further zooming in, and it was confirmed that the breakpoints for the deletion in the UDeb-BY0012 genome and for the duplication in the CBS2919 are at the same locus.**

To assess the ploidy of the genomes, we used an approach based on allele depth ratios. In a diploid heterozygote, alleles at a given variant site will have an approximately similar coverage (this can vary more in low-coverage regions), while at a heterozygous variant site on a trisomic chromosome, 1:2 and 2:1 ratios are to be expected. In a tetrasomic chromosome, heterozygous variant sites will have 1:3 and 1:1 allele depth ratios, etc. By exporting allele depth values and by plotting them onto the reference, we visually determined chromosome copy numbers and used this to correct coverage plots for ploidy. Coverage and allele ratios were manually cross-compared especially in non-diploids and aneuploids.

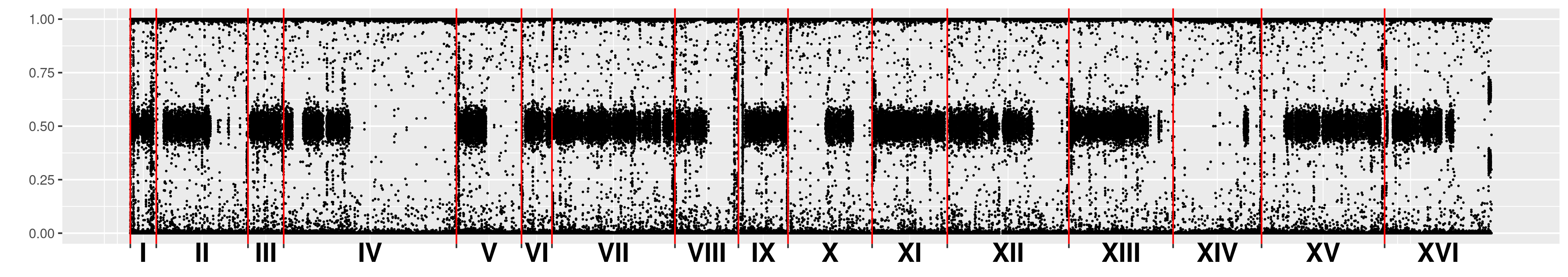

**Above: Example of an allele (depth) ratio plot in a diploid (strain BTC). The y axis shows allele depth ratios for individual called alleles, the x axis shows chromosomal positions, chromosomes are separated by red lines. Note the prevalence of heterozygous variants with approx. 50-50% allele ratios along with homozygous regions with approx. 0-100% ratios.**

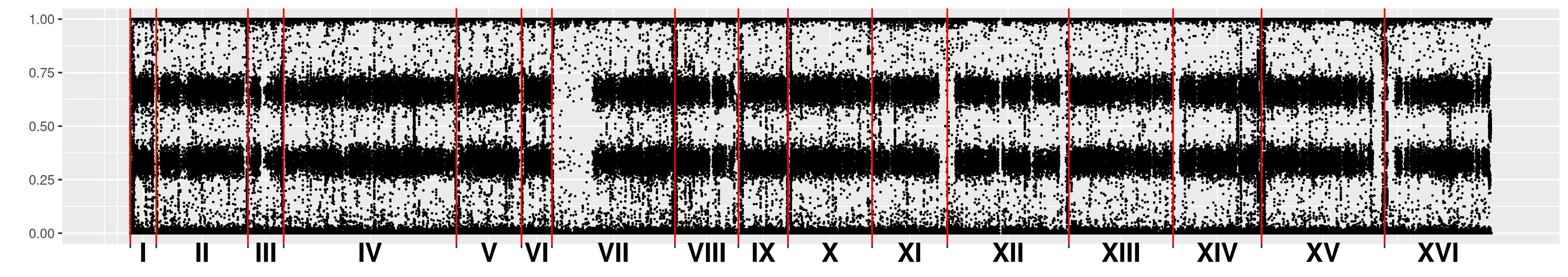

**Above: Example of an allele ratio plot in a triploid (strain BTA), note the prevalence of heterozygous variants with approx. 33-66% allele depth ratios.**

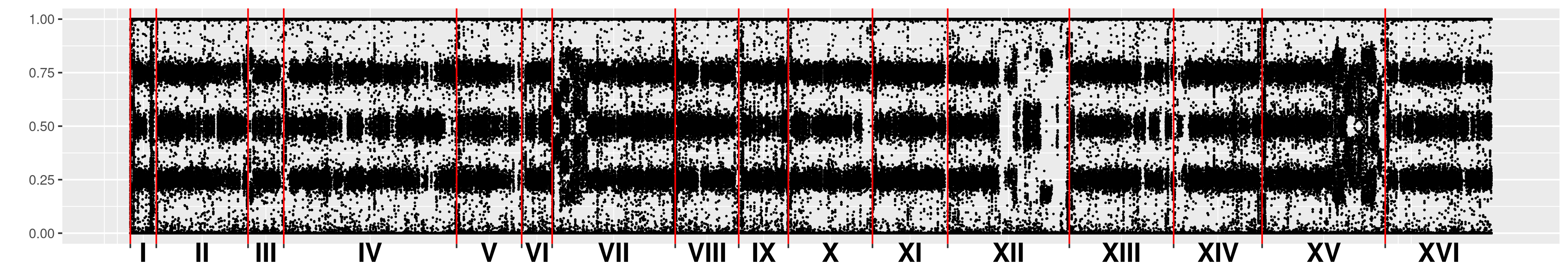

**Above: Example of an allele ratio plot in a tetraploid (BY0017), note the prevalence of heterozygous variants with approx. 50-50% and approx. 25-75% allele depth ratios.**

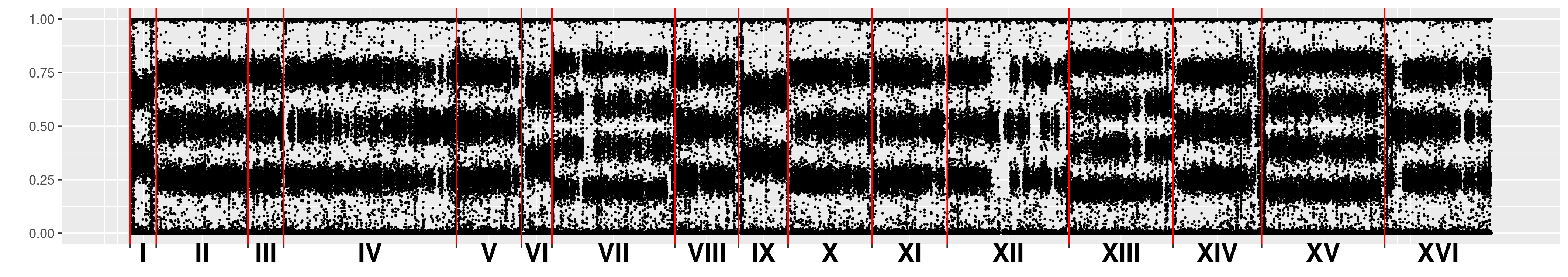

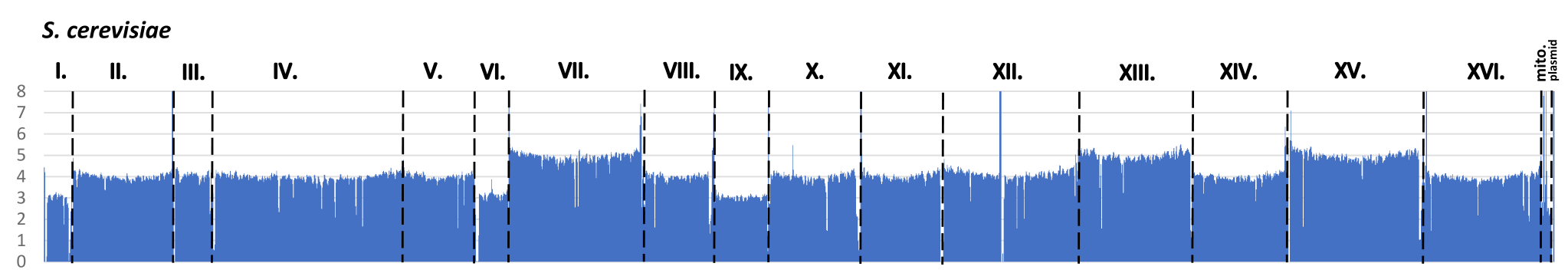

**Above: Example of an allele ratio plot and coverage plot in a tetraploid aneuploid (strain BMM), note change in allele ratios co-occurring with coverage (and thus copy number) changes along the chromosomes.**

It must be noted that homozygous non-haploid genomes can not be properly assessed by this coverage mapping and allele ratio plotting method. In such rare cases affecting only two genomes, we used data from Peter et al. (2018) who used total DNA content measurement. In a few cases of genomes, the coverage analysis showed highly fluctuating coverage across chromosomes, probably a result of sonication in the library preparation step. However, in these cases the allele ratios were sufficient to determine chromosome copy numbers. In most cases, we did not attempt to delineate deletions and duplications in these genomes based on changes in coverage.

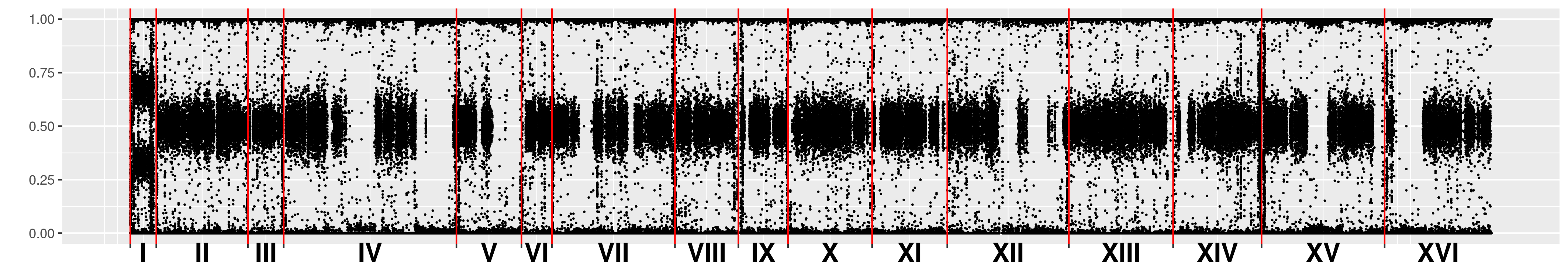

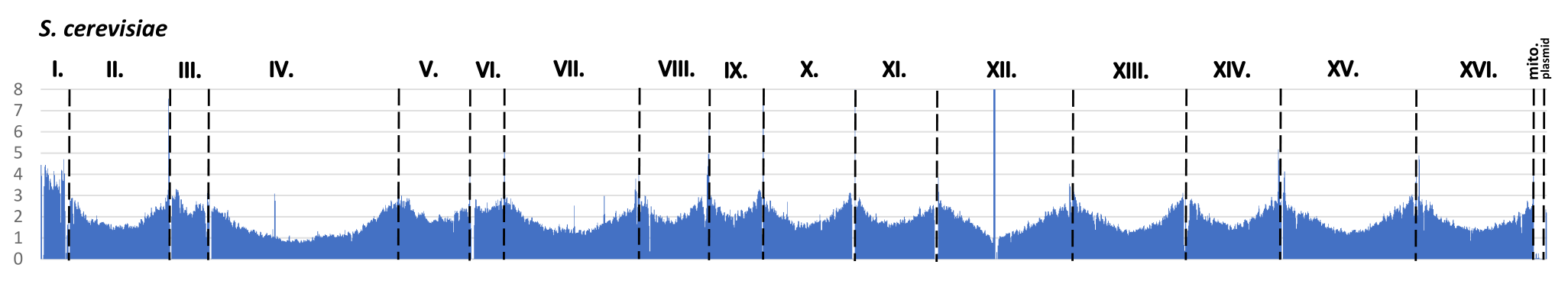

**Above: Example of an allele ratio plot and coverage plot in a diploid genome severely affected by coverage fluctuations (MUCL52901), note the fluctuating coverage and the still consistent allele ratios.**

**Confirming genome structure variations by long-read phasing and limitations.**

To confirm that large-scale genome structure variations can be identified by mapping short-read data to the S288c reference genome and analyzing this for coverage and allele ratios, we applied Oxford Nanopore sequencing, and produced an assembly with short-read polishing for the tetraploid UDeb-BY0012 isolate. The software nPhase mapped long reads to this reference and built haplotigs by forming clusters from long reads. Clusters were used to produce consensus sequences, which were compared across clusters and identical ones were merged. The result of this is a number of non-mergeable, color-coded haplotigs represented on a graph with subgraphs by the software. Each subgraph displays the predicted haplotigs for a different chromosome, each predicted haplotig is on a different row on the y axis, and the x axis displays the position along the chromosome. Haplotigs may cover larger portions of a chromosome and thereby approach representing true, physical haplotypes of the genome, or may be shorter if the software was unable to connect a haplotig at one locus with nearby loci. If the software determined similar, yet non-mergeable haplotigs at a given locus, these will be separated. Thus, more than four delineated haplotigs may represent four true haplotypes in a tetraploid if these could not be merged properly due to bioinformatics constraints. It is noted that a tetraploid genome may have less than four haplotypes, if two or more chromosomal copies are identical. A tetrasomic chromosome showing exclusively 50-50% allele depth ratios in short-read based analysis is, for example, expected to have two haplotypes, ideally represented by two haplotigs when long-read based phasing is applied.

In the case described below, the yeast UDeb-BY0012’s tetraploid heterozygous genome is characterized by a terminal duplication on the right arm of chr. III an interstitial deletion on chromosome VII affecting three of four chromosomal copies in both cases. Long-read phasing clearly identified an increase from three to four haplotigs at the locus of the terminal duplication and a drop in the number of haplotigs from four to three at the locus of the deletion. The location of these changes corresponded to the coverage and allele ratio changes obtained by short-read based analysis, thereby the long-read based approach confirmed the results of the short-read based analyses for this genomic regions, as shown below.

**Below: Short-read-based analysis compared to long-read phasing in the case of the UDeb-BY0012 genome. Top: short-read based coverage plot as in the above examples, *y* axis is corrected for copy number. Middle: allele depth ratio plots, as in the above examples. Bottom: haplotigs phased by nPhase (in groups of four chromosomes). The relatively short chr. I and the longer chr. II. were almost completely phased by the nPhase algorithm. Chr. III shows a characteristic terminal duplication on the right arm of chr. III. visible on coverage and allele ratio plots, and at the same locus, the number of haplotigs identified by nPhase increases (marked with red arrowhead). However, nPhase did not identify the expected four haplotigs on the left arm (this can occur if two of four chromosomal copies are almost identical). The software could not phase most of the remaining chromosomes entirely, but produced haplotigs of mostly several hundred kilobases. The characteristic interstitial deletion on chrom. VII identified using short-read data was visible as a local drop from four to three haplotigs (marked by red arrowhead). In the case of chr. XII, the right arm showed a characteristic change in the allele depth ratio plot, with ~50-50% ratios dominating. The same region was only phased to two haplotigs by nPhase (marked by red arrowhead).**

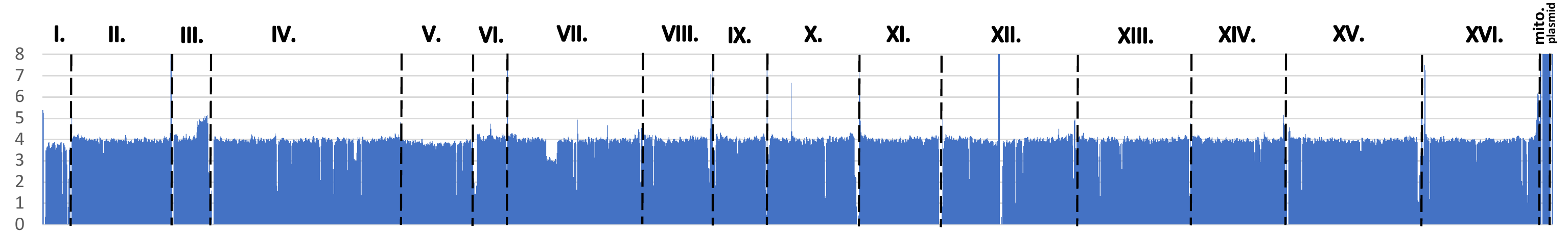

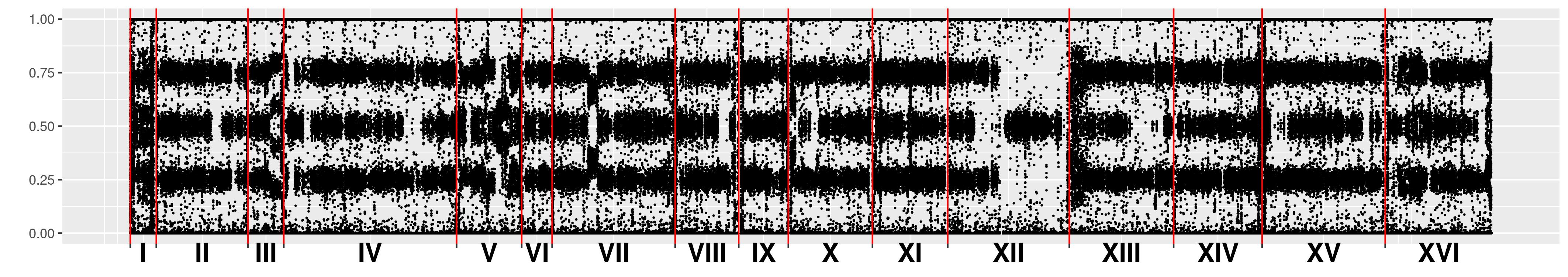

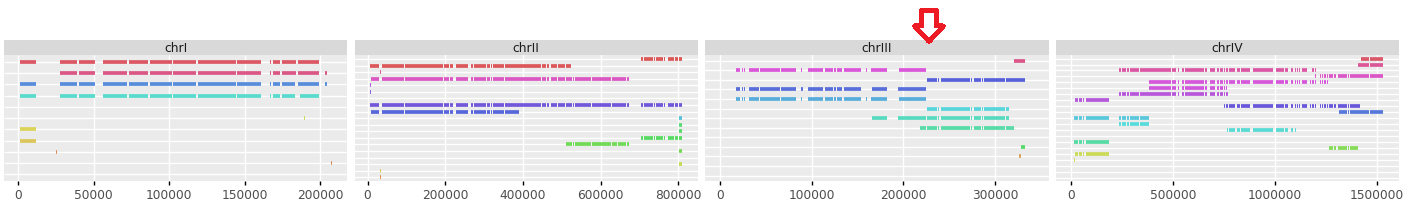

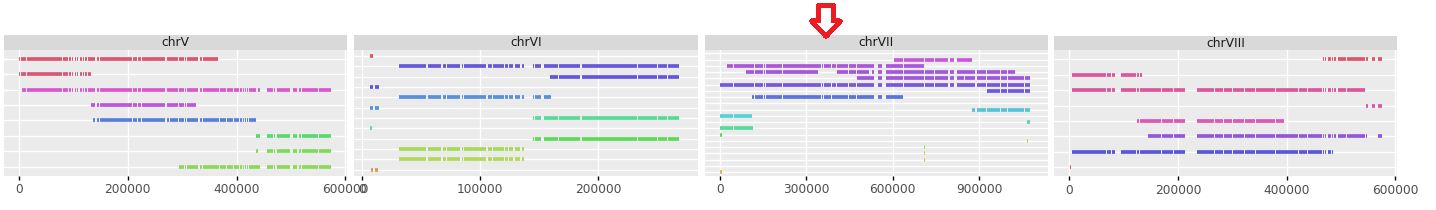

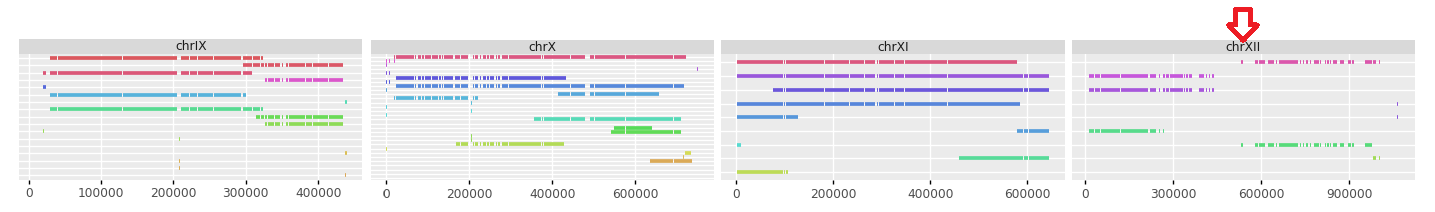

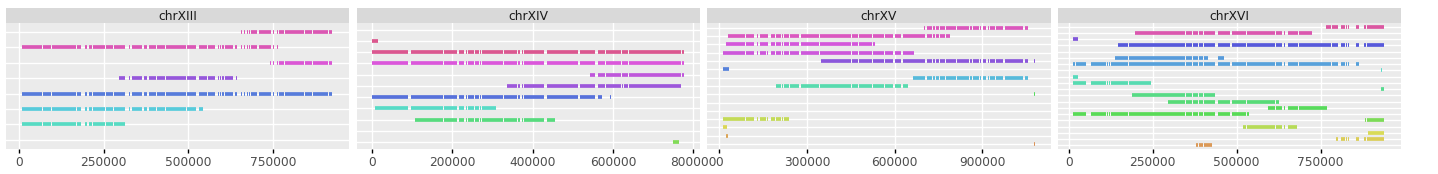

**Below: alignment of the UDeb-BY0012 haploid assembly against the S288c reference genome, showing high synteny.**

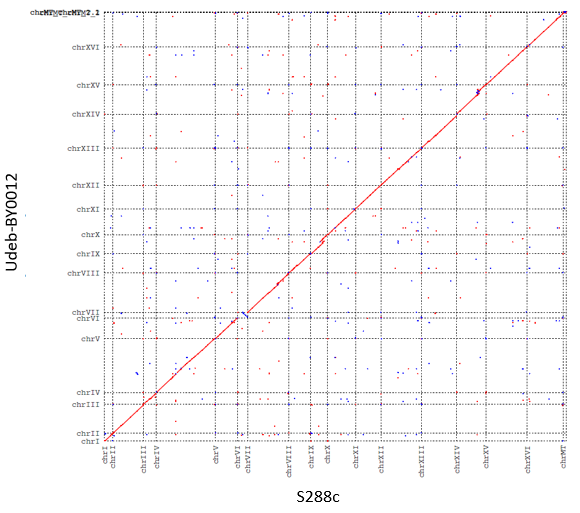

Finally, it is noted that the analyses described here are not suitable to identify inversions or reciprocal translocations, especially if they are in a heterozygous form *i.e.* affecting only one or two copies in a tetraploid. A mapping-based approach is used both in the case of short-reads and in the case of the nPhase software. Mapping the read coverage or allele ratio changes from chromosomes affected by these structural variations would in general be indistinguishable from those of non-affected chromosomes. For example, a hypothetical reciprocal translocation in a tetraploid genome between a single copy of chr. IV and a single copy of chr. XV would result in altogether two novel chromosomes with translocated regions and six unchanged chromosomes (when only chr. IV and XV are considered). During library preparation and short-read sequencing, however, the amount of reads for any given region of the chromosomes would be unchanged. During mapping, reads would be mapped to the reference genome and the translocation would not be detected. In the case of long-reads, the reads spanning the translocated region would not map correctly and would not be merged to haplotigs.

However, reciprocal translocations between non-homologous chromosomes would affect the physical sizes of chromosomes and may thus result in novel bands in karyotyping. The intensity of such bands can be expected to depend on copy numbers of the chromosomal variants they represent in a karyotyping gel.

**Yeast species identification in sourdough.**

We assessed the microbial diversity of sourdoughs obtained for this study. Non-*Saccharomyces* yeasts from sourdoughs were identified by Sanger sequencing: the GoTaq G2 polymerase (Promega, Madison, WI, USA) was used to amplify the variable region of the 26S ribosomal large subunit with primers NL1 (GCATATCAATAAGCGGAGGAAAAG) and NL4 (GGTCCGTGTTTCAAGACGG). The PCR products were subjected to capillary sequencing after PCR cleanup with the E.Z.N.A. cycle pure kit (Omega Bio-Tek, Norcross, GA, USA) by the sequencing core facility of the University of Debrecen. Sequenograms were manually inspected in Chromas (Technelysium, Brisbane, Australia) and the NCBI BLAST service was used for species identification, whereby the species with the closest hit in the NCBI GenBank was considered as a putative species identification, and then the sequence of the hit species’ type strain was once again aligned to the query sequence using BLAST. A similarity of >99% with the type was considered a definitive species identification. Sequences were deposited in GenBank. Yeast colonies and cells were photographed as uploaded to FigShare (doi: 10.6084/m9.figshare.28193495)

| **Species** | **Sourdough sample** | **GenBank accession** |
| --- | --- | --- |
| *Pichia kudriavzevii* | Sourdough 1 | PV077634 |
| *Pichia membranifaciens* | Sourdough 1 | PV077635 |
| *Pichia membranifaciens* (rough morphology) | Sourdough 1 | PV077636 |
| *Maudiozyma humilis* | Sourdough 7 | PV077637 |
| *Maudiozyma humilis* | Sourdough 8 | PV077638 |

**GenBank sequence IDs for yeast LSU data**

**Bacterial long-read 16S metabarcoding of sourdough samples.**

Total DNA from the sourdough samples was extracted by the Macherey-Nagel (Düren, Germany) Genomic DNA From Food kit following the manufacturers’ protocols. 16S metabarcoding was carried out using the 16S long-read metabarcoding kit (SQK-16S024) of Oxford Nanopore Technologies (Oxford, UK) according to the manufacturer’s instructions. In the first step of the library preparation a PCR was performed for the amplification of the 16S rRNA gene target region (~1500 bp) and to add a unique barcode to each sample. DNA concentrations were quantified by Qubit™ fluorometer (Invitrogen, Waltham, MA). The initial DNA concentration was 10 ng per sample, the final PCR mix contained 25 µl LongAmp™ Hot Start Taq 2× Master Mix, 10 µl input DNA, 10 µl 16S barcode primers (each) and 5 µl nuclease free water. The reaction was performed using a thermal cycler (Biometra TAdvanced, Analytik Jena, Jena, Germany) with the following PCR conditions: initial denaturation at 95°C for 1 min, 25 cycles of 95°C for 20 s, 55°C for 30 s, 65°C for 2 min, followed by a final extension step at 65°C for 5 min. The amplicons were purified with AMPure XP beads (Beckman Coulter, Brea, CA, USA) and eluted in 10 mM Tris-HCl pH 8.0 with 50 mM NaCl. DNA concentration of the samples was quantified with a Nabi spectrophotometer (MicroDigital, Seongnam-si, South Korea). Approximately 100 fmol of the combined library was loaded into an ONT SpotON flow cell. Super accurate basecalling and de-barcoding in the Guppy v6.4 software were performed after the sequencing, and reads were identified to species level by using the software Emu (Curry *et al.* 2022) (this is the default database of the software, a combination of rrnDB v5.6 and NCBI 16S RefSeq from 17 September, 2020), along with abundance data. The sequencing files for metabarcoding are deposited under the BioProject number PRJNA1220648. Data processing and the visualization of the results were carried out using the web-based platform MicrobiomeAnalyst (Chong *et al.* 2020). First, a taxonomy file was created for the abundance data using Emu and species of the Lactobacilli were updated to current taxonomy. Stacked bar charts with abundance data were generated. Results were uploaded to FigShare (doi: 10.6084/m9.figshare.28193456).
