## Supplementary File S2 for "Commercial *Saccharomyces cerevisiae* baker’s yeasts: strain redundancy, genome plasticity, and colonization of the sourdough environment and the human body"

**Supplementary File S2. Detailed data on clonal clusters.**

AAF-similarity and SNP-similarity in the subclades of the Mixed origin clade is shown, the order of genomes corresponds to the order on the phylogenomic dendrogram (Figure 3 and 5). Clonal clusters are outlined in red on the heatmaps. Each subclade of the Mixed origin clade is shown on separate composite panels of heatmaps and mean values for similarity between members of the clonal cluster and other genomes of the subclade, and average similarity values within the clonal cluster. Heatmap color ranges are given for AAF and SNP similarity separately.

**aff. Mixed origin**

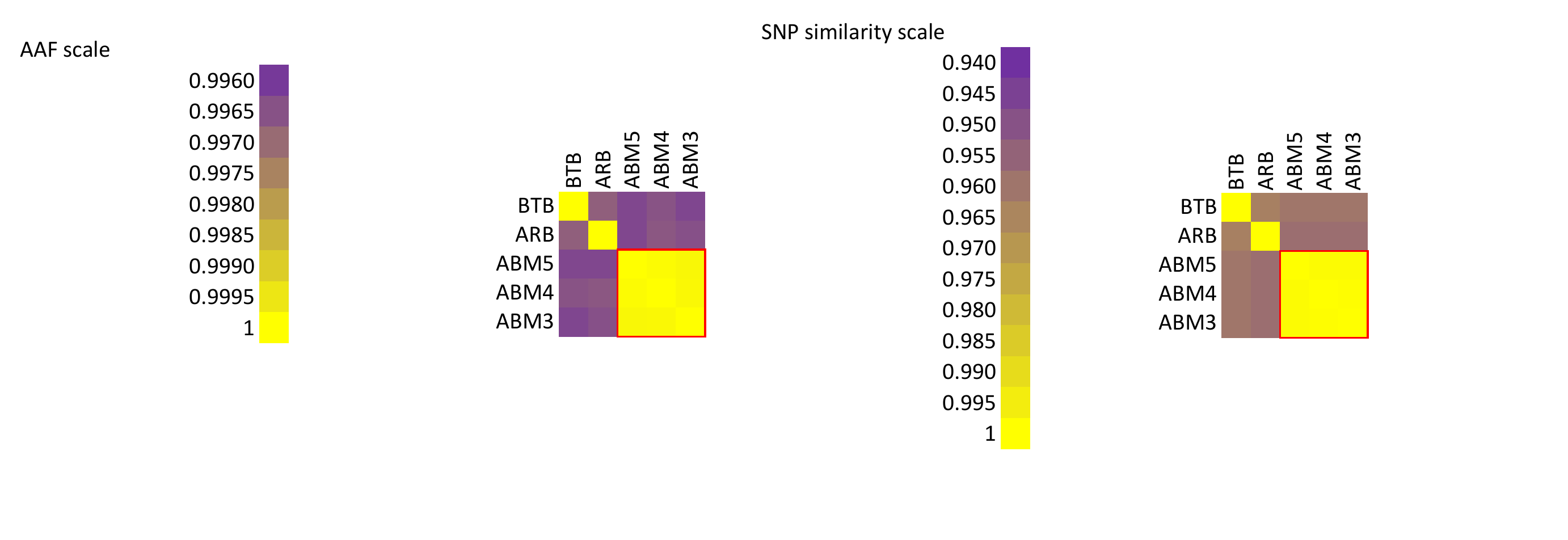

The genomes ABM5, ABM4, and ABM3 formed a clonal cluster.

|  | **AAF similarity** | **SNP similarity** |
| --- | --- | --- |
| **Clonal cluster vs. other genomes** | 0.9964 | 0.9592 |
| **Inside clonal cluster** | 0.9999 | 0.9995 |

**Mixed origin ’a’**

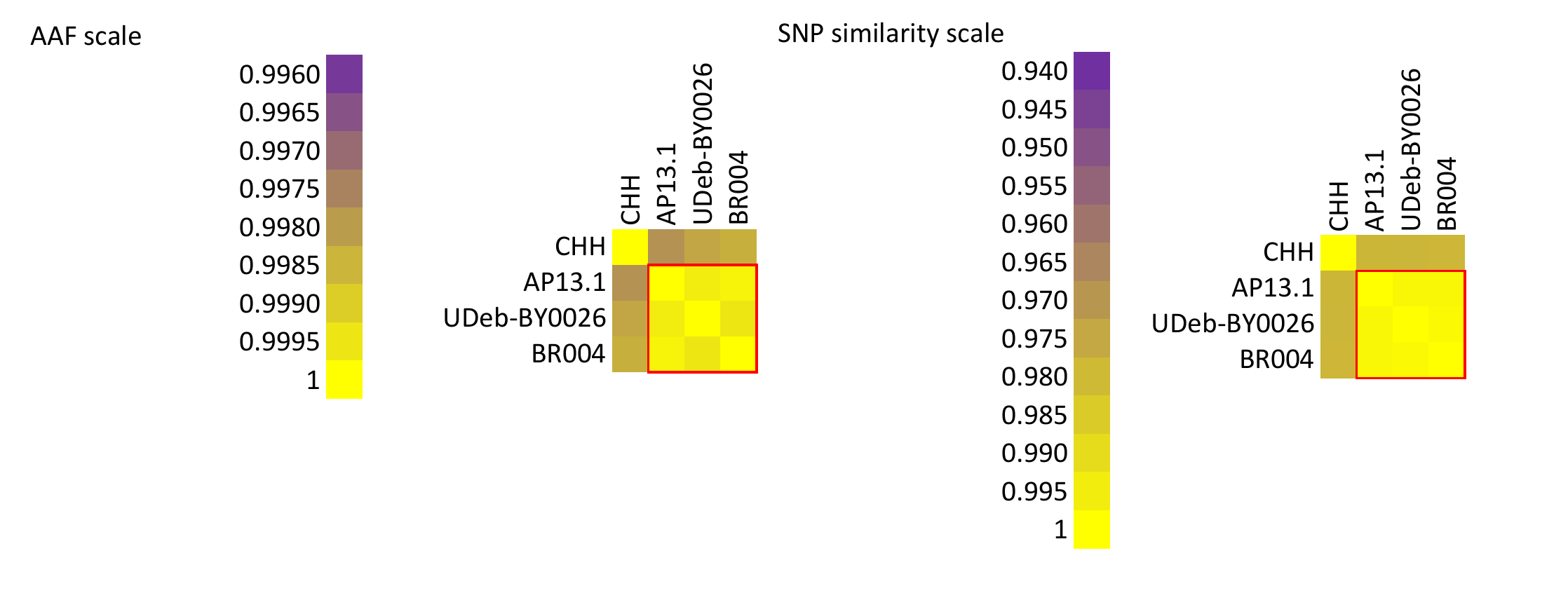

The genomes AP13.1, UDeb-BY0026, and BR004 formed a clonal cluster.

|  | **AAF similarity** | **SNP similarity** |
| --- | --- | --- |
| **Clonal cluster vs. other genomes** | 0.9981 | 0.9790 |
| **Inside clonal cluster** | 0.9998 | 0.9987 |

**Mixed origin ’b’**

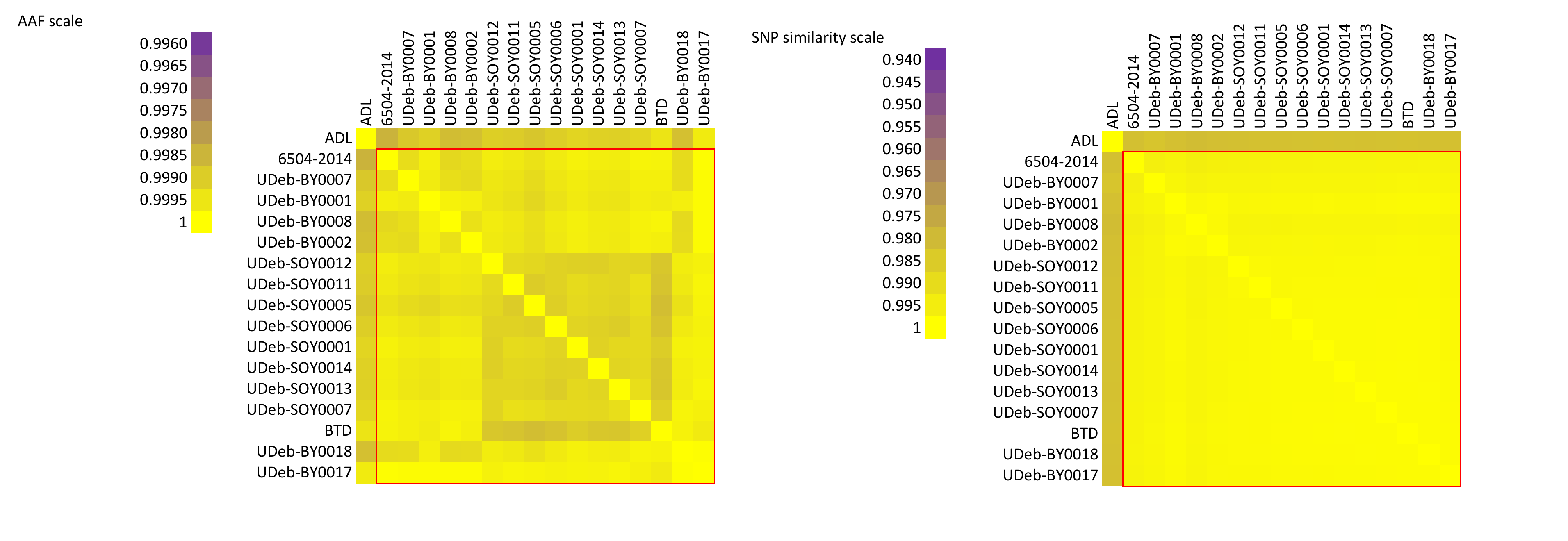

The genomes 6504-2014, UDeb-BY0007, UDeb-BY0001, UDeb-BY0008, UDeb-BY0002, UDeb-BY0018, UDeb-BY0017, UDeb-SOY0012, UDeb-SOY0011, UDeb-SOY0005, UDeb-SOY0006, UDeb-SOY0001, UDeb-SOY0014, UDeb-SOY0013, UDeb-SOY0007, and BTD formed a clonal cluster.

|  | **AAF similarity** | **SNP similarity** |
| --- | --- | --- |
| **Clonal cluster vs. other genomes** | 0.9990 | 0.9822 |
| **Inside clonal cluster** | 0.9995 | 0.9981 |

**Mixed origin ’c’**

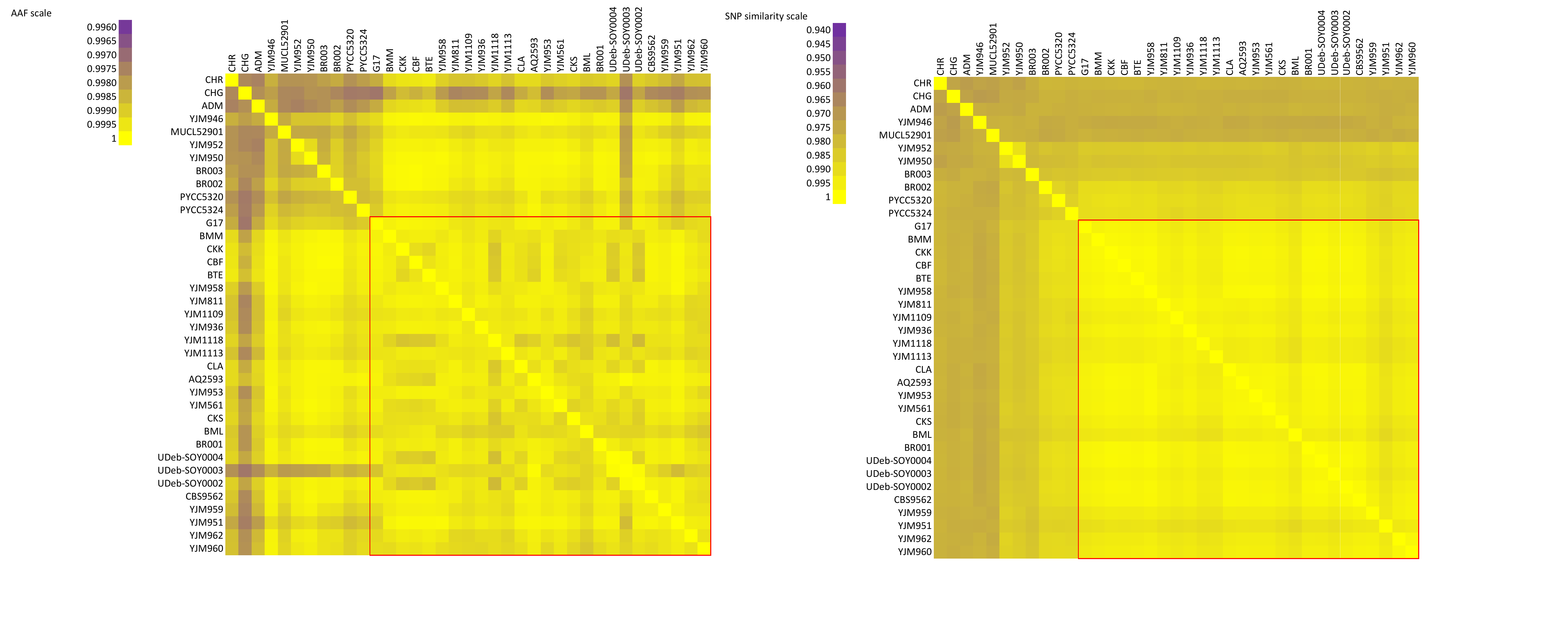

The genomes G17, BMM, CKK, CBF, BTE, CLA, AQ2593, CKS, BR001, UDeb-SOY0004, UDeb-SOY0003, UDeb-SOY0002, YJM958, CBS9562, YJM959, YJM951, YJM962, YJM960, YJM811, YJM1109, YJM936, YJM1118, YJM1113, YJM953, YJM561, and BML formed a clonal cluster.

|  | **AAF similarity** | **SNP similarity** |
| --- | --- | --- |
| **Clonal cluster vs. other genomes** | 0.9993 | 0.9826 |
| **Inside clonal cluster** | 0.9995 | 0.9957 |

**Mixed origin ’d’**

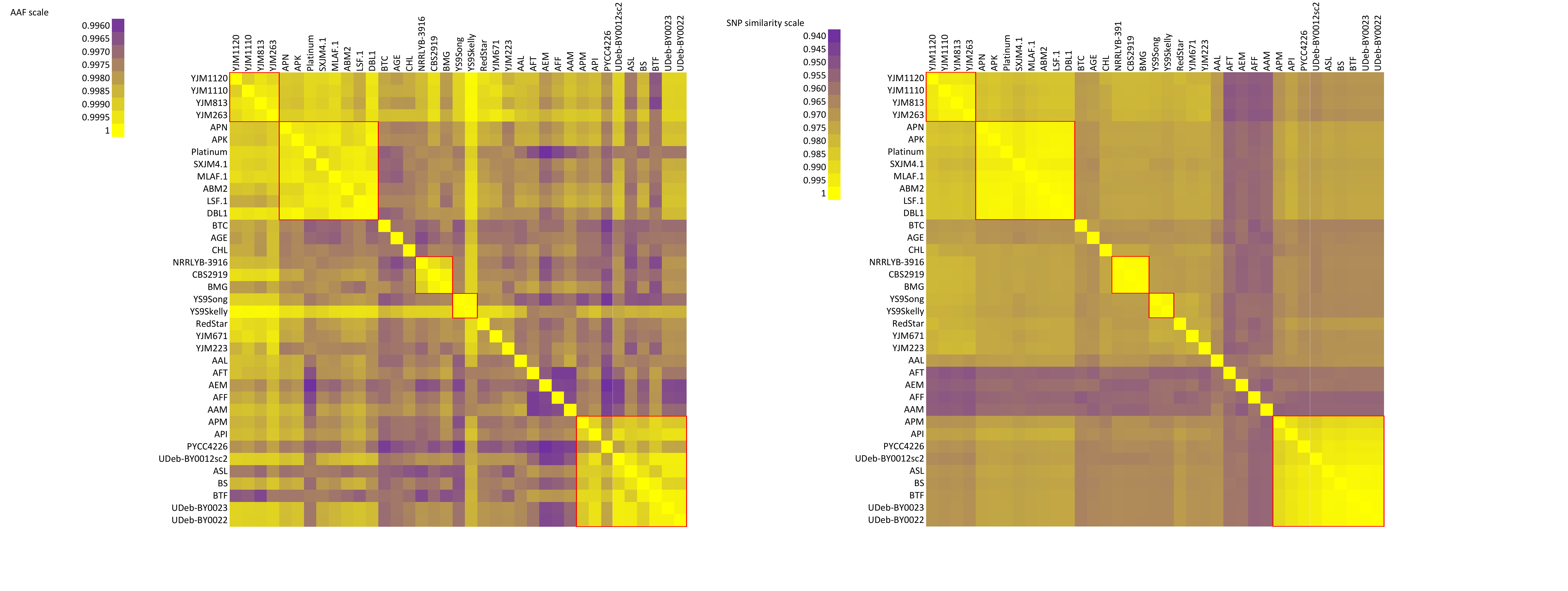

Note: AEM, AFF, and AAM are sporulated haploids thus they are removed from IBS2 panels. YS9Song and YS9Skelly are not evaluated due to low and highly fluctuating coverage.

The genomes YJM1120, YJM1110, YJM813, and YJM263 formed the first clonal cluster.

The genomes APN, Platinum, MLAF.1, ABM2, LSF.1, APK, SXJM4.1, and DBL1 formed the second clonal cluster.

The genomes NRRLYB-3916, BMG, and CBS2919 formed the third clonal cluster.

The genomes APM, API, PYCC4226, UDeb-BY0012sc2, ASL, BS, BTF, UDeb-BY0023, and UDeb-BY0022 formed the fourth clonal cluster.

|  | **AAF similarity** | **SNP similarity** |
| --- | --- | --- |
| **Clonal cluster vs. other genomes** | 0.9987 (first cluster)  0.9980 (second cluster)  0.9977 (third cluster)  0.9976 (fourth cluster) | 0.9729 (first cluster)  0.9723 (second cluster)  0.9698 (third cluster)  0.9680 (fourth cluster) |
| **Inside clonal cluster** | 0.9996 (first cluster)  0.9996 (second cluster)  0.9996 (third cluster)  0.9992 (fourth cluster) | 0.9963 (first cluster)  0.9974 (second cluster)  0.9995 (third cluster)  0.9947 (fourth cluster) |

**Mixed origin ’e’**

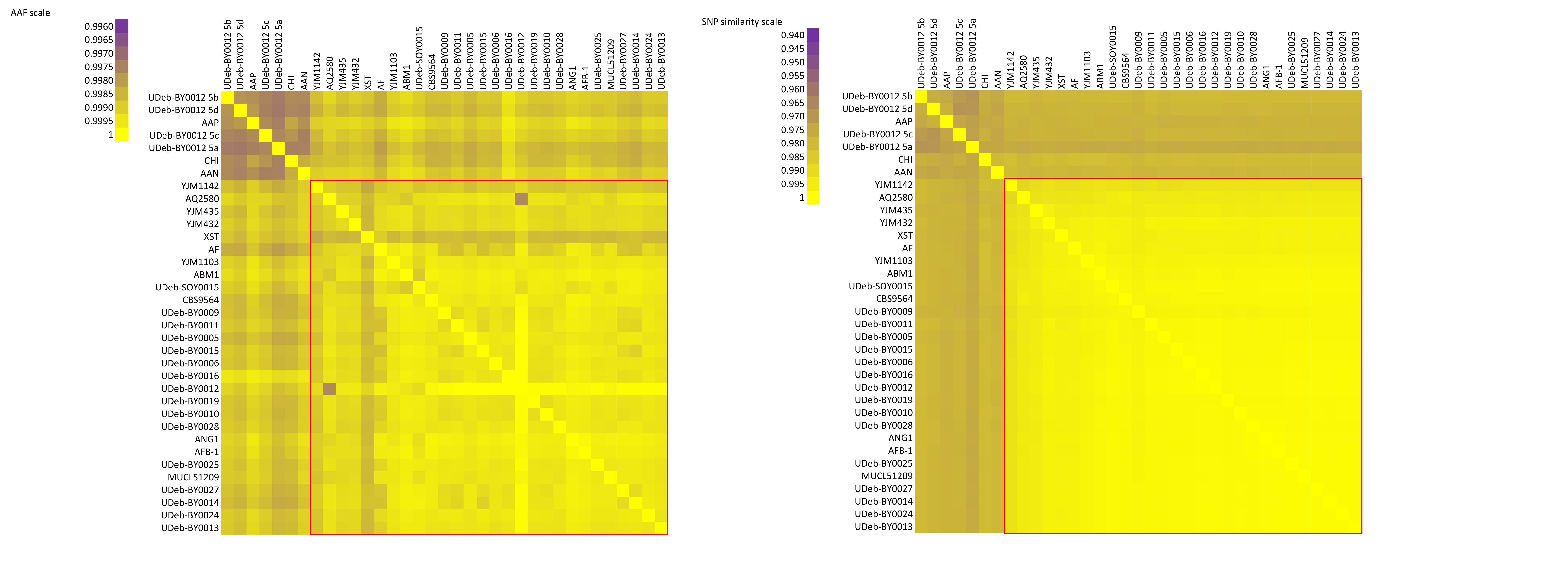

The genomes YJM1142, YJM435, YJM432, CBS9564, UDeb-BY0009, UDeb-BY0011, UDeb-BY0005, UDeb-BY0015, UDeb-BY0006, UDeb-BY0016, UDeb-BY0019, UDeb-BY0010, UDeb-BY0028, AFB-1, UDeb-BY0025, MUCL51209, UDeb-BY0027, UDeb-BY0014, UDeb-BY0024, UDeb-BY0013, AQ2580, XST, AF, YJM1103, ABM1, UDeb-SOY0015, UDeb-BY0012, and ANG1 formed a clonal cluster.

|  | **AAF similarity** | **SNP similarity** |
| --- | --- | --- |
| **Clonal cluster vs. other genomes** | 0.9989 | 0.9787 |
| **Inside clonal cluster** | 0.9994 | 0.9969 |

**Mixed origin ’f’**

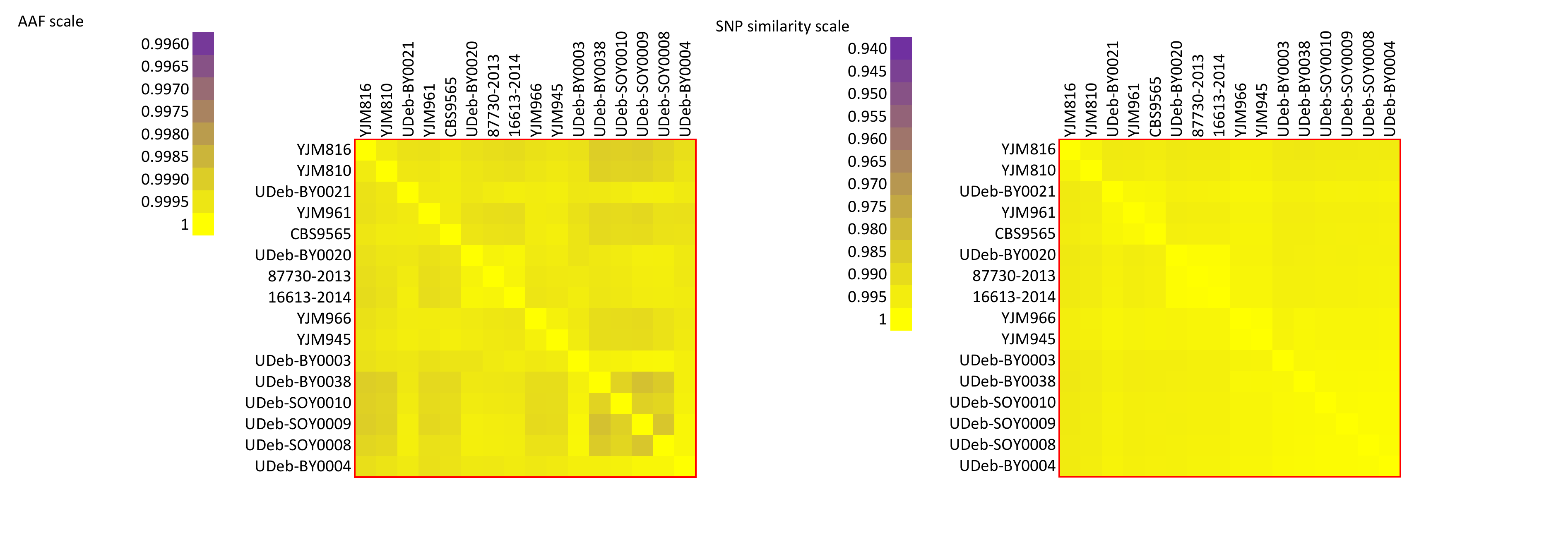

The genomes YJM816, YJM810, UDeb-BY0021, YJM961, CBS9565, UDeb-BY0020, 87730-2013, 16613-2014, YJM966, YJM945, UDeb-BY0003, UDeb-BY0004, UDeb-BY0038, UDeb-SOY0010, UDeb-SOY009, UDeb-SOY008 formed a clonal cluster.

|  | **AAF similarity** | **SNP similarity** |
| --- | --- | --- |
| **Clonal cluster** | 0.9995 | 0.9965 |
