## Supplementary File S3 for "Commercial *Saccharomyces cerevisiae* baker’s yeasts: strain redundancy, genome plasticity, and colonization of the sourdough environment and the human body"

**Supplementary File S3. Coverage and allele ratio plots.**

For every clonal cluster in the Mixed origin subclades, the coverage plots corrected for copy numbers and the allele ratio plots are shown. The isolates and strains belonging to the clonal cluster are listed before plots are shown for the individual genomes.

**aff. Mixed origin**

**aff. Mixed origin clonal cluster of ABM5, ABM4, and ABM3**

ABM5

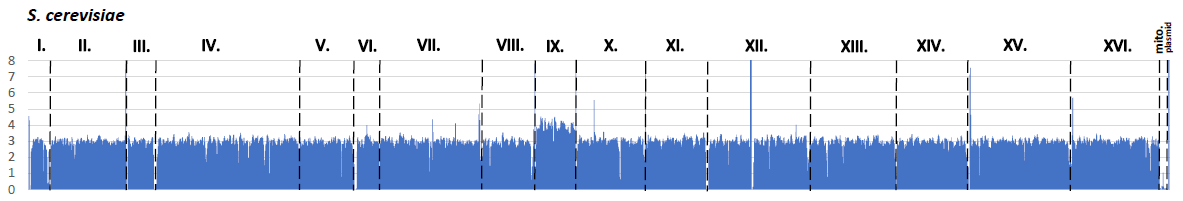

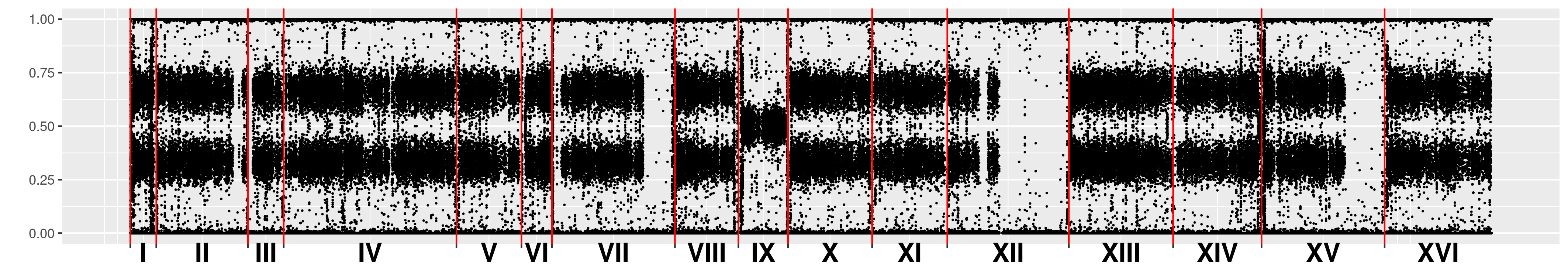

ABM4

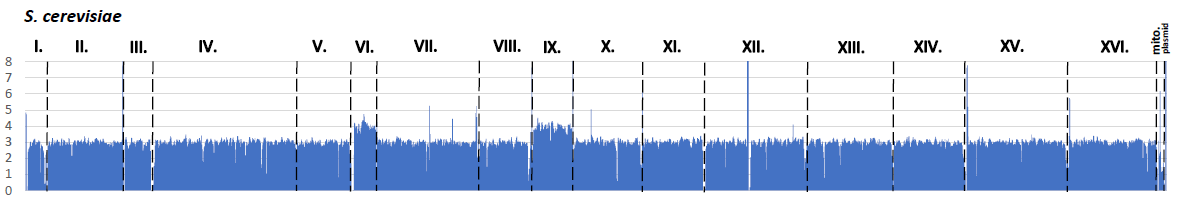

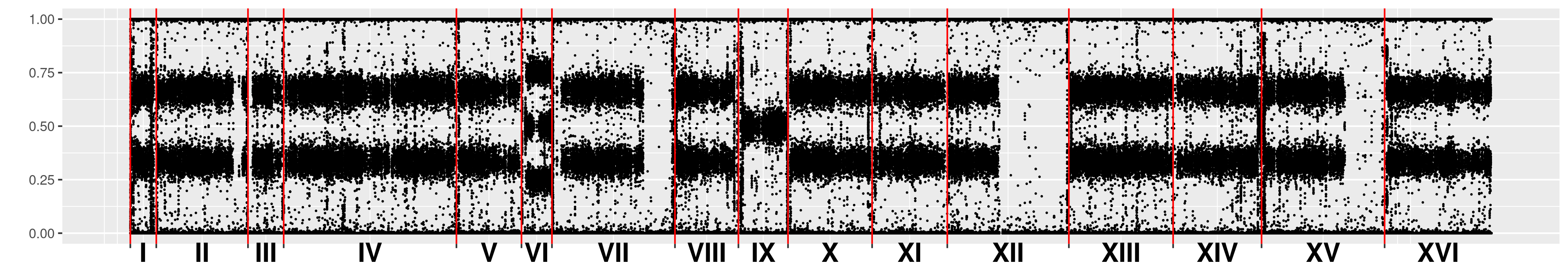

ABM3

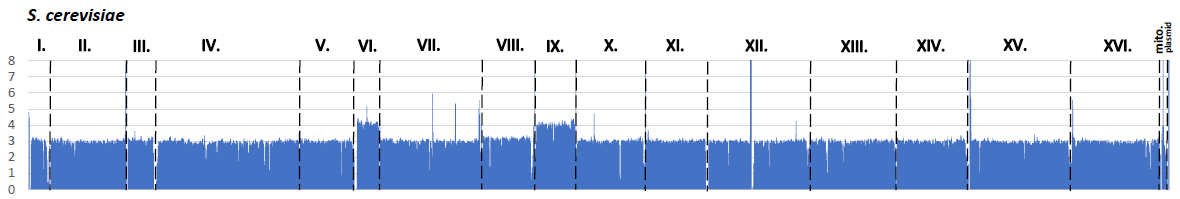

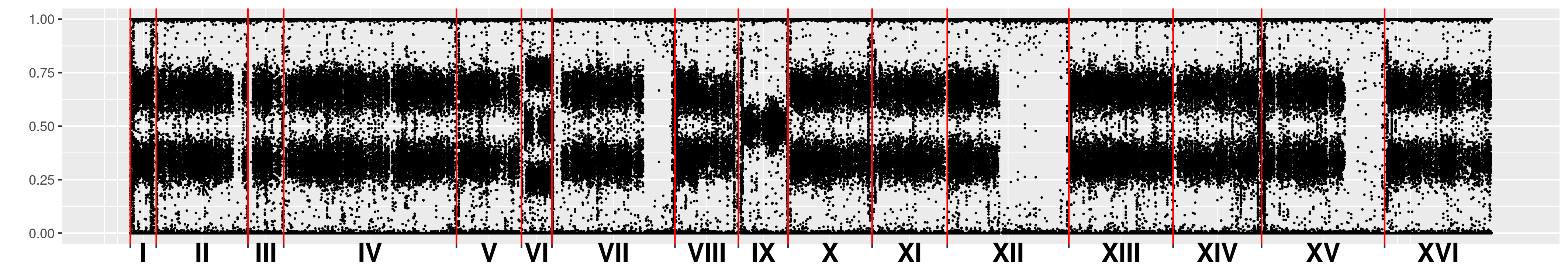

**Mixed origin a**

**Mixed origin a clonal cluster of AP13.1, BY0026, and BR004**

AP13.1

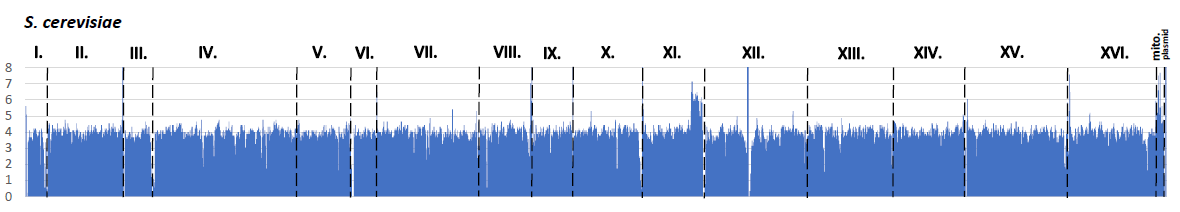

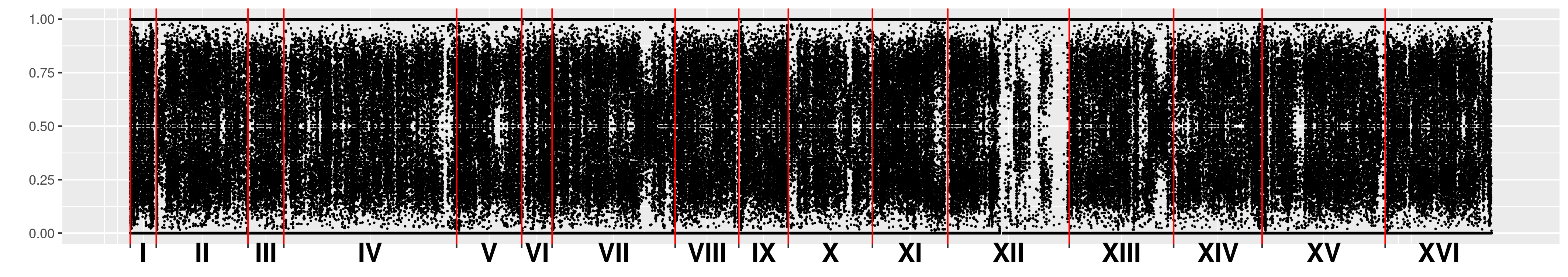

BY0026

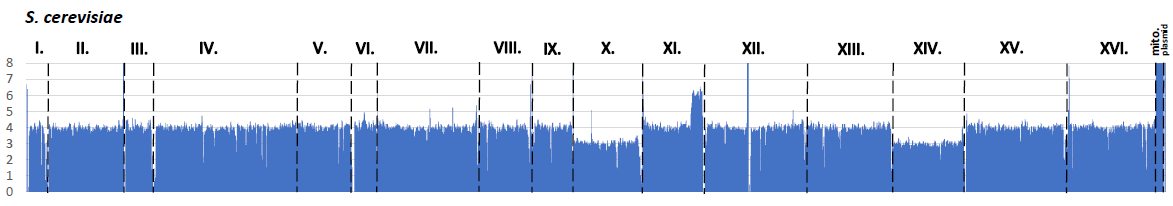

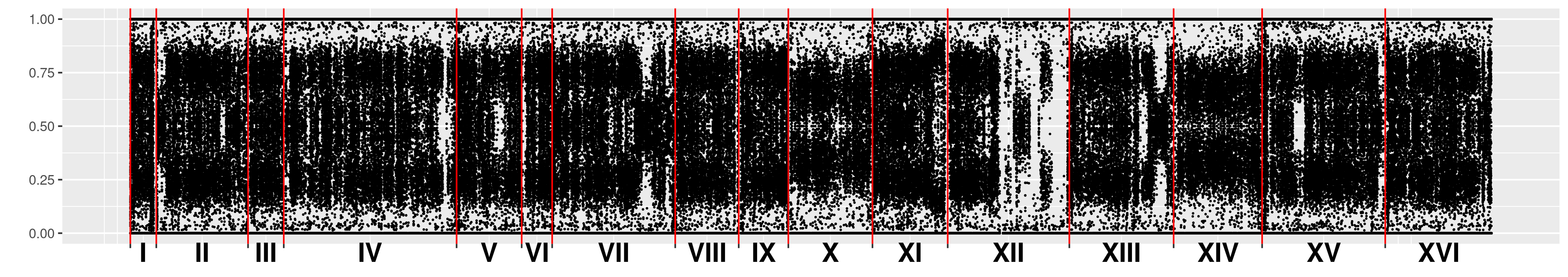

BR004

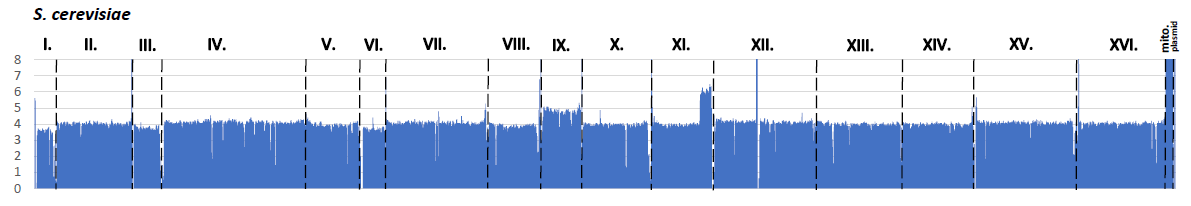

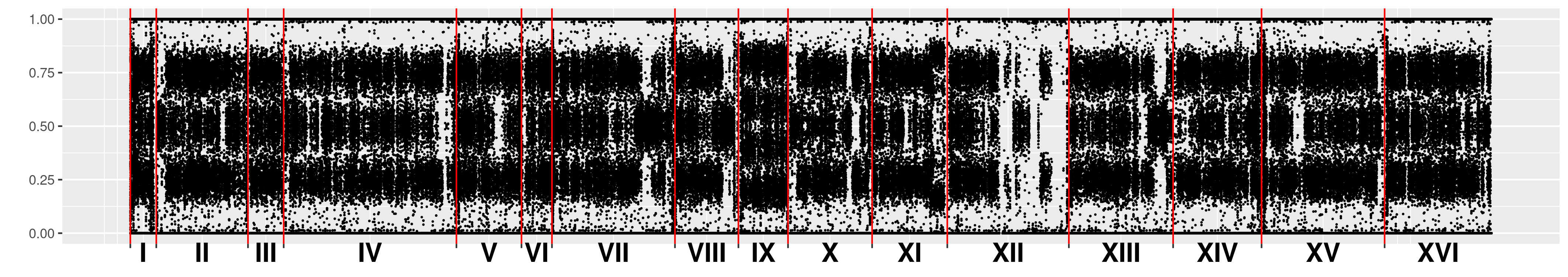

**Mixed origin b**

**Mixed origin b clonal cluster of 6504-2014, UDeb-BY0007, UDeb-BY0001, UDeb-BY0008, UDeb-BY0002, UDeb-BY0018, UDeb-BY0017, UDeb-SOY0012, UDeb-SOY0011, UDeb-SOY0005, UDeb-SOY0006, UDeb-SOY0001, UDeb-SOY0014, UDeb-SOY0013, UDeb-SOY0007, and BTD**

6504-2014

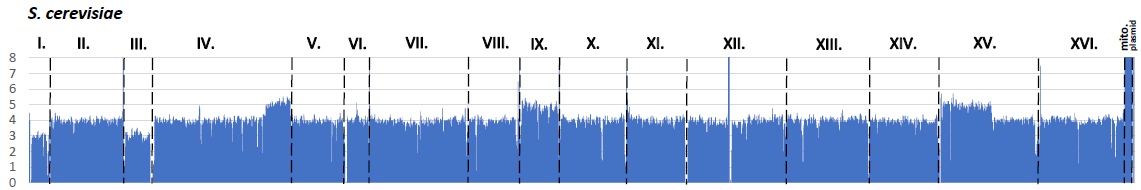

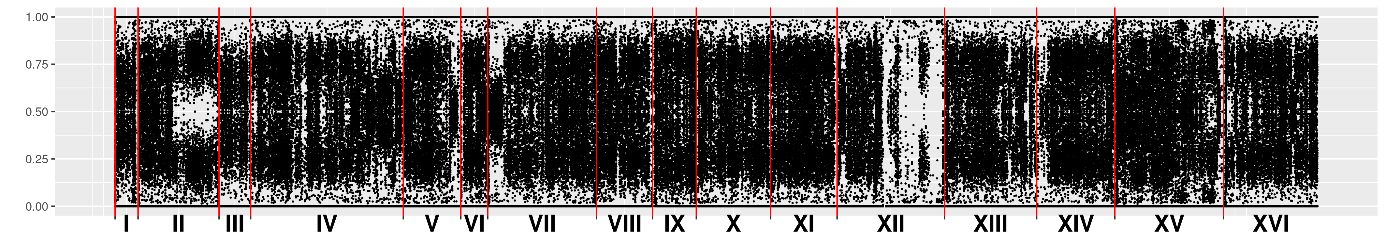

UDeb-BY0007

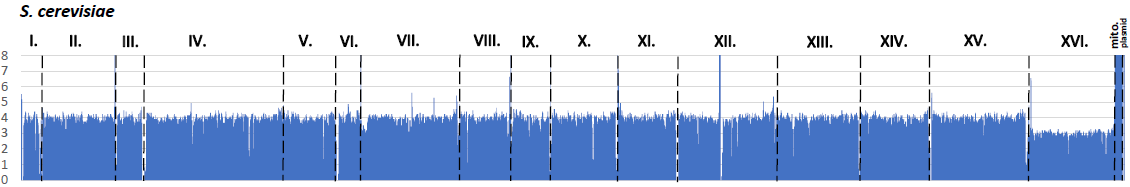

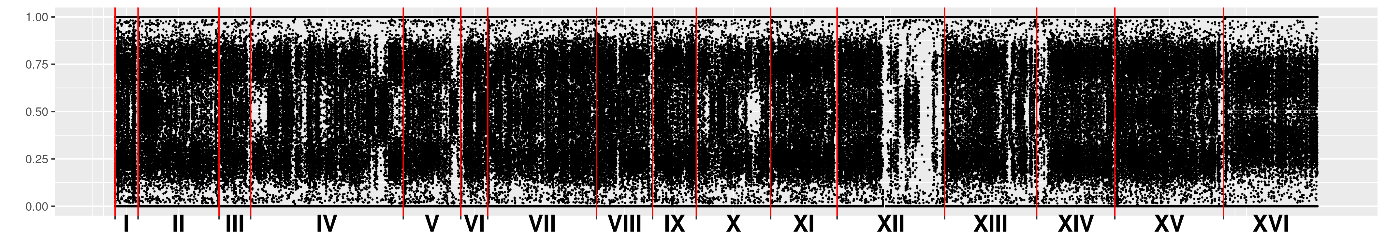

UDeb-BY0001

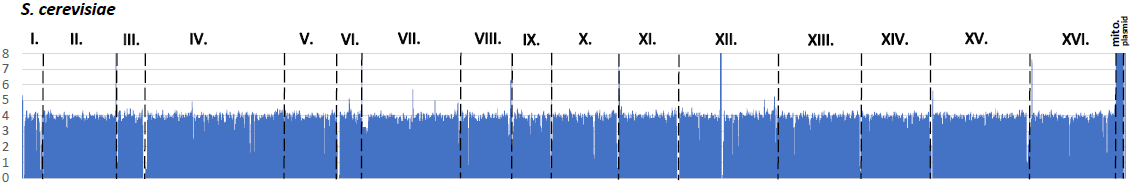

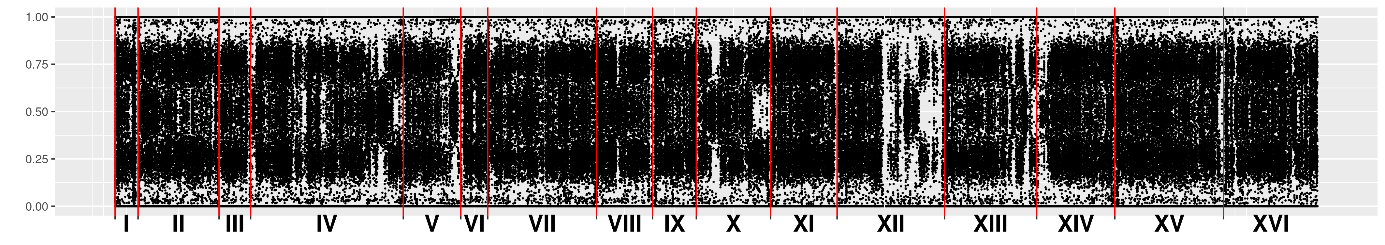

UDeb-BY0008

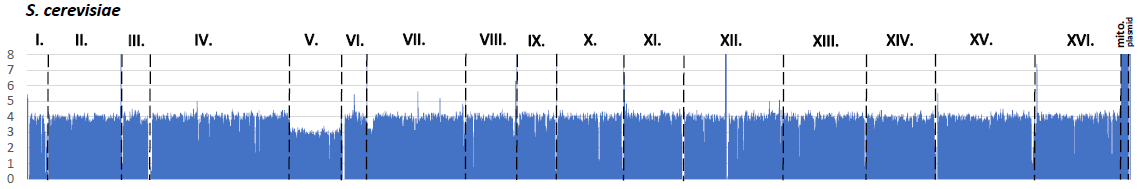

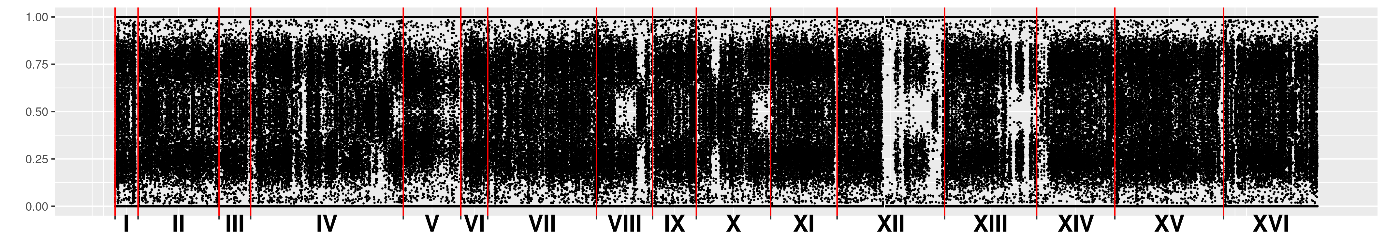

UDeb-BY0002

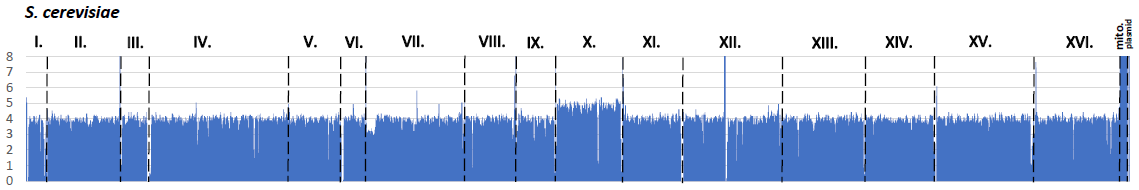

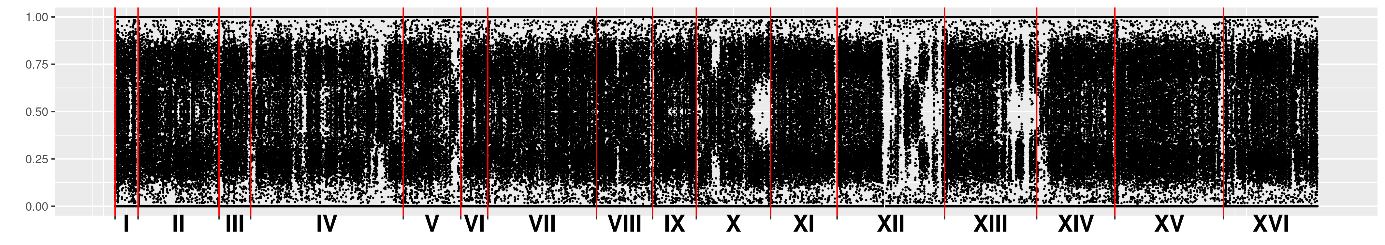

UDeb-BY0018

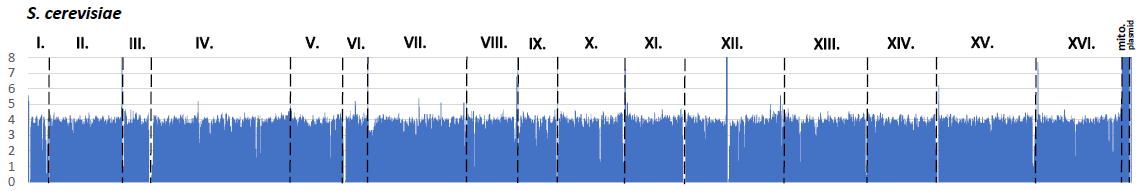

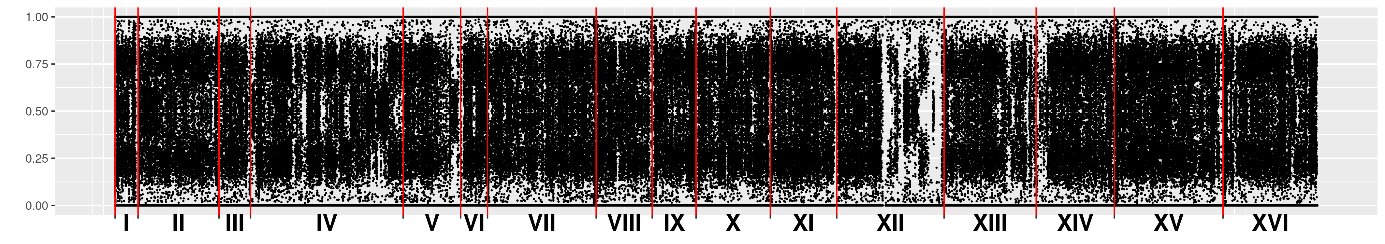

UDeb-BY0017

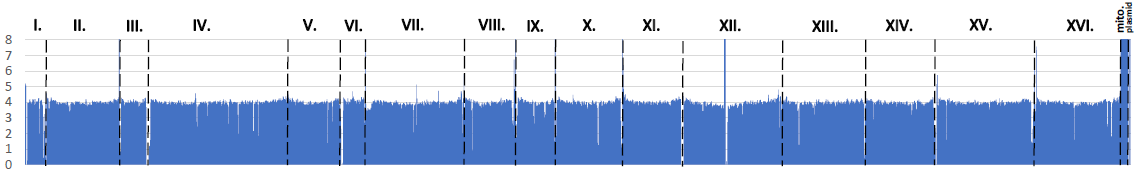

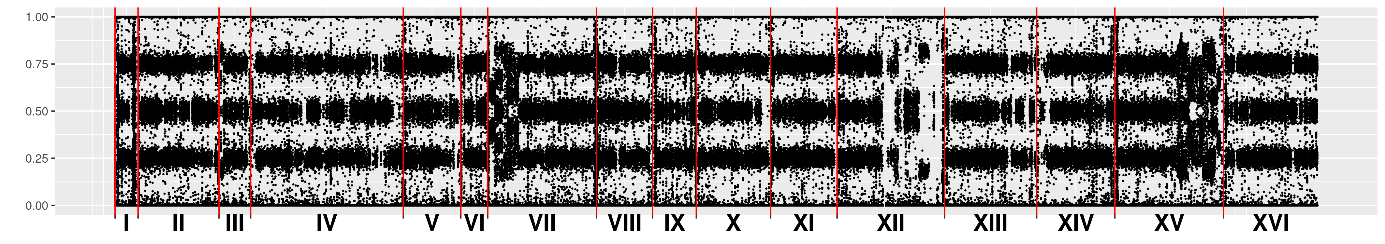

UDeb-SOY0012

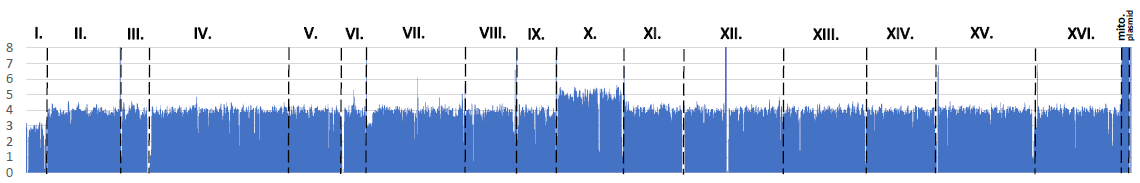

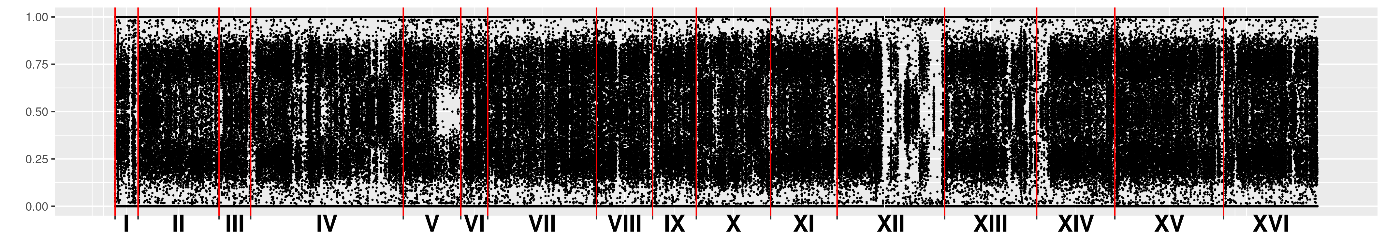

UDeb-SOY0011

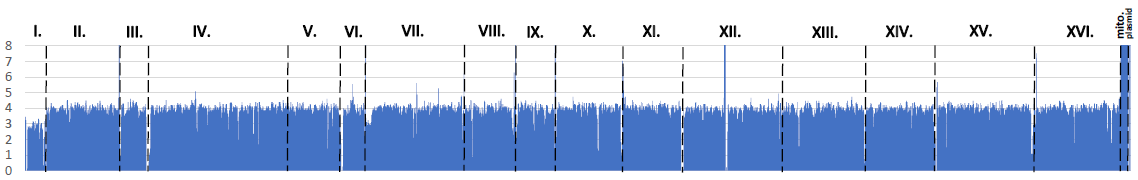

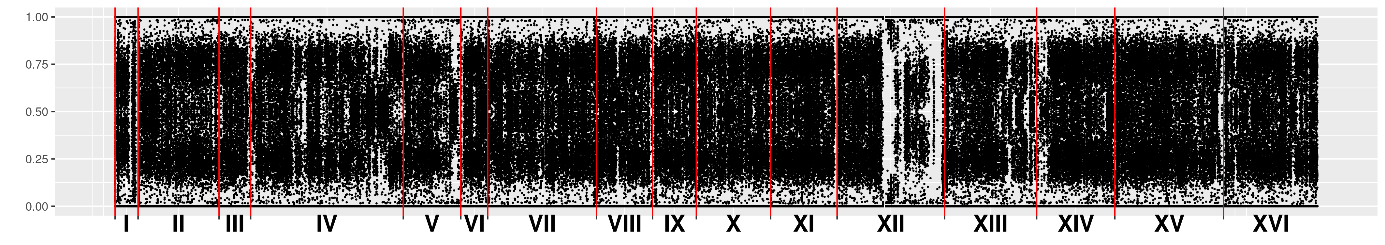

UDeb-SOY0005

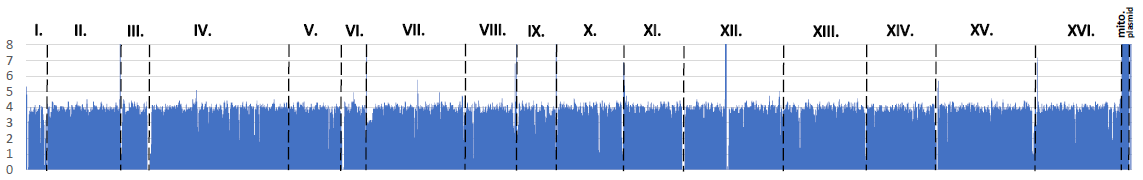

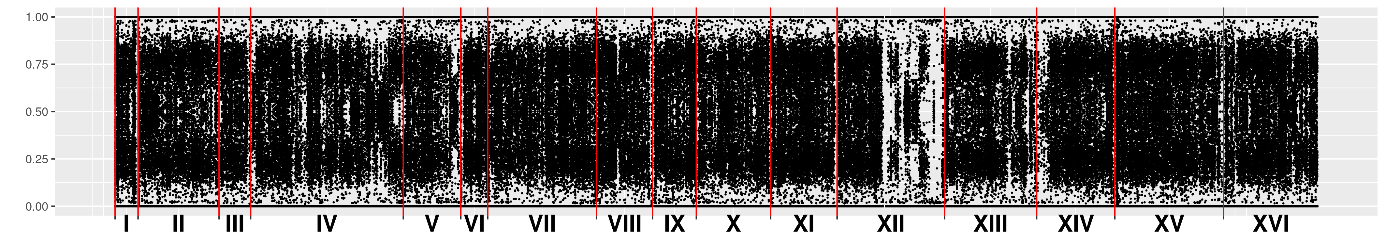

UDeb-SOY0006

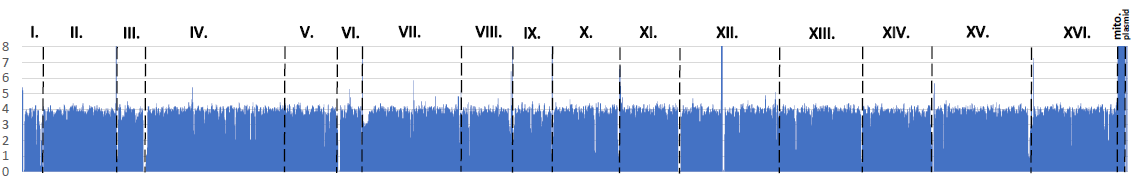

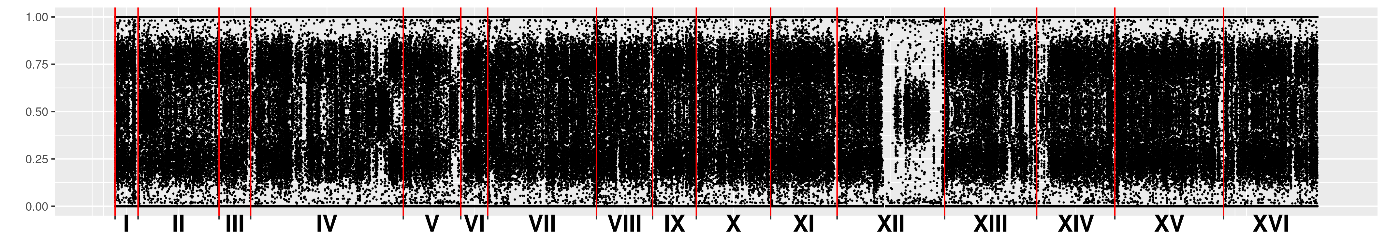

UDeb-SOY0001

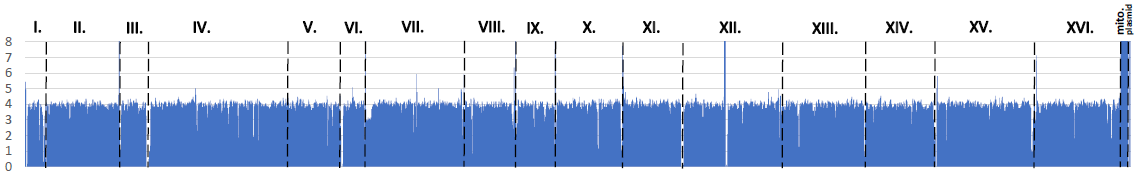

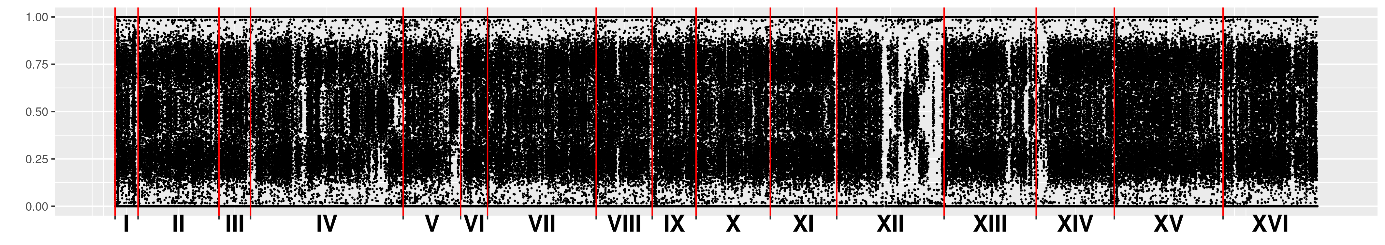

UDeb-SOY0014

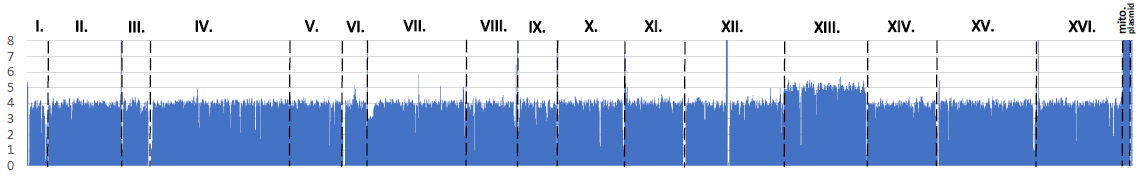

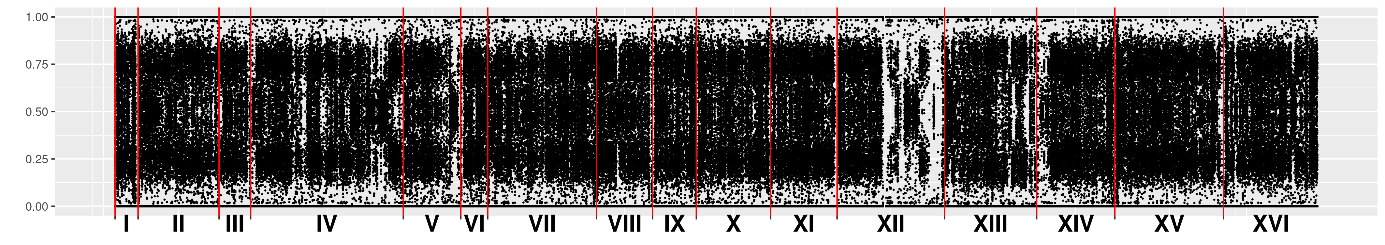

UDeb-SOY0013

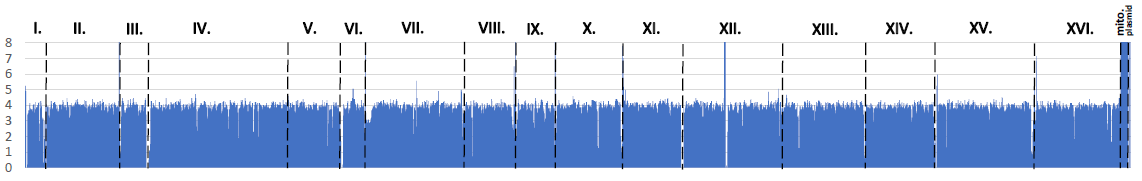

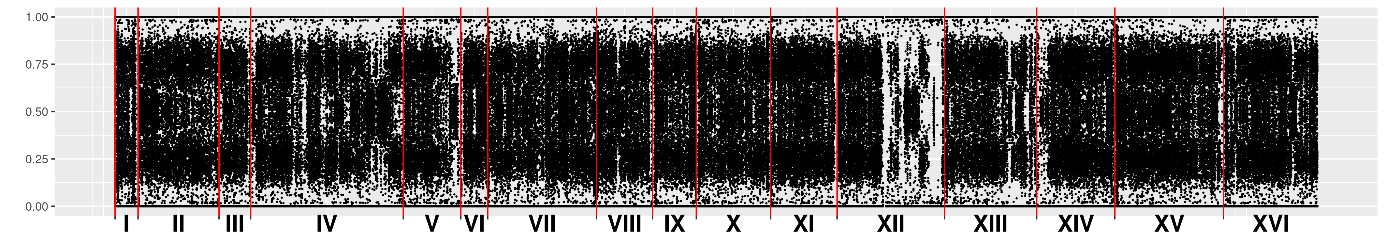

UDeb-SOY0007

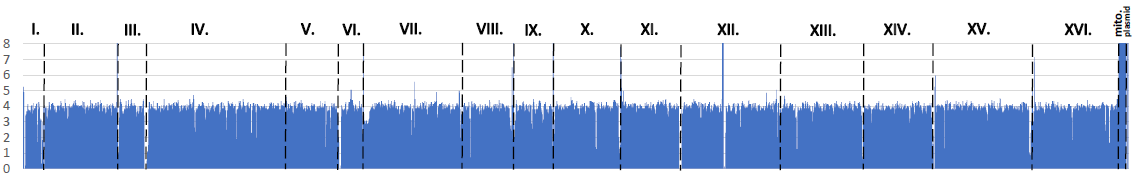

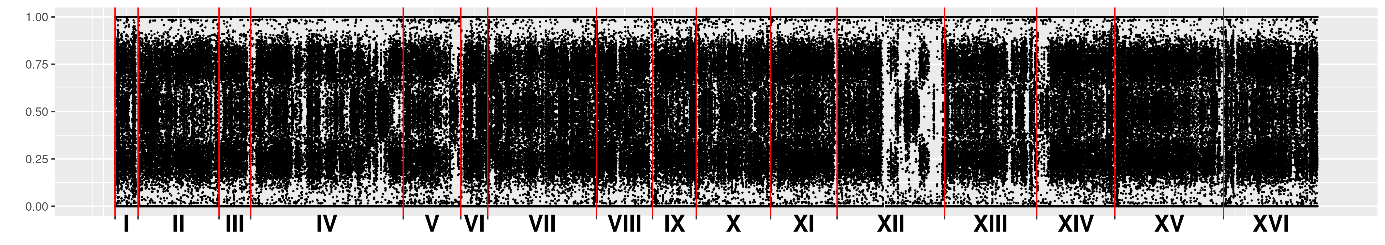

BTD

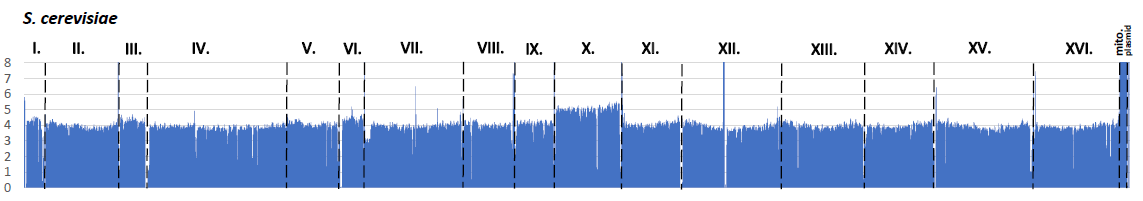

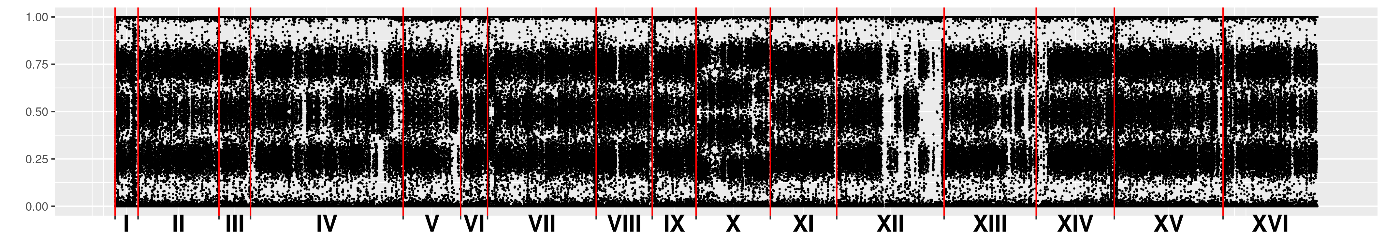

**Mixed origin c**

**Mixed origin c clonal cluster of G17, BMM, CKK, CBF, BTE, CLA, AQ2593, CKS, BR001, UDeb-SOY0004, UDeb-SOY0003, UDeb-SOY0002, YJM958, CBS9562, YJM959, YJM951, YJM962, YJM960, YJM811, YJM1109, YJM936, YJM1118, YJM1113, YJM953, YJM561, and BML**

G17

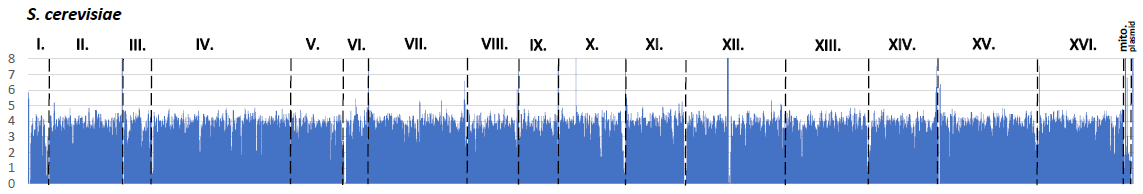

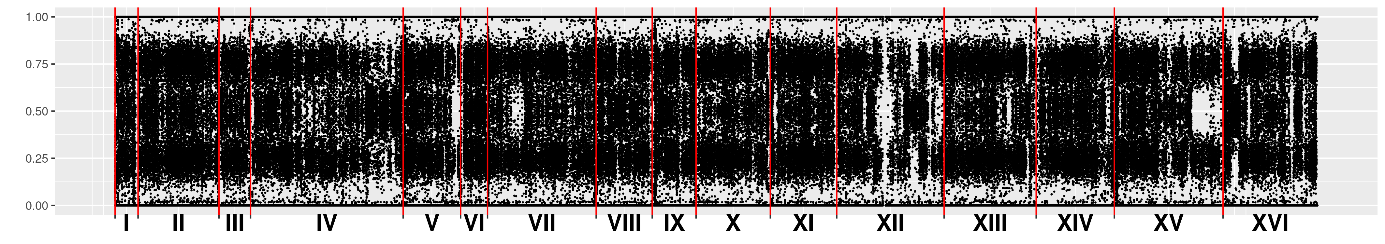

BMM

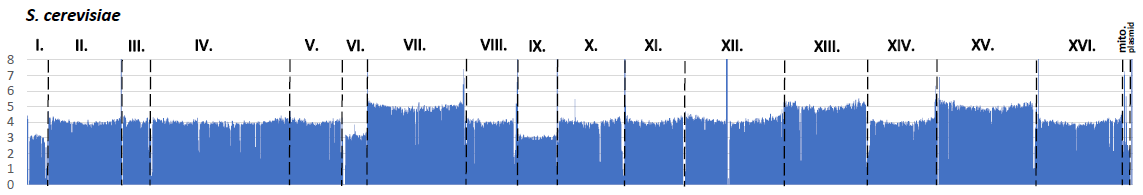

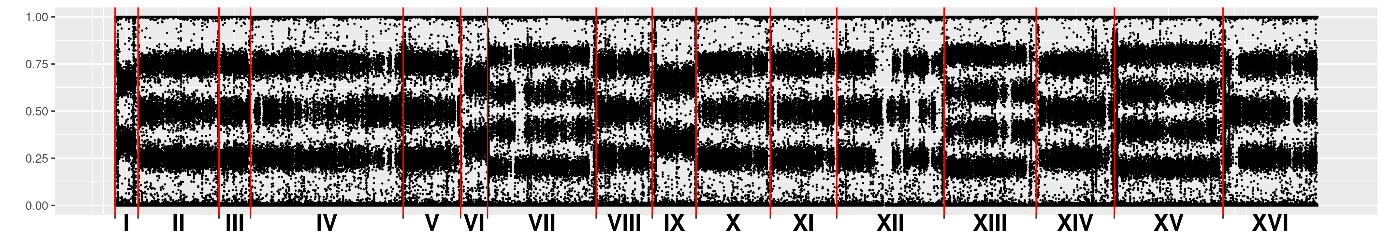

CKK

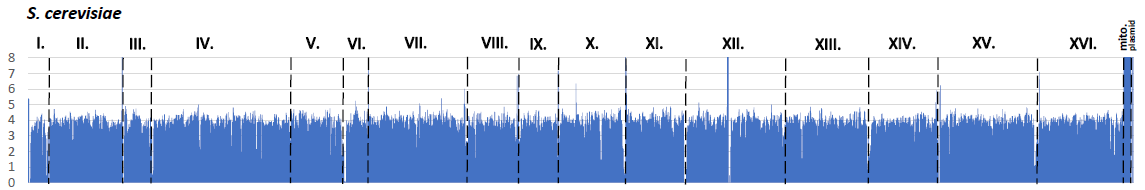

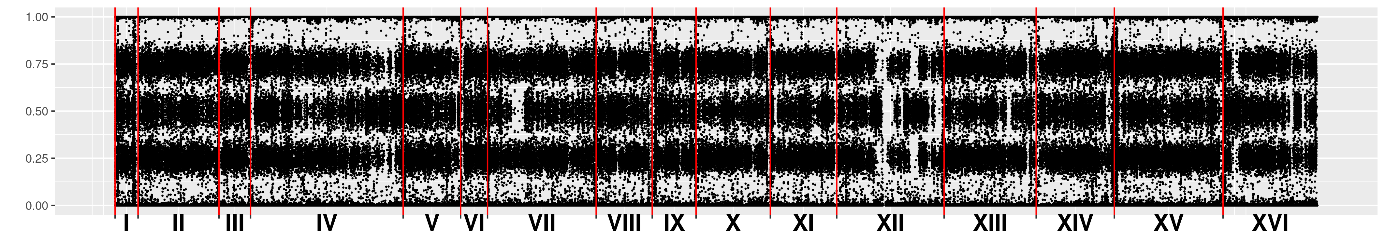

CBF

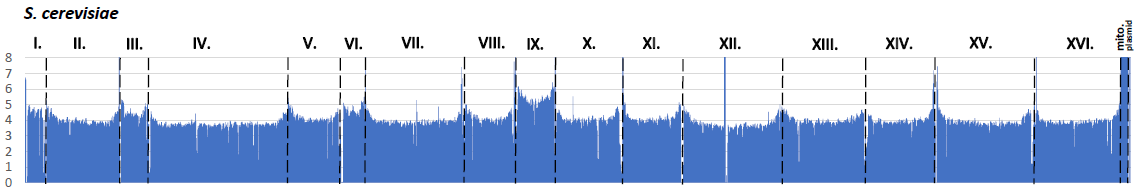

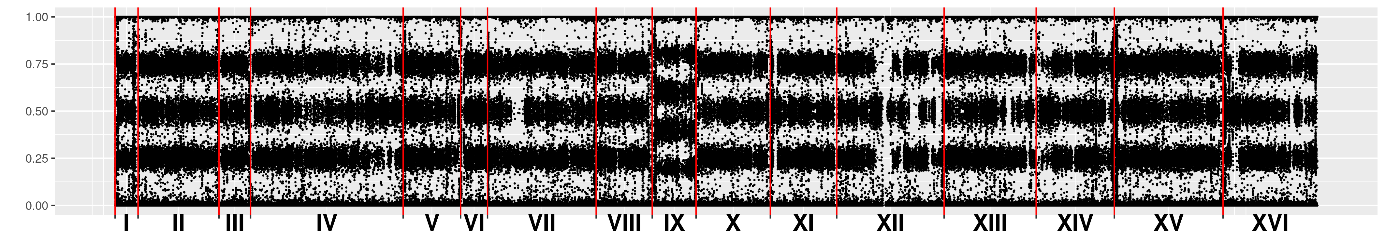

BTE

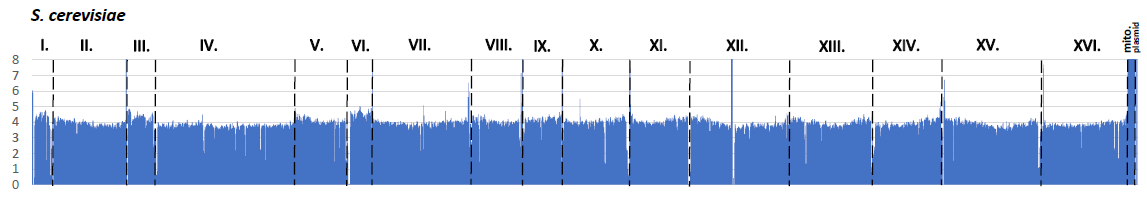

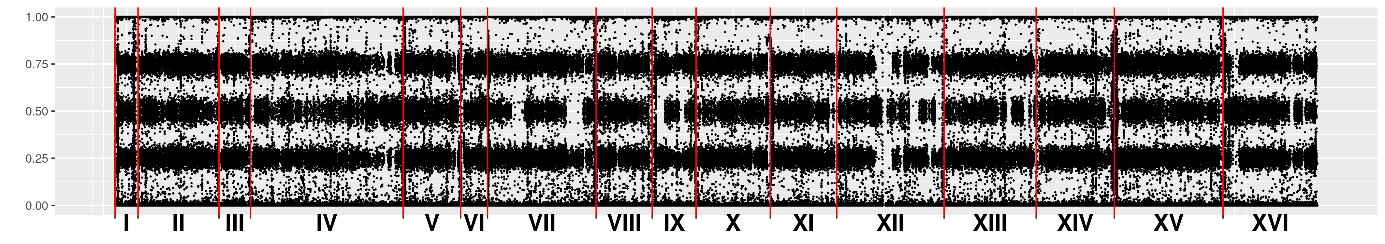

CLA

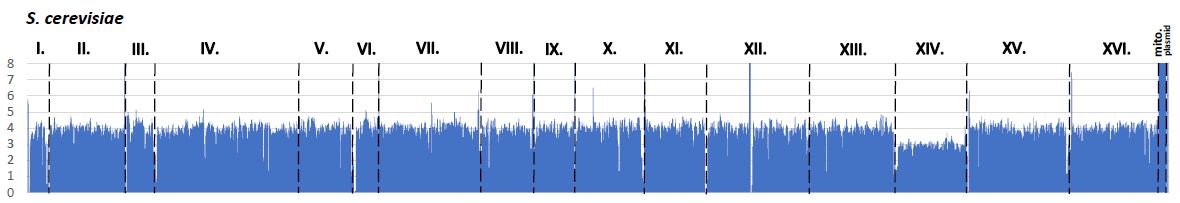

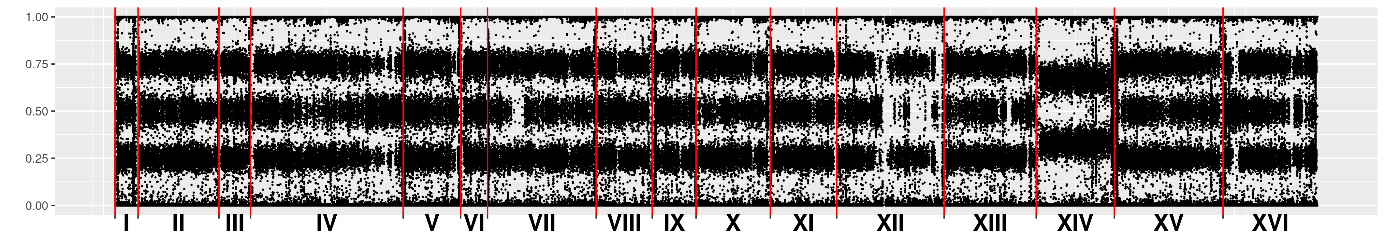

AQ2593

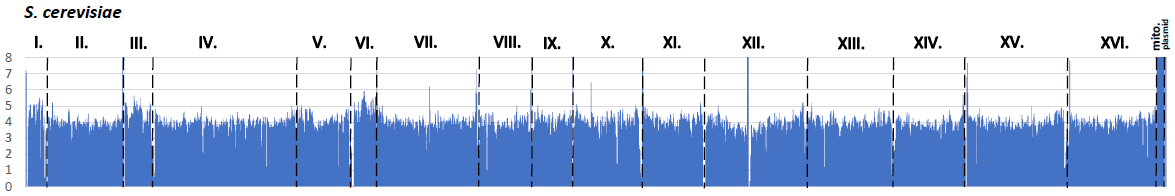

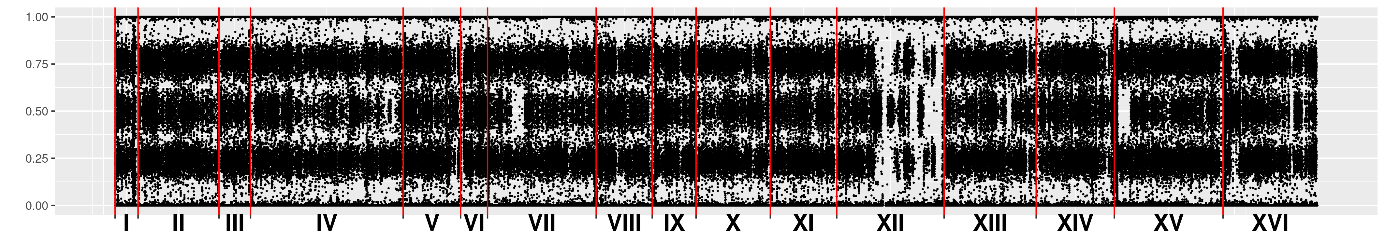

CKS

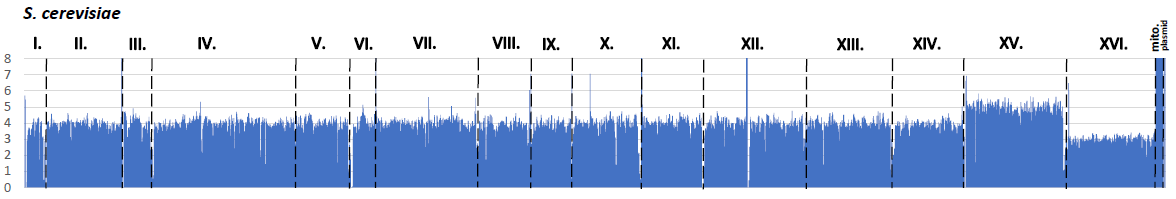

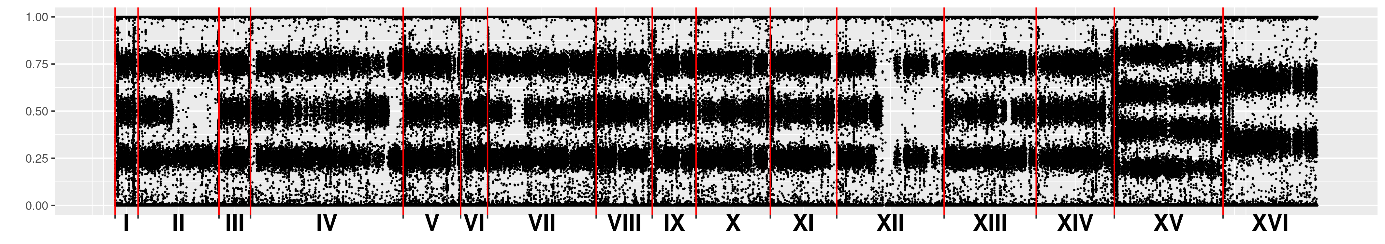

BR001

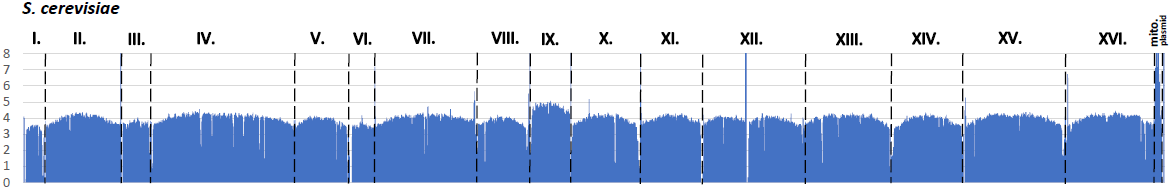

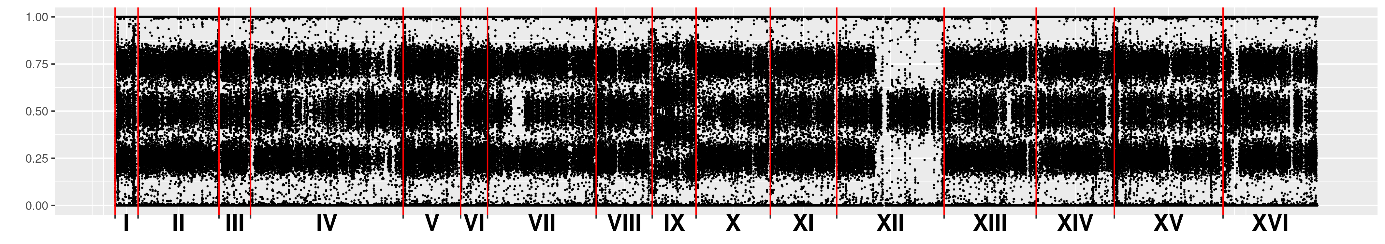

UDeb-SOY0004

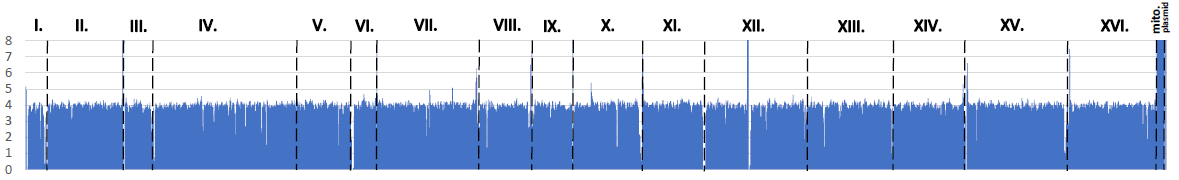

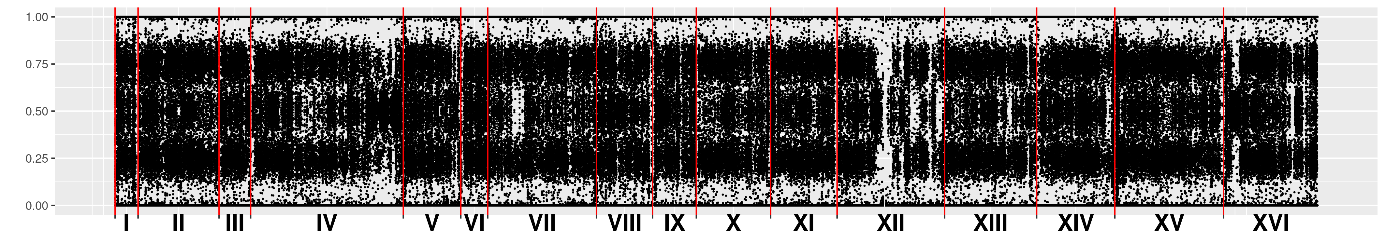

UDeb-SOY0003

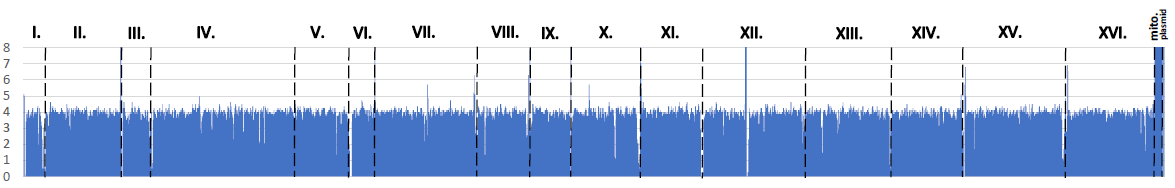

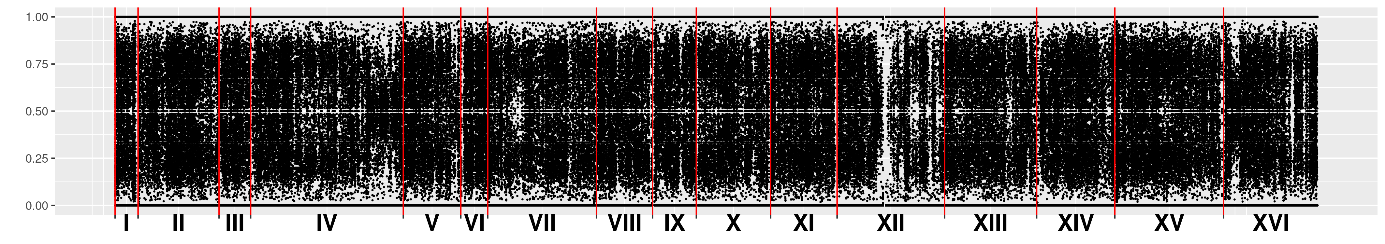

UDeb-SOY0002

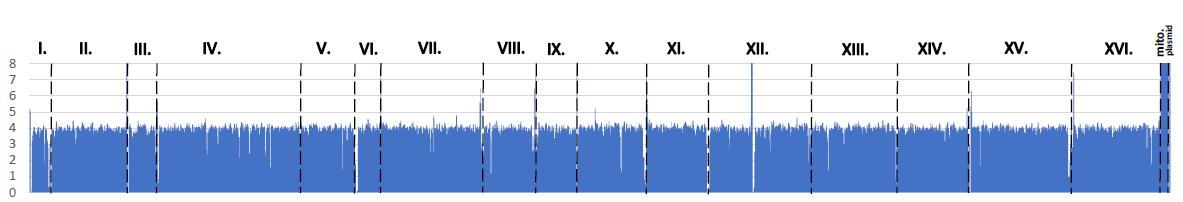

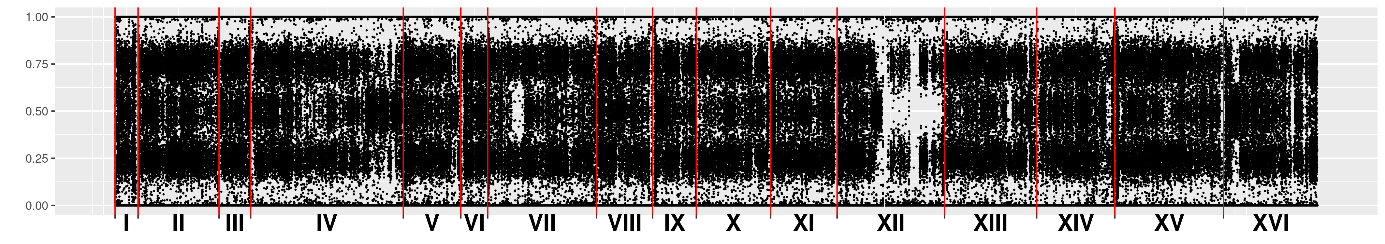

YJM958

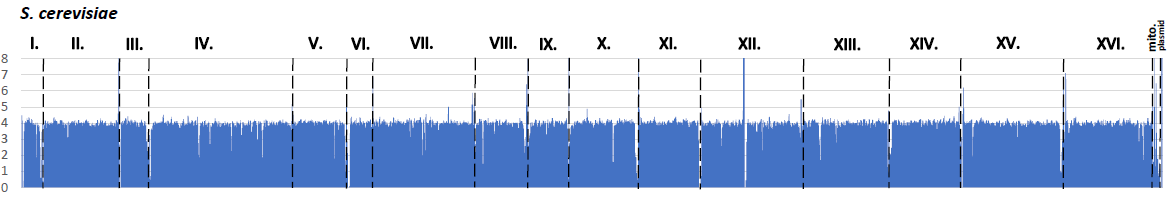

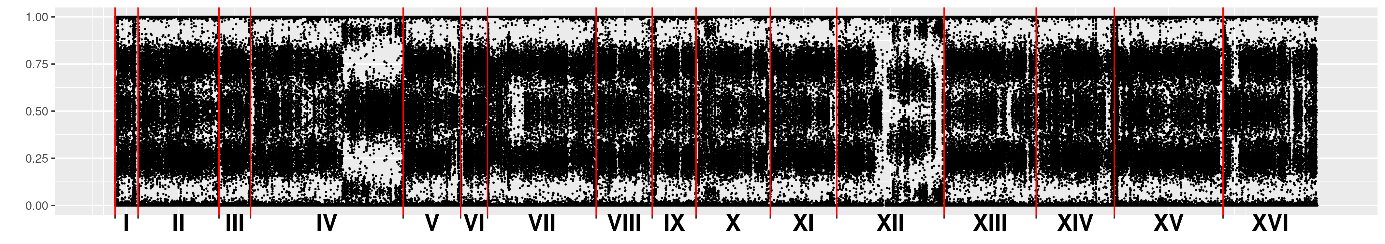

CBS9562

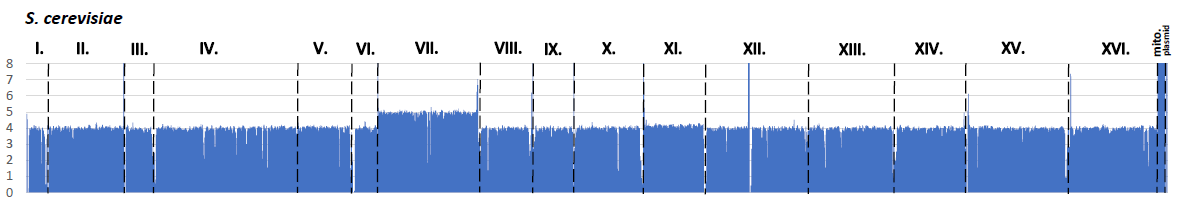

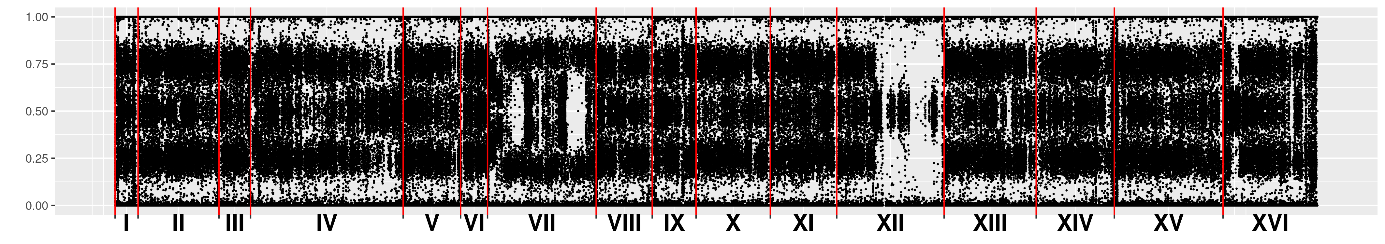

YJM959

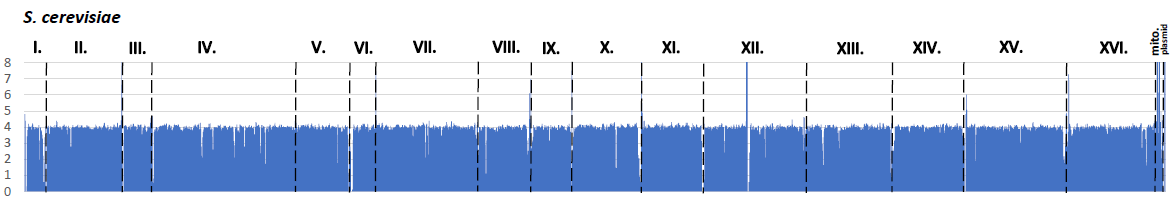

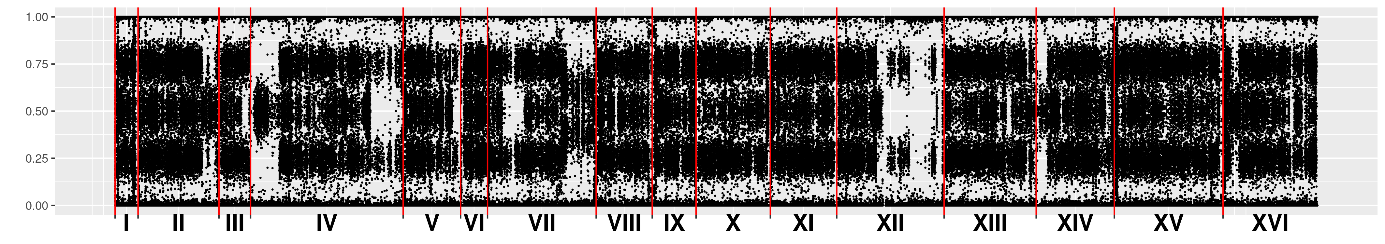

YJM951

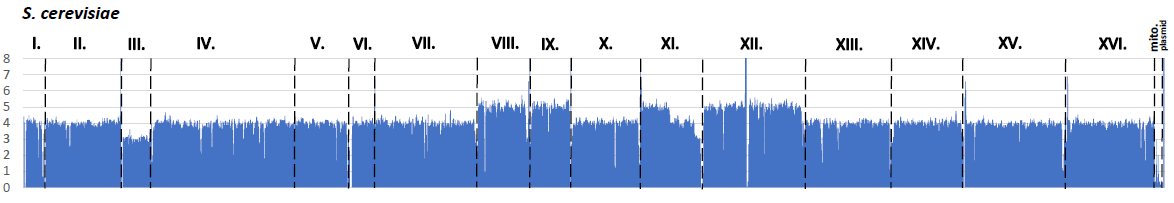

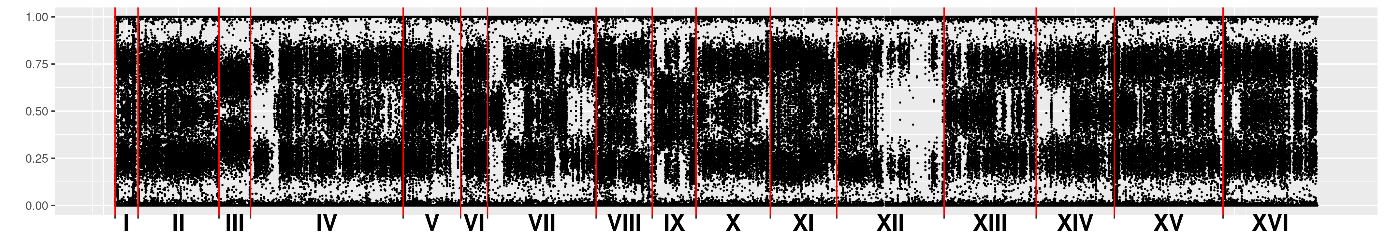

YJM962

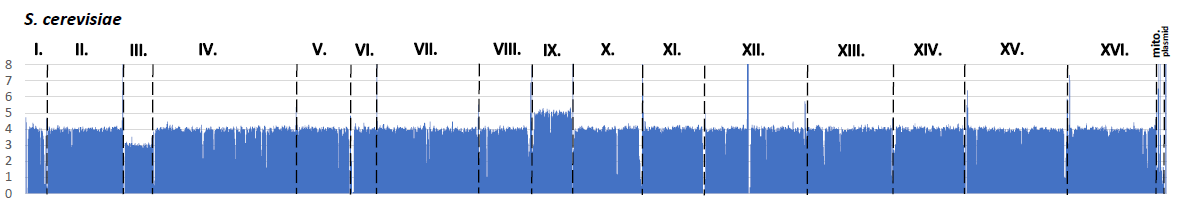

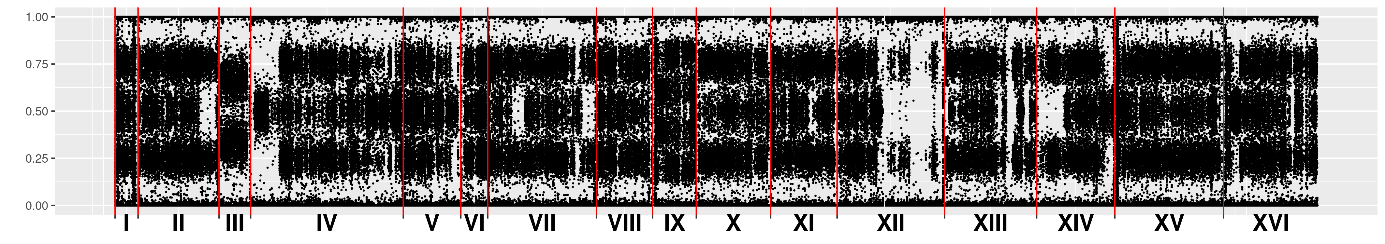

YJM960

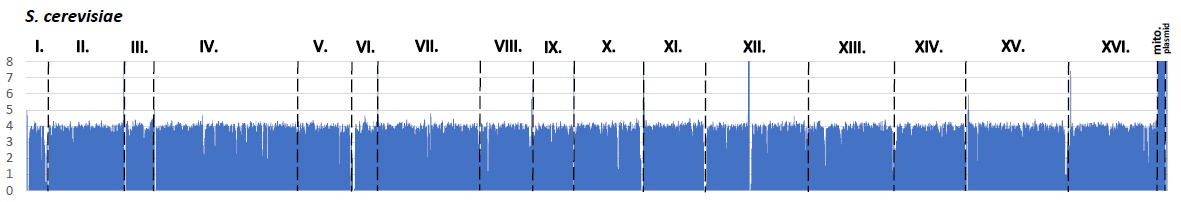

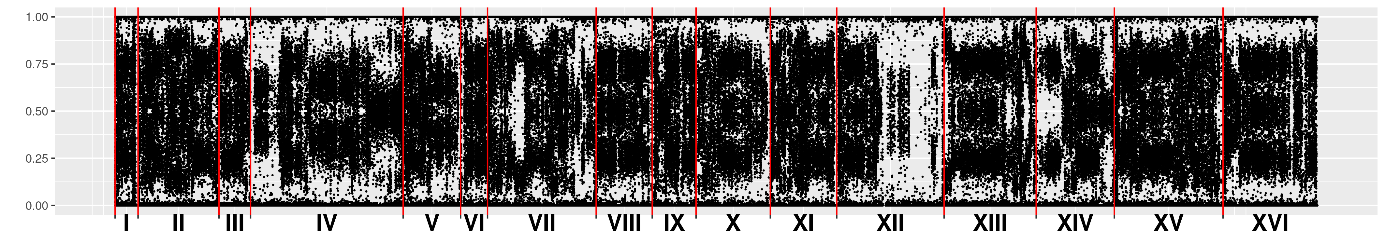

YJM811

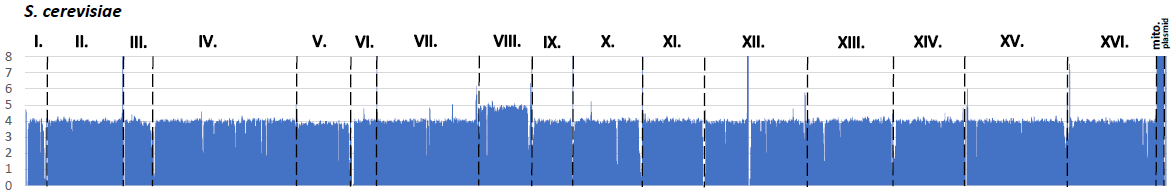

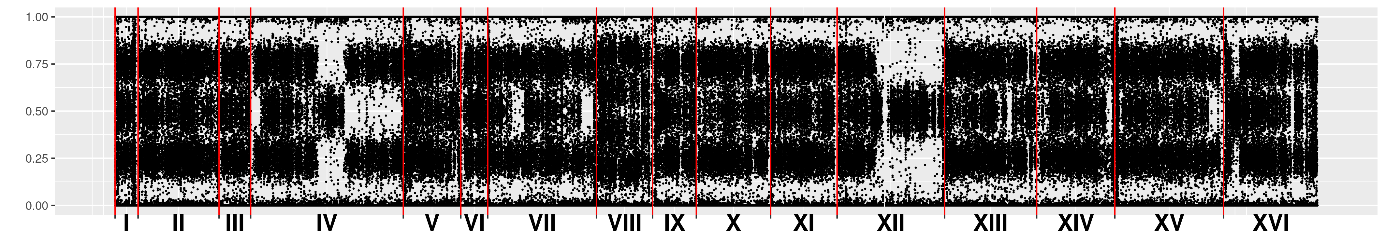

YJM1109

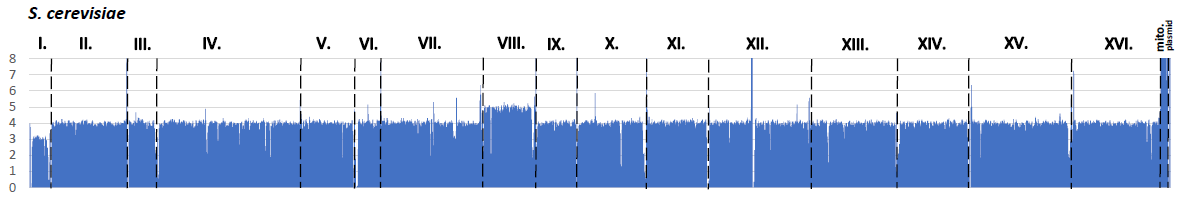

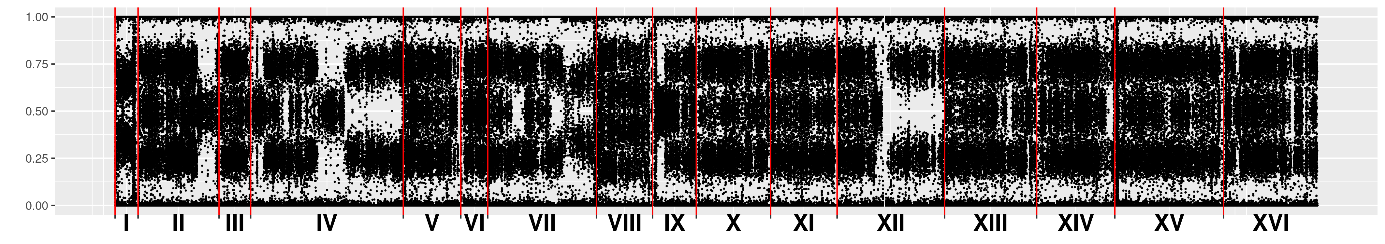

YJM936

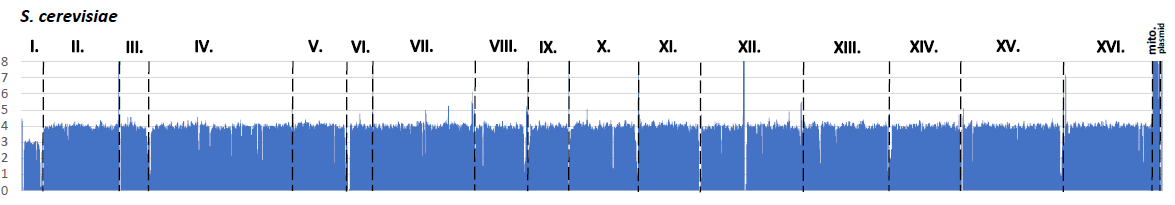

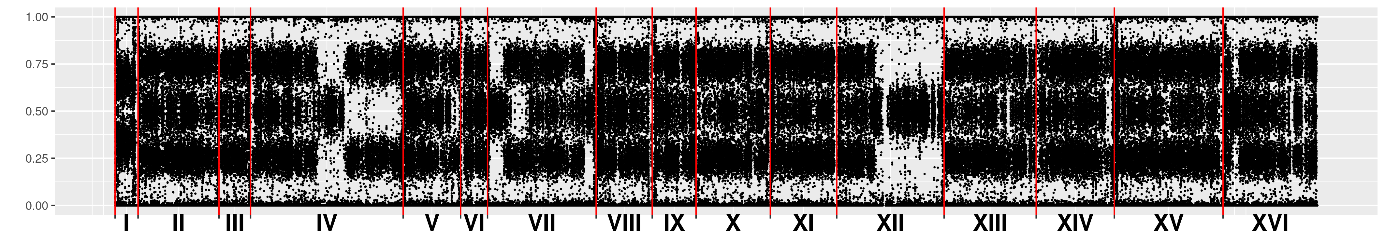

YJM1118

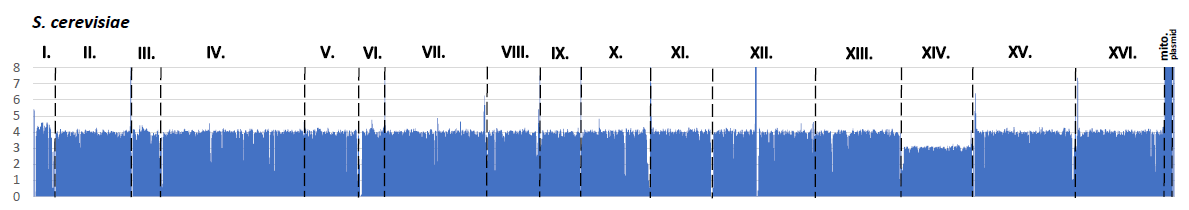

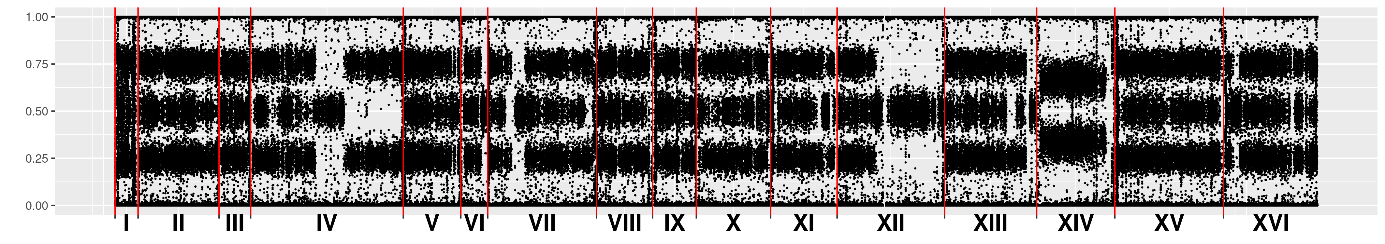

YJM1113

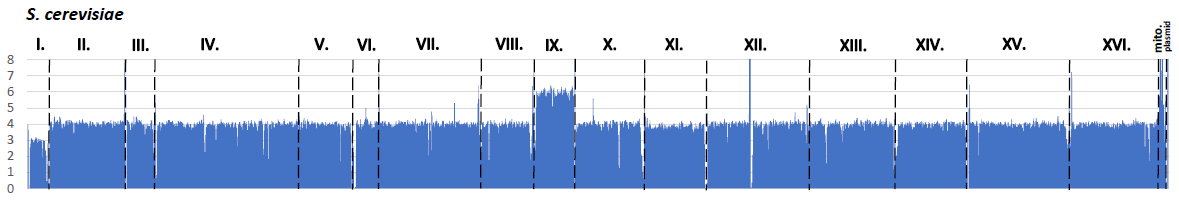

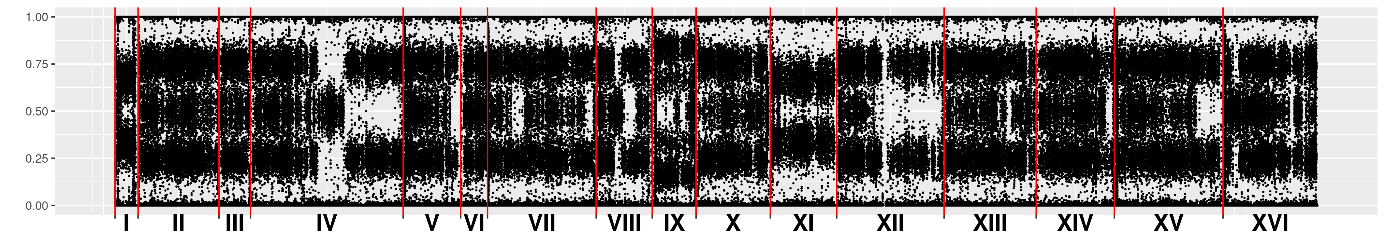

YJM953

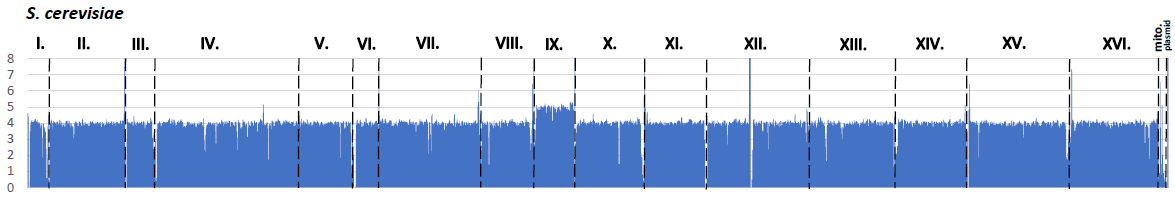

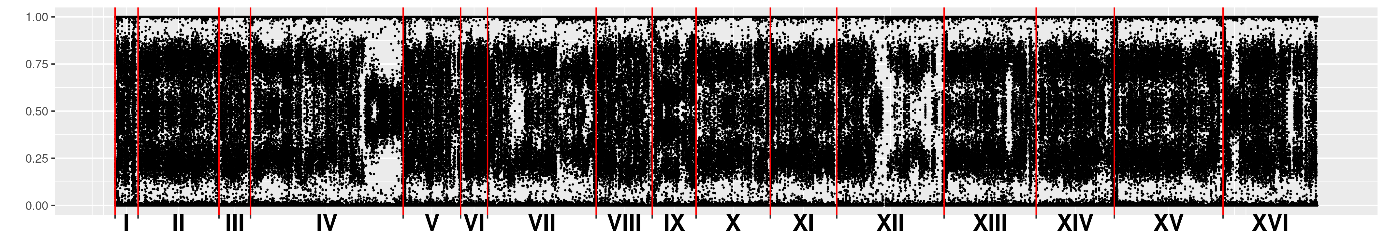

YJM561

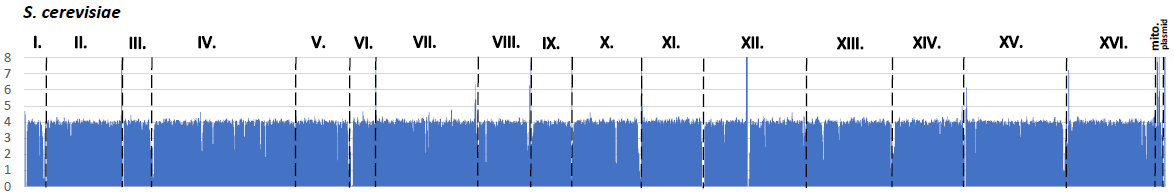

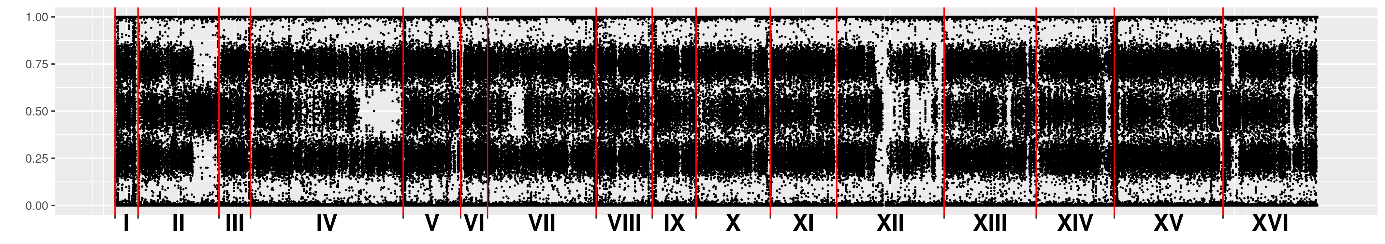

BML

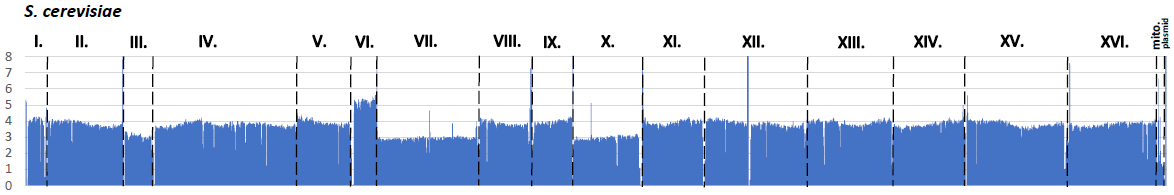

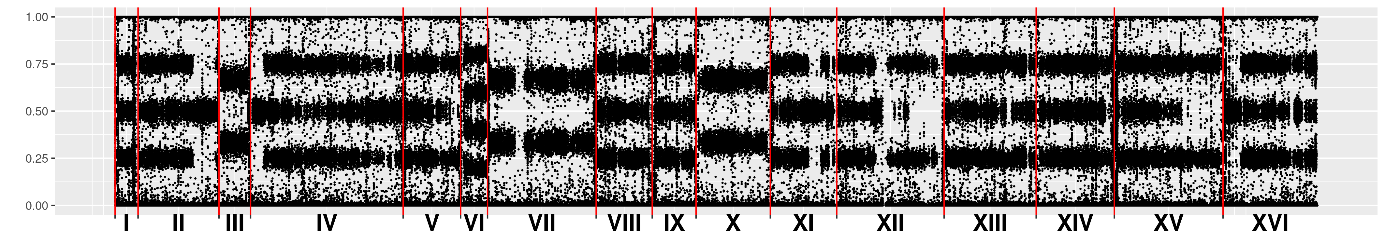

**Mixed origin d**

**Mixed origin d clonal cluster of YJM1120, YJM1110, YJM813, and YJM263**

YJM1120

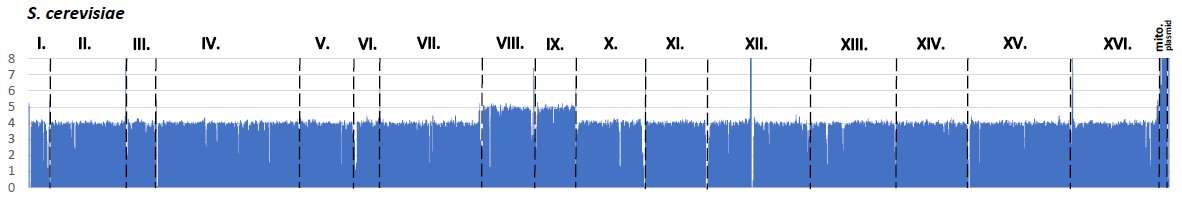

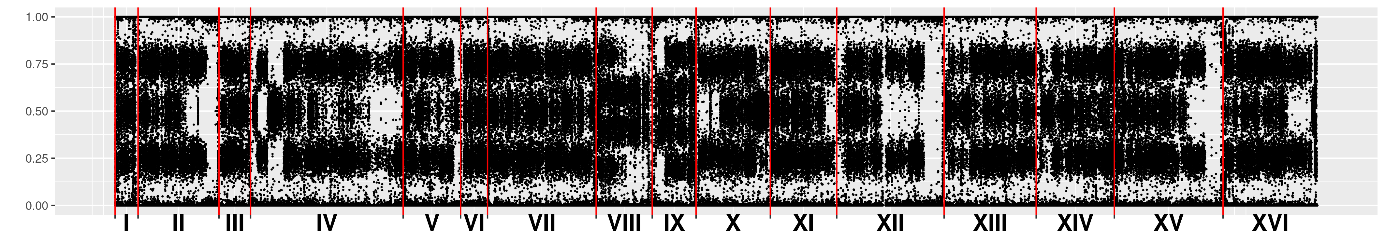

YJM1110

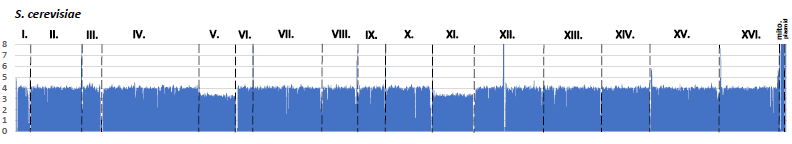

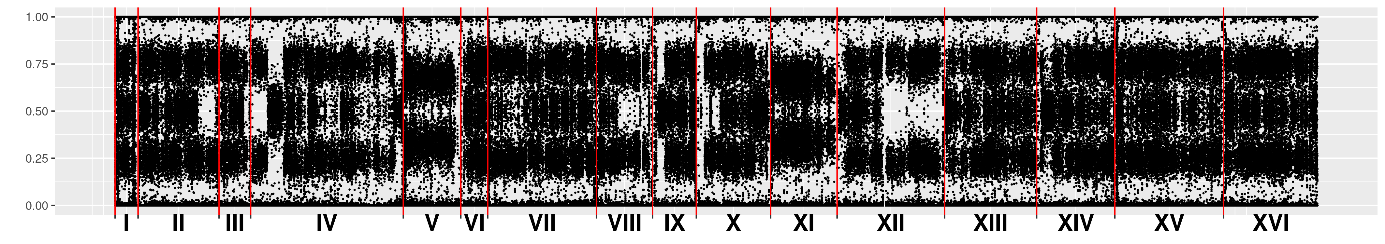

YJM813

YJM263

**Mixed origin d clonal cluster of APN, Platinum, MLAF.1, ABM2, LSF.1, APK, SXJM4.1, and DBL1**

APN

Platinum

MLAF.1

ABM2

LSF.1

APK

SXJM4.1

DBL1

**Mixed origin d clonal cluster of NRRLYB-3916, BMG, and CBS2919**

NRRLYB-3916

BMG

CBS2919

**Mixed origin d clonal cluster of APM, API, PYCC4226, UDeb-BY0012sc2, ASL, BS, BTF, UDeb-BY0023, and UDeb-BY0022**

APM

API

PYCC4226

UDeb-BY0012sc2

ASL

BS

BTF

UDeb-BY0023

UDeb-BY0022

**Mixed origin e**

**Mixed origin e clonal cluster of YJM1142, YJM435, YJM432, CBS9564, UDeb-BY0009, UDeb-BY0011, UDeb-BY0005, UDeb-BY0015, UDeb-BY0006, UDeb-BY0016, UDeb-BY0019, UDeb-BY0010, UDeb-BY0028, AFB-1, UDeb-BY0025, MUCL51209, UDeb-BY0027, UDeb-BY0014, UDeb-BY0024, UDeb-BY0013, AQ2580, XST, AF, YJM1103, ABM1, UDeb-SOY0015, UDeb-BY0012, and ANG1**

YJM1142

YJM435

YJM432

CBS9564

UDeb-BY0009

UDeb-BY0011

UDeb-BY0005

UDeb-BY0015

UDeb-BY0006

UDeb-BY0016

UDeb-BY0019

UDeb-BY0010

UDeb-BY0028

AFB-1

UDeb-BY0025

MUCL51209

UDeb-BY0027

UDeb-BY0014

UDeb-BY0024

UDeb-BY0013

AQ2580

XST

AF

YJM1103

ABM1

UDeb-SOY0015

UDeb-BY0012

ANG1

**Mixed origin f**

**Mixed origin f clonal cluster of YJM816, YJM810, UDeb-BY0021, YJM961, CBS9565, UDeb-BY0020, 87730-2013, 16613-2014, YJM966, YJM945, UDeb-BY0003, UDeb-BY0004, UDeb-BY0038, UDeb-SOY0010, UDeb-SOY009, UDeb-SOY008**

YJM816

YJM810,

UDeb-BY0021

YJM961

CBS9565

UDeb-BY0020

87730-2013

16613-2014

YJM966

YJM945

UDeb-BY0003

UDeb-BY0004

UDeb-BY0038

UDeb-SOY0010

UDeb-SOY0009

UDeb-SOY0008
