## Supplementary File S4 for "Commercial *Saccharomyces cerevisiae* baker’s yeasts: strain redundancy, genome plasticity, and colonization of the sourdough environment and the human body"

**Supplementary File S4. Gross chromosomal rearrangements.**

The various types of large deletions and duplications are listed according to chromosomes. Affected clonal clusters are listed. Abbreviations were given to all detected GCR events, these are the same as in Figure 6–8 and Supplementary File S2. Note that delta sequences are Ty1 retrotransposon long terminal repeats, sigma sequences are Ty3 retrotransposon long terminal repeats, tau sequences are Ty4 retrotransposon long terminal repeats, Ty1, Ty3, and Ty4 are retrotransposons; ARS sequences are Autonomously Replicating Sequences.

| **Deletions** | | | | | | |
| --- | --- | --- | --- | --- | --- | --- |
| **Abbreviation** | **Type** | **Location** | **Loci** | **Clonal cluster affected** | **Frequency** | **Chromosomal copies affected** |
| -on I | terminal deletion | I: 138620-230218 | from YALCdelta2 to right telomere | Mixed origin d: APN to DBL1 | 6 of 7 isolates | 2 |
| -on V | terminal deletion | V: 435946-576874 | from YERCdelta16 to right telomere (same loci as +on V) | Mixed origin d: APM to UDeb-BY0022 | 1 of 9 isolates | 1 |
|  |  |  |  | Mixed origin e | 1 of 26 (and 2 unknown) |  |
| -on VII.a | terminal deletion | VII: 1-74054 | from left telomere to intergenic region | Mixed origin b | 15 of 16 isolates | 1 |
| -on VII.b | interstitial deletion | VII: 319113-405226 | from YGLCdelta5 to YGLCsigma3 | Mixed origin e | 25 of 26 (and 2 unknown) | 1 |
| -on VII.c | interstitial deletion | VII: 712545-779482 | from YGRWTy3-1 to YGRCdelta25 | Mixed origin e | 1 of 26 (and 2 unknown) | 1 |
| -on XI | terminal deletion | XI: 578750-666816 | from intergenic region, recombination hotspot in Pan et al. (2011), to right telomere | Mixed origin c | 1 of 26 isolates | 1 |
| -on XIV.a | interstitial deletion | XIV: 561139-632492 | from ARS1422 to YNRCtau3 | Mixed origin d: APN to DBL1 | 3 of 7 isolates | 1 |
| -on XIV.b | terminal deletion | XIV: 632493-784334 | from YNRCtau3 to right telomere | Mixed origin d: APN to DBL1 | 3 of 7 isolates | 2 |
| -on XV | terminal deletion | XV: 577195-1091291 | from *EFT1* to right telomere | Mixed origin b | 1 of 16 isolates | 1 |
| -on M | almost complete loss of mtDNA | Mitochondrial DNA | Mitochondrial DNA | aff. Mixed origin | 1 of 3 isolates | not applicable |

| **Duplications** | | | | | | |
| --- | --- | --- | --- | --- | --- | --- |
| **Abbreviation** | **Type** | **Location** | **Loci** | **Clonal cluster affected** | **Frequency** | **Chromosomal copies affected** |
| +on III | terminal duplication | III: 200961-316620 | from MAT-locus and YCR041W to right telomere | Mixed origin e | 26 of 26 isolates (and 2 unknown) | 1 |
| +on IV | terminal duplication | IV: 1245186-1531933 | from *EFT2* to right telomere | Mixed origin b | 1 of 16 isolates | 1 |
| +on V | terminal duplication | V: 435946-576874 | from YERCdelta16 to right telomere (same loci as -on V) | Mixed origin d: NRRLYB-3916 to CBS2919 | 1 of 3 isolates | 1 |
| +on VII.a | terminal duplication | VII: 1 to 405228 | from left telomere to YGLCsigma3 | Mixed origin d: NRRLYB-3916 to CBS2919 | 3 of 3 isolates | 1 |
| +on VII.b | terminal duplication | VII: 712545-1090940 | YGRWsigma5 recombination hotspot in Pan et al. (2011) to right telomere | Mixed origin d: APM to UDeb-BY0022 | 1 of 9 isolates | 1 |
| +on IX | terminal duplication | IX: 325732 to 439888 | intergenic region upstream to YNCI0009W tRNA gene, recombination hotspot in Pan et al. (2011) to right telomere | Mixed origin d: NRRLYB-3916 to CBS2919 | 3 of 3 isolates | 1 |
| +on X | interstitial duplication | X: 204254-354843 | from ARS1008 to YJLCdelta4/ YJLCdelta5 | Mixed origin d: NRRLYB-3916 to CBS2919 | 3 of 3 isolates | 1 |
| +on XI.a | terminal duplication | XI: 1-314193 | from left telomere to YKLWdelta7 | Mixed origin c | 1 of 26 isolates | 1 |
| +on XI.b | terminal duplication | XI: 517451-666816 | from ARS1116, recombination hotspot in Pan et al. (2011) to right telomere | Mixed origin a | 3 of 3 isolates | 2 |
| +on XII | terminal duplication | XII: 1- approx. 97612 | from left telomere to intergenic region upstream to POM33, recombination hotspot in Pan et al. (2011) | Mixed origin e | 1 of 26 (and 2 unknown) | 1 |
| +on XIII | terminal duplication | XIII: 1-209069 | from left telomere to intergenic region, near recombination hotspot in Pan et al. (2011) | Mixed origin f | 1 of 16 | 1 |
| +on XVI.a | terminal duplication | XVI: 520521-948066 | from intergenic region, recombination hotspot in Pan et al. (2011) to right telomere | Mixed origin e | 1 of 26 (and 2 uncertain) | 1 |
| +on XVI.b | terminal duplication | XVI: 769926-948066 | from YPRWsigma2 to right telomere | Mixed origin e | 1 of 26 (and 2 uncertain) | 1 |
