## Supplementary File S5 for "Commercial *Saccharomyces cerevisiae* baker’s yeasts: strain redundancy, genome plasticity, and colonization of the sourdough environment and the human body"

**Supplementary File S5. Companies and brands of yeasts in the baking industry.**

Companies, subsidiaries, and brands were identified for sequenced yeasts in the Mixed origin group wherever possible. For each subclade, the number of sequenced strains and isolates is shown (y axis) with color coding in the below graph.

|  | Unknown | Company 22 | Company 12 | Company 10 | #REF! | Company 1 | Fleischmann & Co | Company 4 | Company 8 | Company 11 | Company 17 | Company 18 | Company 14 | Company 3 |
| --- | --- | --- | --- | --- | --- | --- | --- | --- | --- | --- | --- | --- | --- | --- |
| Mixed origin a | 3 | 1 | 0 | 0 | 1 | 0 | 0 | 0 | 0 | 0 | 0 | 0 | 0 | 0 |
| Mixed origin b | 2 | 0 | 3 | 2 |  | 0 | 0 | 0 | 0 | 0 | 0 | 0 | 0 | 1 |
| Mixed origin c | 5 | 0 | 2 | 0 |  | 1 | 0 | 0 | 0 | 0 | 0 | 0 | 0 | 0 |
| Mixed origin d | 16 | 0 | 2 | 0 |  | 2 | 2 | 1 | 2 | 0 | 0 | 0 | 0 | 0 |
| Mixed origin e | 19 | 0 | 1 | 0 |  | 1 | 0 | 0 | 0 | 1 | 2 | 2 | 1 | 0 |
| Mixed origin f | 4 | 0 | 0 | 0 |  | 0 | 0 | 0 | 0 | 0 | 0 | 0 | 0 | 2 |

**Number of yeast genomes associated with each company in the various subclades.**

The companies and subsidiaries associated with the yeast genomes are listed below, brand names are included in the description.

| **Company** | **Description** |
| --- | --- |
| Company 1a | Globally active, UK-based food processing and retailing company |
| Company 1b | Multinational bakery ingredient, yeast, and oenology company, a division of Company 1a |
| Company 1c | American subsidiary of Company 1a, licences the name “Fleischmann’s Yeast” from Company 1b, manufactures in Canada |
| Company 2 | Germany based multinational family-owned discount supermarket chain |
| Company 3a | Globally active, Canadian yeast production company founded in 1853, owns multiple subsidiaries and brands, *e.g.* for brewing yeast products, or for wine yeasts |
| Company 3b | South African company specialized in yeast and oenology products founded in 1923, acquired by Company 3a in 2006 |
| Company 3c | German food production company founded in 1862, formerly family-owned, became a brand in the German subsidiary of Company 3a; the brand is also used for yeasts produced in Austria and marketed by Company 2 |
| Company 3d | Subsidiary of Company 3 based in UK, formerly independent company founded in 1932. Owns a baking yeast brand and production facilities in the UK formerly owned by Company 1a, but sold due to competition concerns to Company 3a |
| Company 4 | Food manufacturer and brewery from Japan |
| Company 5 | Multinational retail group headquartered in France |
| Company 6 | Multinational pharmaceutical company founded in 1953 in France that obtained the patent for the original *S. 'boulardii'* strain CNCM I-745 from Henri Boulard in the 1950s. Has production plants in France and Morocco; produces probiotic yeast under various brand names for the global market. |
| Company 7 | Austrian animal nutrition company founded in 1983 |
| Company 8 | German family-owned multinational food production company founded in 1891, produces primarily in Germany |
| Company 9 | European chain of retail stores specialized in cosmetics, healthcare items, household products and health food headquartered in Germany, founded in 1973, sells "bio" yeasts under its own bio-food brand. |
| Company 10 | Austrian family-owned independent foodstuff production company founded in 1915, producing in Austria |
| Fleischmann | Company founded in the US by European immigrants from Austria-Hungary in the 1860s, started production of shelf stable granular yeast in the 1940s. Company ceased operations, brand currently owned by Associated British Foods. The original company promoted the consumption of live yeast as a health supplement in the Yeast for Health campaign from 1917 to 1924, and later continued promoting new strains as vitamin-rich supplements. Applied irradiation to yeasts to enhance vitamin D levels as early as 1929 |
| Company 11 | Food manufacturing company from Australia |
| Company 12a | Globally active, French yeast production company founded in 1853, owns multiple subsidiaries and brands |
| Company 12b | Belgian yeast producer founded in 1949, now subsidiary of Company 12a |
| Company 12c | Beer, wine, and spirit yeast business unit of Company 12a, also based in France, with multiple production plants in Europe, owns multiple brands, *e.g.* a brand names used for brewing yeasts |
| Company 12d | Hungarian subsidiary of Company 12a. The Hungarian subsidiary was an independent company founded in 1876, specialized in baking yeast manufacturing, its name is now a brand name |
| Company 12e | Italian subsidiary of Company 12a, manufacturing a locally branded yeast |
| Company 12f | US subsidiary of Company 12a, specialized in baking yeast production, with multiple production sites in the US |
| Company 13 | German global discount supermarket chain |
| Northwestern Yeast Company | Yeast manufacturer company founded in 1893 in Chicago, IL, USA, famous for two brands, Magic Yeast and Yeast Foam. The company successfully marketed yeasts as health products with a campaign started in the 1920s. Company ceased to exist. |
| Company 14 | Nutritional supplement manufacturing company from Spain |
| Company 15 | Globally active food manufacturing company founded in 1973 in Turkey/Türkie, part of a larger group |
| Company 16 | Central European discount supermarket chain based in Germany |
| Company 17 | German foodstuff production company founded in 1920, producing in Germany |
| Company 18 | Hungarian foodstuff production company, manufactures yeasts under the brand name of a well-known former family brand founded in 1925 |
| Company 19 | Multinational franchise founded in the Netherlands, managing food retail stores |
| Company 20 | Austrian fresh pastry company |
| Company 21 | British multinational groceries and general merchandise retailer operating worldwide |
| Company 22 | German company manufacturing baking products, founded in 1975 by a merger of two independent companies, predecessor companies were originally founded in 1793 |

**Companies and subsidiaries associated with the yeast genomes in this study. Companies active currently are listed only with identifiers.**
