## Supplementary File S6 for "Commercial *Saccharomyces cerevisiae* baker’s yeasts: strain redundancy, genome plasticity, and colonization of the sourdough environment and the human body"

**Supplementary File S6. Sporulation and spore viability data.**

Sporulation and spore viability for commercial baker’s yeasts isolated in this study in the Mixed origin clade. Spore viability was tested on isolates showing at least 20% sporulation, below that, no spore viability data is available as spore isolation in high numbers from low-sporulating isolates is technically unfeasible.

| **Group** | **Isolate** | **Sporulation %** | **Combined spore viability %** | **Number of tetrads with 4, 3, 2, 1, or 0 viable spores** | | | | |
| --- | --- | --- | --- | --- | --- | --- | --- | --- |
|  |  |  |  | **4 alive** | **3 alive** | **2 alive** | **1 alive** | **0 alive** |
| **Mixed origin ‘a’** | **UDeb-BY0026** | 1 | n.d. | n.d. | n.d. | n.d. | n.d. | n.d. |
| **Mixed origin ‘b’** | **UDeb-BY0001** | 35 | 55 | 0 | 4 | 4 | 2 | 0 |
|  | **UDeb-BY0002** | 54 | 55 | 2 | 2 | 3 | 2 | 1 |
|  | **UDeb-BY0007** | 77 | 27.5 | 0 | 1 | 1 | 6 | 2 |
|  | **UDeb-BY0008** | 38 | 52.5 | 1 | 4 | 2 | 1 | 2 |
|  | **UDeb-BY0017** | 80 | 57.5 | 2 | 1 | 5 | 2 | 0 |
|  | **UDeb-BY0018** | 58 | 70 | 2 | 5 | 2 | 1 | 0 |
| **Mixed origin ‘d’** | **UDeb-BY0022** | 5 | n.d. | n.d. | n.d. | n.d. | n.d. | n.d. |
|  | **UDeb-BY0023** | 4 | n.d. | n.d. | n.d. | n.d. | n.d. | n.d. |
| **Mixed origin ‘e’** | **UDeb-BY0005** | 44 | 60 | 1 | 4 | 3 | 2 | 0 |
|  | **UDeb-BY0006** | 25 | 57.5 | 1 | 2 | 6 | 1 | 0 |
|  | **UDeb-BY0009** | 11 | n.d. | n.d. | n.d. | n.d. | n.d. | n.d. |
|  | **UDeb-BY0010** | 19 | n.d. | n.d. | n.d. | n.d. | n.d. | n.d. |
|  | **UDeb-BY0011** | 11 | n.d. | n.d. | n.d. | n.d. | n.d. | n.d. |
|  | **UDeb-BY0012** | 31 | 87.5 | 7 | 1 | 2 | 0 | 0 |
|  | **UDeb-BY0013** | 6 | n.d. | n.d. | n.d. | n.d. | n.d. | n.d. |
|  | **UDeb-BY0014** | 8 | n.d. | n.d. | n.d. | n.d. | n.d. | n.d. |
|  | **UDeb-BY0015** | 22 | 50 | 0 | 2 | 5 | 2 | 0 |
|  | **UDeb-BY0016** | 6 | n.d. | n.d. | n.d. | n.d. | n.d. | n.d. |
|  | **UDeb-BY0019** | 4 | n.d. | n.d. | n.d. | n.d. | n.d. | n.d. |
|  | **UDeb-BY0024** | 9 | n.d. | n.d. | n.d. | n.d. | n.d. | n.d. |
|  | **UDeb-BY0025** | 15 | n.d. | n.d. | n.d. | n.d. | n.d. | n.d. |
|  | **UDeb-BY0027** | 18 | n.d. | n.d. | n.d. | n.d. | n.d. | n.d. |
|  | **UDeb-BY0028** | 15 | n.d. | n.d. | n.d. | n.d. | n.d. | n.d. |
| **Mixed origin ‘f’** | **UDeb-BY0003** | 7 | n.d. | n.d. | n.d. | n.d. | n.d. | n.d. |
|  | **UDeb-BY0004** | 5 | n.d. | n.d. | n.d. | n.d. | n.d. | n.d. |
|  | **UDeb-BY0020** | 6 | n.d. | n.d. | n.d. | n.d. | n.d. | n.d. |
|  | **UDeb-BY0021** | 0 | n.d. | n.d. | n.d. | n.d. | n.d. | n.d. |
